# Model-based Standardization of Correlation Coefficients Improves Multi-Omic Clustering and Biological Signal Discovery

**DOI:** 10.1101/2025.11.17.688875

**Authors:** Max Robinson, Heeju Noh, Lance Pflieger, Noa Rappaport

**Affiliations:** Institute for Systems Biology, Seattle, WA 98109; Phenome Health, Seattle WA 98109

## Abstract

Multi-omic data pose a particular challenge for Weighted Correlation Network Analysis (WCNA or WGCNA) due to (platform- or) batch-specific characteristics, such as resolution, accuracy, dynamic range, and sources of spurious variation. When unaccounted for, these differences can result in a bias toward single-batch clusters as well as greater sensitivity to “noisier” batches during clustering. Here we propose mitigating these effects using null models fitted separately to the bulk of analyte-analyte correlations within each batch and across each pair of batches. We then map the batch-specific null models to a standard null model, removing batch-dependent distributional differences. This approach is compatible with any correlation-based clustering approach. Since the null model represents information not captured in individual pairwise correlations, we show how to incorporate this additional information into both distance-based clustering and WCNA. For distance-based clustering, we increase distances corresponding to correlations consistent with the null model. For WCNA, we provide a new soft threshold (adjacency) function based on the likelihood of a correlation under the null model. The resulting network can be easily incorporated into the WCNA workflow. These methods are implemented in R package `standardcor`, and we illustrate the package on simulated data as well as an existing multi-omic dataset.

## Introduction

Biological systems are now measured by multiple high-throughput technologies (‘omics), potentially providing counts of gene transcripts as well as amounts of proteins, metabolites, lipids, and other molecular data from the same individual or biological sample(Chen et al. 2012; Gibbs et al. 2014; Price et al. 2017; Shen et al. 2024). A multi-omic dataset combines data about interrelated aspects of a biological system–regulation of transcription, translation, and localization, as well as enzymatic activities–with the goal of identifying subsystems associated with particular system behaviors(Gibbs et al. 2014; Price et al. 2017; Earls et al. 2019).

This goal is often approached through correlation-based clustering, a geometric approach to grouping subsystem components through pairwise correlation coefficients. In a systems biology approach, each analyte is imagined as a point in space, located by its measurements across samples, and correlation is used to capture relationships between component pairs. Similarity between two analytes *X* and *Y*, measured across the same set of (e.g. biological) samples, can be estimated as a correlation coefficient ρ(*X, Y*) which has a simple geometric interpretation as the cosine of an angle (see Supplement S1),

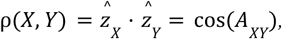

where 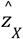 and 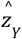 are the unit vectors in the direction of *X* and *Y*, respectively. While distances between analyte abundances capture relationships on an absolute scale (offsets), correlations capture relationships on a relative scale (slopes). These relationships are transformed into geometric patterns by converting the correlations to non-negative values, either distances (smaller-is-closer) or similarities (larger-is-closer), and these representations support different clustering criteria and methods.

Ward(Ward 1963) proposed the distance-based hierarchical clustering approach (R function `hclust()(The R Project for Statistical Computing)`) in which analytes are progressively grouped into larger sets while minimizing an objective function; in Ward’s minimum variance method, the two sets which least increase the total variance (the within-cluster sum of squared distances from the cluster centroids) are merged at each step until only one set remains. The resulting tree can then be “cut” at a chosen level to identify a partitioning set of clusters. Another well-known distance-based method, *k*-means clustering(Ball and Hall 1967; Lloyd 1982), partitions the analytes into a specified number of clusters *k* to minimize the total variance; it does not construct an entire hierarchy and is typically faster than hierarchical clustering on the same data. Both hierarchical and *k*-means clustering algorithms are heuristic solutions of an NP-hard problem(Megiddo and Supowit 1984) and may return suboptimal results.

WCNA(Horvath 2011) is a systems biology approach to analyzing ‘omics data(Zhang and Horvath 2005) which uses an alternative, similarity-based approach to produce clusters. Beginning from pairwise correlations, WCNA constructs a weighted network with edge weights (adjacencies) between 0 and 1. While an unweighted network could be constructed by thresholding, including every edge with an adjacency above a threshold *t* ≥ 0, a weighted network replaces discontinuous behavior at a single threshold with “soft thresholding”, the opportunity for a smooth transition in behavior at all adjacencies(Horvath 2011). In a network, a cluster corresponds to a region of high adjacency among interconnected nodes. After constructing an initial adjacency network, WCNA uses the Topological Overlap Measure (TOM)(Li and Horvath 2007) to find the density of interconnection surrounding each edge and converts it to a distance to identify clusters via hierarchical clustering and a dynamic cutting algorithm(Langfelder et al. 2008). Clustering in WCNA is also heuristic.

General clustering algorithms like Ward and WCNA can be applied to multi-omic data with the simple “concatenated clustering” strategy(Zhang et al. 2022), in which the ‘omics datasets are concatenated and clustered as a single dataset. However, each ‘omics platform (or batch) may have unique experimental sources of variation, and pairwise correlations within a batch or across the same pair of batches will exhibit a characteristic mean and variance(Zhang et al. 2022). These distributional differences favor construction of clusters enriched for analytes from the same batch, and should be mitigated to yield best results.

Methods developed specifically to address the challenges of multi-omic clustering (reviewed in (Zhang et al. 2022)) have not yet taken the form of a separate process prior to concatenated clustering. For example, moCluster(Meng et al. 2016) attempts to reduce the impact of distributional differences in the correlation matrix with a weighting strategy during construction of a correlation network. Multi-Omics Factor Analysis (MOFA)(Argelaguet et al. 2018) uses dimensionality reduction to identify covarying sets of analytes as latent factors, and the weights in the construction of the latent factors provide an indirect means of reweighting the contributions of different pairwise correlations. While MOFA is not explicitly a clustering method, the latent factors provide a lower-dimensional space in which distance-based clustering can be performed.

Here we present a general preparatory method enabling use of the general concatenated clustering strategy. We propose fitting null models to the observed correlation coefficients within each batch and between each pair of batches to explicitly account for batch biases. These null models can then be used to standardize correlations across batches, enabling concatenated clustering beginning from the standardized correlations. However, the null models can also assist in separating significant from insignificant correlations during clustering.

### Significance of correlation coefficients

While significance tests for correlation coefficients are routinely used, it is not entirely clear what distribution to use for a null model of correlation coefficients. Consider an experiment in which each of *N* biological samples is represented by *n* measurements which are Normally distributed, and our null hypothesis is that they are uncorrelated across samples. We compute the pairwise correlations ρ among these *N* samples, and under the null hypothesis (*E*[ρ] = 0), the standard paired sample *t*-test statistic, 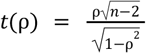, can be used; *t*(ρ) follows Student’s *t* distribution with *n* − 2 degrees of freedom(Kutner et al. 2004). When *E*[ρ] ≠ 0 this test does not apply, but we can test the more general null hypothesis ρ = *E*[ρ] using the Fisher transformation 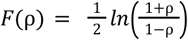 to obtain the statistic 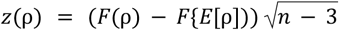 is known to be approximately Normally distributed(Fisher 1915). Both of these tests provide distributions for nonlinear functions of ρ, neither of which suggests the form of an appropriate null distribution for the correlation coefficients ρ themselves. Both the t-test and the Fisher transformation depend on *n*, the dimensionality of the data being assessed, however the standard translation of a correlation to squared Euclidean distance, *D*^2^ (ρ) = 1 − ρ, does not. This suggests that the distances currently used for correlation-based clustering do not properly account for the geometric properties of correlation coefficients in *n* > 2 dimensions.

In an early approach to analyzing gene expression data, Ji et al.(Ji et al. 2005) suggested using a mixture of two Beta distributions to model observed correlation coefficients, citing the flexibility of the Beta distribution for representing a distribution on a fixed interval. In their model, one Beta distribution served as a null model and the other served as a model of non-null correlations. Below, we will argue that spurious correlations of each type (i.e. between each specific pair of batches) should follow a Beta distribution, and that type-specific Beta null models provide a means to standardize the different types of correlations to a reference null model.

With this theoretical motivation and plan, we developed R package `standardcor(standardcor)` which supports three new capabilities. First, it provides a methodology for standardizing correlations computed within and across multiple data sources (‘omics), enabling the use of concatenated clustering and standard clustering approaches on multiomic data. Second, it provides a new approach to convert correlation coefficients to network adjacencies in a way that accounts for statistical significance, and we show that this approach provides improved clustering when incorporated into WCNA. Third, we provide a “distance penalty” approach to produce a distance sensitive to the statistics of correlation coefficients for distance-based clustering. To validate these three methods, we measure the improvement to concatenated clustering using both distance-based and similarity-based methods on synthetic multiomic data. Finally, we demonstrate the usefulness of this approach by identifying improved associations of clusters with age on the biologically relevant, multi-omic Arivale dataset.

## Results

We have developed R package `standardcor(standardcor)` to support correlation-based clustering of single-omic and multi-omic datasets. Its main innovation is the use of separate null models of correlation coefficients for analytes within each batch (‘omics platform) and for analytes from one batch versus analytes from a different batch. These null models provide a new method for transforming correlation coefficients into adjacencies (or distances), and for multiomic datasets, the means to standardize the statistical significance of correlation values across ‘omics platforms (batches) and therefore concatenated correlation-based clustering of multiomic data using algorithms developed for unstructured datasets.

### Fast, sparse Spearman correlations

The first step is to compute pairwise correlation coefficients *Z* = {*z*_*ij*_} between all analytes measured on a common set of samples. We find an especially time- and space-efficient Spearman correlation function (`SparseSpearmanCor2()(were made available under the BSD 2-c…)`) useful for transcriptomics and multi-omic datasets, either of which may contain tens of thousands of analytes. Although we use Spearman correlation here, the methodology presented here is appropriate for a variety of correlation coefficients, including biweight midcorrelation (WGCNA::bicor()(Hardin et al. 2007)). For comparison, we include results for `bicor()` on an experimental dataset (see Arivale below).

### A null model for correlation coefficients

While Ji(Ji et al. 2005) recommended Beta distributions for their flexibility, we argue that spurious correlation coefficients should follow a Beta distribution, providing a stronger justification. The argument stems from the formula for Pearson’s sample correlation coefficient. Suppose *X* and *Y* are two *n*-dimensional points (or a set of *n* pairs, (*x*_*i*_, *y*_*i*_)). Using 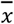 and 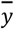 as the means of *X* and *Y* and *s*_*x*_ and *s*_*y*_ as their sample standard deviations, the sample correlation coefficient can be written as

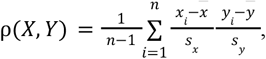

and the contribution of a specific pair *i* is 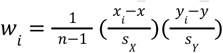. Separating positive and negative terms,

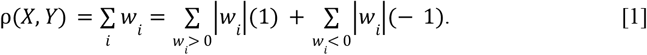

The correlation coefficient is therefore the result of a *weighted* vote between two alternatives, correlation (+1) and anticorrelation (−1). Since *ab* ≠ 0 and *a*/*b* have the same sign, pair *i* “votes” for correlation when the slope of the line through 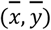 and (*x*_*i*_, *y*_*i*_) is positive 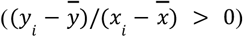, or for anticorrelation when it is negative. Setting the magnitude of the weights equal for the moment (|*w*_*i*_| = 1/*n*) and letting k be the number of positive votes, this sum becomes the Quadrant Count Ratio (QCR)(Kader and Franklin 2008):

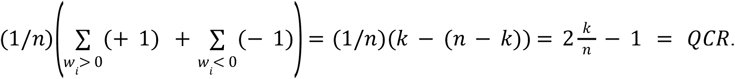

The QCR is the result of an unweighted voting process and thus a function of a Binomial random variable *k* ~ *B*(*n, f*), with each success a +1 vote, each failure a −1 vote, and probability of success 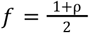. The sampling distribution for estimates of *f* is known to be a Beta distribution *f* ~ β(ν, ω), with *v* = *k* successes and ω = *n* − (*k* − 1) failures(Hastings et al. 2000). Since ν and ω depend only on the vote totals and need not be integers, the weighted sum (Eq. 1) follows the same logic to a Beta distribution for the probability of success, yielding the form of our null model:

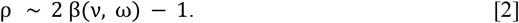

The parameters (ν, ω) are related to *n* and *f* by the equations

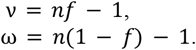

In the case addressed by Student’s *t*-test 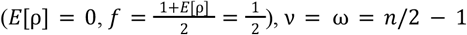 and the null distribution depends only on *n*:

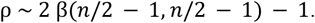

This vote counting argument applies equally to Spearman correlation, `bicor()`, and any other measure of correlation with the same general form as Pearson’s, but note that it does not necessarily apply to association measures sometimes used in WCNA, e.g. Mutual Information or Hellinger’s(Zhang and Wong 2022), which have a different form.

Data simulated with *n* independent Gaussian dimensions support the correctness of this Beta null model (Fig. 1, see also (Relationship between random n-vectors…)). Upon further investigation (Supplement S2), when *E*[ρ] = 0 Student’s *t*-test and the test based on this Beta distribution are identical up to numerical error. When *E*[ρ] ≠ 0, Fisher’s transformation *z* = *arctanh*(ρ) (Fisher 1915) provides an approximately Gaussian *z*-statistic; our simulations show that the Beta model appears to be exact (Supplement S2) for Gaussian marginal distributions and that Fisher’s transformation appears to converge to the Beta model as *n* increases. We have also shown (Supplement S3) that the marginal distributions need not be Gaussian for the null distribution to be our Beta model, but that some marginal distributions are inconsistent with the Beta null model (the ones we have found have infinite variance). In summary, our simulations show this Beta null model applies for a broad class of marginal distributions and provides an accurate exact test even when *E*[ρ] ≠ 0.

**Figure 1.**
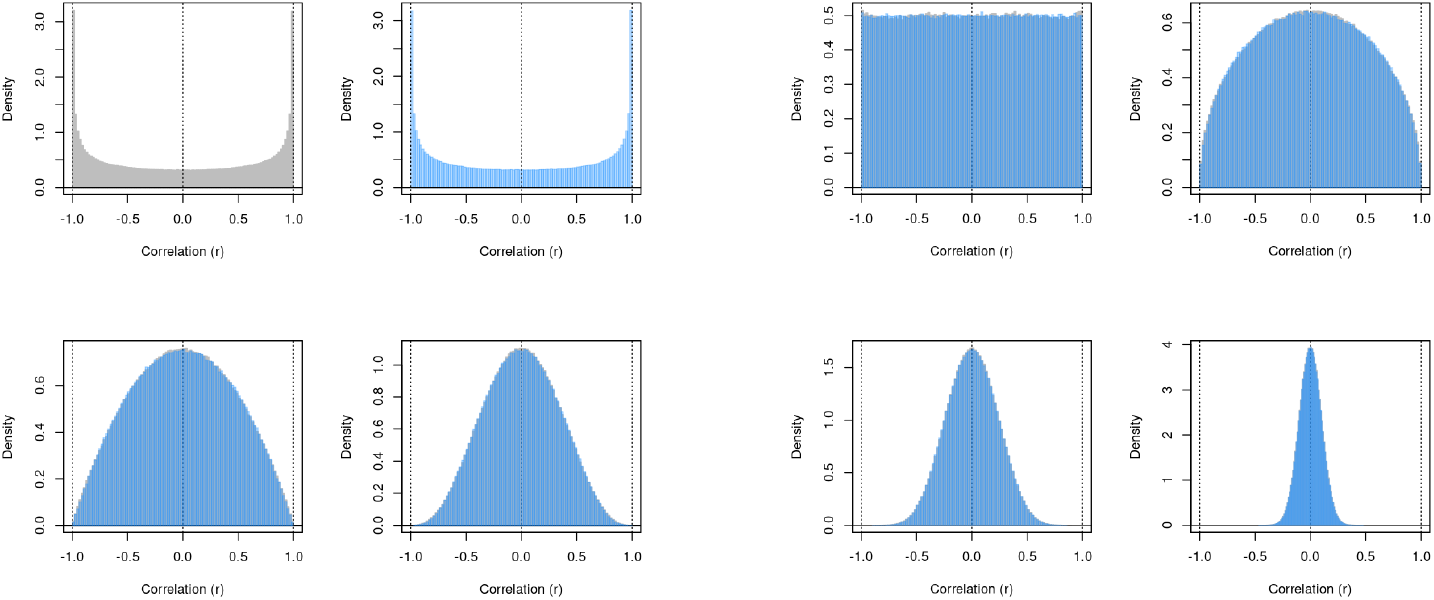
Distribution of correlations of random n-vectors and the Beta null model. Upper left pair: A histogram of simulated 3-vector correlations (gray) and a probability density plot of β(*v, v*), *v* = *n*/2 − 1 (blue), illustrate the correctness of the Beta null model for *n* = 3. Correctness is illustrated with overlaid histogram and density for *n* = 4 and *n* = 5 (upper right pair), *n* = 6 and *n* = 10 (lower left pair); and *n* = 20 and *n* = 100 (lower right pair). For additional analysis, see Supplement S3.

### Practical estimation of null models

We observe that the distribution of pairwise correlation coefficients may differ according to the measurement technologies (or batches thereof) being compared. For *K* ‘omics sets we therefore fit a null model of the form Beta(*v, w*) to the correlation coefficients of each of the *K*(*K* + 1)/2 pairs of batches, resulting in *K*(*K* + 1) fitted parameters. We assume each subset may include a minority of outliers, and therefore expect a parameter estimation method resistant to the presence of outliers will be required to find appropriate parameters.

The standard methods for estimating the parameters of the Beta distribution are the method of moments and maximum likelihood, however neither of these methods is outlier resistant(Hastings et al. 2000). Package `standardcor` includes a novel heuristic estimation procedure, `estimateShape()`, which we observe to find better estimates in the presence of outliers, and provides additional parameters to assist the user in refining its estimates. For multiomic data we provide a function (`multiOmicModel()`) which automates fitting all required null models for a list of batches.

### Standardizing correlations across data sources

While the fitted null models described above are sufficient for producing adjacencies using the facilities in the package, we find it convenient to convert the entire correlation matrix *Z* to a matrix of “standardized correlations” sharing a single reference null model *B*(*v*_0_, *w*_0_). This standardized correlation matrix *Z*_*c*_ = {*S*(*z*_*ij*_)}, has each standardized correlation *S*(*z*_*ij*_) at the same quantile of the reference null distribution as the quantile *z*_*ij*_ represents in its corresponding fitted null distribution *B* (*v*_*ij*_, *w*_*ij*_):

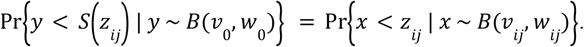

*S*(*z*_*ij*_) is computed by `standardizeFromModel()` using the R functions `qbeta()` and `pbeta()`. Standardized correlations are intended to reduce the bias toward single-batch clusters in a multi-omic dataset; we evaluated their ability to meet this goal, as well as their impact on cluster quality, for both distance-based and similarity-based clustering methods (see below).

### Enhanced Distance

`standardcor` currently uses the null model in non-equivalent ways to support distance-based and similarity-based clustering methods^1^. For distance-based clustering, we use the null model probability density function (unsigned interpretation of correlations) or cumulative density function (signed interpretation) to extend the standard squared Euclidean distance associated with a correlation coefficient. Let 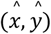 denote a pair of vectors on the (*n* − 1)-dimensional unit sphere. The probability

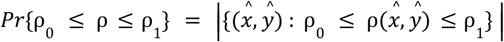

is the size of the set of vector pairs which yield a correlation coefficient in the given range. As *n* increases, a larger fraction of the pairs of vectors on (*n* − 1)-dimensional unit sphere are nearly orthogonal (Fig. 1), and the significance of a fixed correlation coefficient ρ ≠ 0 will increase. In contrast, the squared Euclidean distance (1 − ρ signed interpretation of correlation, or 1 − |ρ| unsigned) corresponding to ρ does not change with *n*. In order to reflect the change in significance with *n*, we propose adding a distance penalty based on the probability under the null model (*Pr*{*x* ≥ ρ} signed, or the probability density *Pr*{|ρ|} unsigned). We refer to this as Enhanced Distance, and provide the function `standarcor::betaDistance()` to compute it.

### Adjacencies from the null model

WCNA is similarity-based rather than distance-based and constructs a weighted correlation network using similarity edge weights (adjacencies). The adjacencies serve as a soft thresholding mechanism as follows. Given adjacencies *a*_*ij*_ = *A*(ρ_*ij*_) between 0 and 1, a hard threshold *t* would exclude edges *e*_*ij*_ from an unweighted correlation network when *a*_*ij*_ < *t*. A weighted correlation network includes all edges *e*_*ij*_ with *a*_*ij*_ > 0, annotating each edge with its adjacency. The adjacency can be interpreted as the probability of including *e*_*ij*_ in an unweighted network for an unknown hard threshold *t* > 0, allowing edges to have different degrees of influence on clustering (soft thresholding).

We use a null model of correlation coefficients to build an adjacency function *A*(ρ) as follows. Given an observed set of correlation coefficients, we assume that the majority come from the null model (ρ ~ 2*B*(ν, ω) − 1) but that an unknown subset do not (the “outliers”). Any hard threshold *t* ≥ 0 with correlations |ρ| > *t* considered outliers would exclude some number *FN*(*t*) of outliers (false negatives) and include some number *FP*(*t*) of null correlations (false positives). We define the adjacency of correlation ρ as the fraction of outliers among the correlations *misclassified* at threshold *t* = ρ:

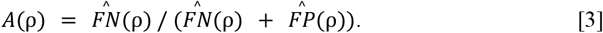

We compute *A*(ρ) from the observed correlation coefficients with the same null model ρ ~ 2*B*(ν, ω) − 1 by first estimating the number of null *N* of null correlations that were observed. Making the assumption that the middle half of the observed correlations are the middle half of the null distribution yields an estimate of *N* and an estimate of *FP*(ρ):

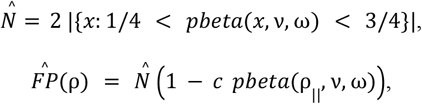

where *c* = 2 and ρ_||_ = |ρ| if the correlations are interpreted as association (an “unsigned” network), or *c* = 1 and ρ_||_ = ρ (a signed network). While this assumption may overestimate the number of null correlations, when some outliers are present and the null distribution is correct it is unlikely to be far wrong, because we expect more null correlations than outliers and the most easily distinguishable outliers lie at the extremes of the null distribution. Counting the number *U*(ρ) of observed correlations exceeding threshold ρ_||_ then provides a corresponding estimate of *FN*(ρ):

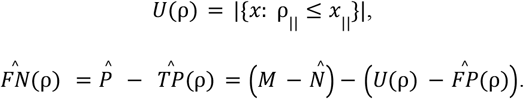

Package `standardcor` separates conversion of correlations into adjacency into two-steps, first tabulating the adjacency function (`nullModelAdjacencyTable()`) and then finding the adjacency corresponding to each correlation from the table by interpolation (`interpolatedAdjacency()`). The primary reason for this design was to simplify visualizing the adjacency function.

### Testing Strategy

We tested whether converting correlations to standard squared Euclidean distance or to Enhanced Distances provide better clustering using simulated abundance profiles (see below). Each test dataset was a matrix representing 100 samples (rows) and 800 analytes (columns), 400 in ‘omics batch “A” and 400 in ‘omics batch “B”, to represent a simple multi-omic experiment. The analytes were randomly partitioned into *n* = 5, 10, or 20 “true” clusters (at least 20 analytes each), and pairwise correlations were designed at two difficulty levels (high and low SNR), using a method based on Cholesky Decomposition(Kaiser and Dickman 1962) to generate correlated data(see Methods). At both high and low SNR, design goals were that correlations between analytes in the same cluster (signal) were higher than those between analytes in different clusters (noise), and correlations between analytes from the same batch were higher than between analytes from different batches (see Methods); all within-cluster correlations were positive. Within-cluster, within-batch correlations were 0. 2 ≤ ρ ≤ 0. 4(high SNR) or 0. 1 ≤ ρ ≤ 0. 2 (low SNR). For each number of clusters and difficulty level we generated 100 replicates, a total of 6 scenarios, 600 test datasets. Each of the 7 clustering methods was run to produce the same number of predicted clusters as true clusters using an unsigned interpretation of correlations.

### Enhanced Distance: Example

For clarity, we illustrate a comparison of two distance-based clustering methods, both versions of Ward’s minimum variance hierarchical clustering but using different correlation-based distances, on a single test dataset at high SNR and *n* = 20 clusters (Fig. 2). Method “HCLUST” used squared Euclidean distance and provided weak but consistent contrast between within-group (dark, smaller) and between-group (light, larger) pairs (Fig. 2a, left). Method “HCLUST-ES” used Enhanced distance (computed from Standardized correlations) and provided both stronger contrast (Fig. 2a, right) and increased silhouette scores for most analytes (Fig. 2b, right), indications of greater cluster separation. Enhanced distance resulted in higher clustering accuracy in this example (0.86 vs 0.83). For reliable results, each test was run on 100 replicate datasets.

**Figure 2.**
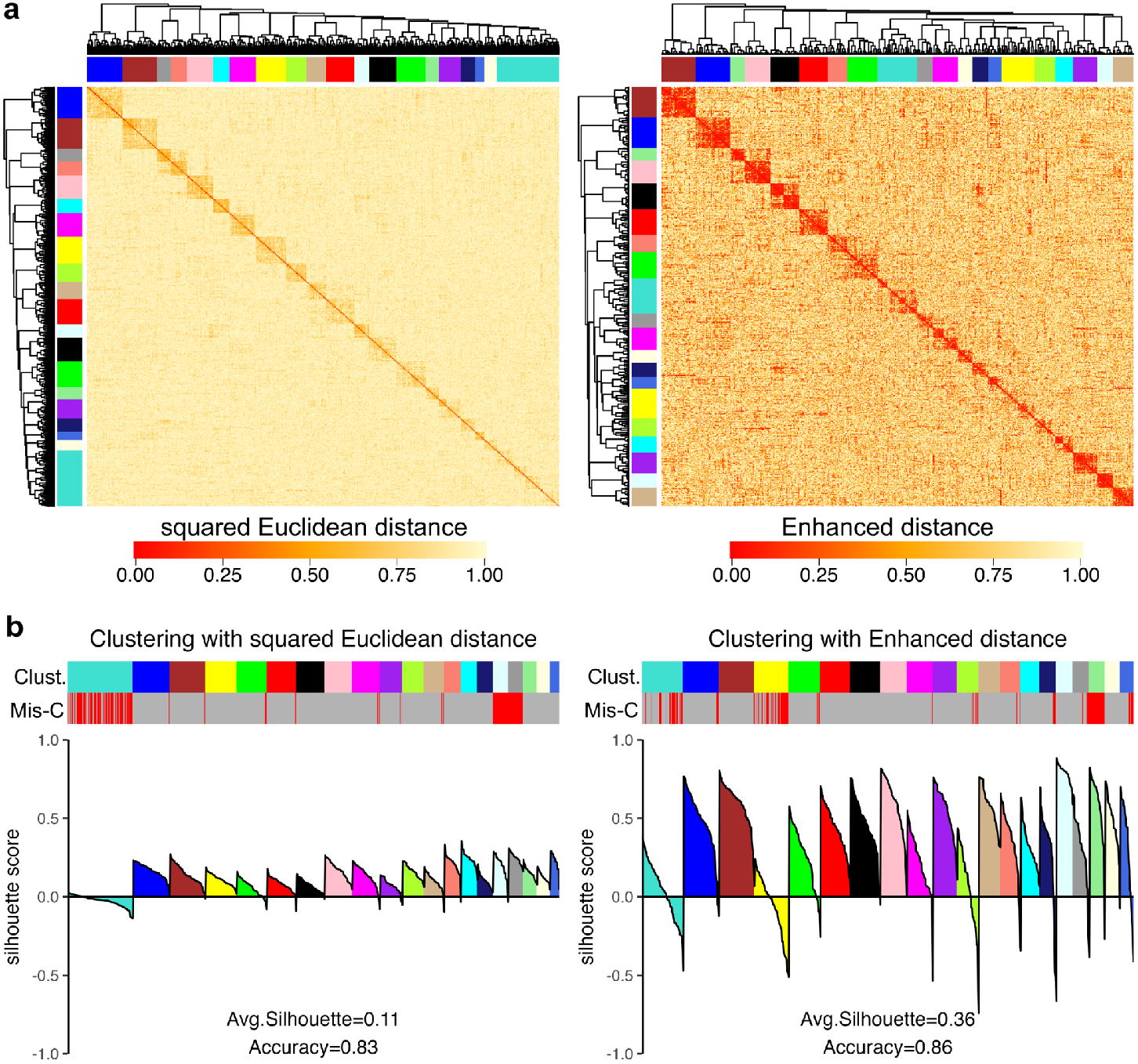
Ward’s clustering of simulated correlated data based on squared Euclidean distance without (left) and with (right) null model-based enhancement (Enhanced Distance, right) on a single simulated test dataset with 20 clusters and high SNR. (a) Heatmaps show correlation distances between analyte pairs; both distances are in the range [0, 1]. (b) Silhouette plots (bottom) depict where each analyte lies relative to the center of two clusters: its predicted cluster (at 1) and the nearest other cluster (at −1), with 0 equidistant from both. Clusters are sorted from most to least members; within each, analytes are sorted by decreasing silhouette score. Colored bars indicate predicted cluster (Clust., upper) and incorrect (red) or correct (grey) clustering assignment (Mis-C, lower) as described for computing accuracy.

### Full evaluation

We evaluated the relative usefulness of three new definitions of distance as well as two new definitions of network adjacency against the standard distance and standard WCNA adjacency using the previously described set of 600 test datasets. For convenience, we named these variants as follows. “HCLUST” refers to Ward’s hierarchical clustering method (R function `hclust()(The R Project for Statistical Computing)`) using squared Euclidean distance. “HCLUST-S” is the same, except that the correlation coefficients were standardized with null models (described above; see Methods) prior to conversion to a distance. “HCLUST-ES” refers to the same method as HCLUST-S, except that the standardized correlations were converted to Extended Distance rather than to squared Euclidean distance (described above; see Methods).

For comparison, we also included a distance-based method based on the most recent version (MOFA2) of Multi-Omics Factor Analysis(MOFA2), a recent unsupervised machine learning method designed specifically for integration and analysis of multi-omics data. MOFA2 is not specifically designed for clustering; instead it identifies latent factors that explain the variation shared across different omics layers using a Bayesian model, and is more directly comparable to Principal Component Analysis(Shlens 2014). Our “MOFA” method uses MOFA2 to analyze each test dataset and position each analyte on 30 latent factor coordinates, then uses the Ward hierarchical clustering method on squared Euclidean distances computed from the analyte positions (see Methods). We tested each distance-based method for a significant difference from the HCLUST method of the median, using the paired Wilcoxon signed rank test across 100 replicates.

For similarity-based clustering we tested two new definitions of adjacency for Weighted Correlation Network Analysis against “WCNA” (WCNA, run as described(Langfelder and Horvath 2008)). The standard WCNA procedure fits a power law constant *k* to convert squared Euclidean distances to network adjacencies, *a*(ρ|*k*) = (1 − ρ)^*k*^, then converts adjacencies to topological overlap measure (TOM) prior to hierarchical clustering. Method “WCNA-S” follows the same protocol, but begins with our Standardized correlations. Method “WCNA-M” uses our null Model-based adjacency function (Equation 3, above) instead of a power law, and continues the WCNA framework beginning with converting these adjacencies to TOM.

In our tests on simulated data (Fig. 3), we assessed clustering quality using “accuracy”, defined as the maximum fraction of correctly assigned analytes for any 1:1 matching of the predicted clusters to the true clusters. We tested each method against the standard method of the same type (HCLUST or WCNA) under 6 scenarios (see Testing Strategy, above). For a comparison with results using other metrics (“purity”, silhouette score), see Supplementary Figure 1.

**Figure 3.**
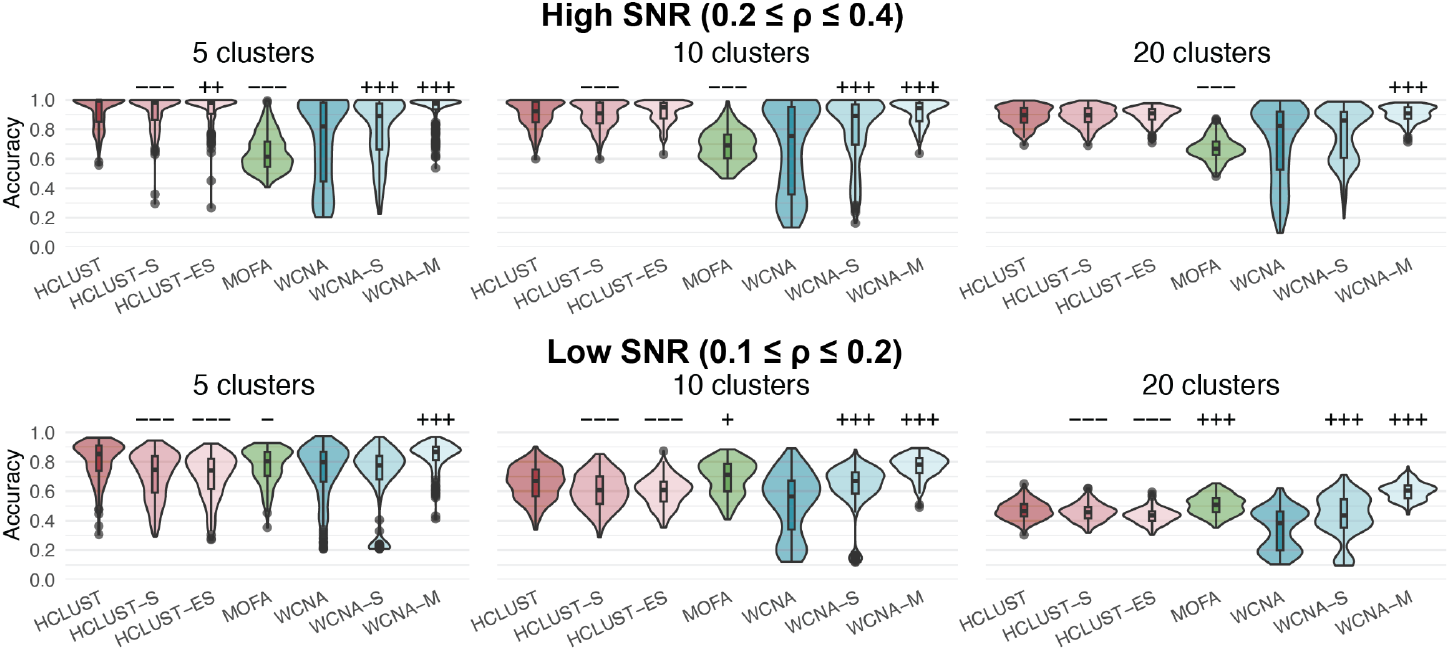
Comparison clustering accuracy for distance-based (HCLUST) and similarity-based (WCNA) clustering methods under different distances/adjacencies on simulated multi-omic data (see Methods). Each violin plot shows results for 100 replicate datasets under 6 scendarios: 5, 10, or 20 clusters (left, middle, right) with high SNR (average within-cluster correlations: 0.2-0.4 within-batch, 0.1-0.2 between-batch) or low SNR (average within-cluster correlations: 0.1-0.2 within-batch, 0.0-0.1 between-batch). *Correlation-based distances*: HCLUST (red): squared Euclidean (1 − ρ). HCLUST-S (pink): squared Euclidean using standardized correlations (1 − ρ_*std*_). HCLUST-ES (pink): Extended Distance using standardized correlations (see Methods). *Latent factor distance*: MOFA (green): squared Euclidean distance in a space of 30 latent factors. *Adjacencies*: WCNA (blue): (1 − ρ)^*k*^, for *k* fitted to data (see Methods). WCNA-S (blue): (1 − ρ_*std*_)^*k*^, *k* fitted to data, using standardized correlations. WCNA-M (pink): adjacencies computed from standardized correlations and the standardized null model. The reference method for comparisons was HCLUST (for distances) or WCNA (adjacencies). Significance assessed by Wilcoxon signed rank test of median accuracy against the reference method. Significance levels annotated as *p* < 0. 05 (+/-), *p* < 0. 01 (++/--), or *p* < 0. 001 (+++/---), without correction for multiple testing (30 tests).

In the comparison of distance-based methods, HCLUST-S is never better than HCLUST, and HCLUST-ES is typically worse than HCLUST-S. Thus, neither standardization nor Enhanced Distance improves distance-based clustering. When used for distance-based clustering, MOFA was significantly lower in median accuracy than HCLUST in the tests with high SNR (high SNR, *n* = 5: MOFA 0. 614 vs. HCLUST 0. 995, *p*=3.5e-33; *n* = 10: 0. 691 vs. 0. 923, *p*=6.1e-31; *n* = 20: 0. 669 vs. 0. 897, *p*=5.2e-34), but better in 2 of 3 tests with low SNR and 10-20 clusters (low SNR, *n* = 5: 0. 804 vs. 0. 853, *p*=2.2e-2; *n* = 10: 0. 712 vs. 0. 669, *p*=1.06e-2; *n* = 20: 0. 508 vs. 0. 464, *p*=1.2e-6). In contrast, in the similarity-based clustering tests WCNA-S was consistently as good or significantly better than WCNA (in 4 of 6 scenarios, *p* < 1e-3), and WCNA-M performed significantly better in all settings on both accuracy (*p* < 1e-3 for all 6 scenarios) and purity (*p* < 1e-3 for all 6 scenarios). We conclude that our new adjacency function was successful, and that standardization is more beneficial in similarity-based clustering than in distance-based clustering.

### Analysis of the Arivale dataset

We tested and compared HCLUST, HCLUST-S, HCLUST-ES, WCNA, WCNA-S, and WCNA-M on data collected as part of the Arivale wellness program(Price et al. 2017). This multi-omic dataset consists of 44 clinical tests, 921metabolites, and 274 proximity extension assay-based proteomics measurements (Olink panels CVD2, CVD3, and INF(Lundberg et al. 2011)) collected from participants just prior to the start of their wellness program (see Methods). We separated correlation coefficients according to the ‘omics batches of the analyte pair and fitted a null model to each (6 models), then standardized the correlations to a standard null model, ρ ~ 2*B*(ν_0_, ν_0_) − 1, with ν_0_ = 32 (Figure 4) and constructed a null model-based adjacency function (Figure 5). We then clustered the analytes by each method (see Methods).

**Figure 4.**
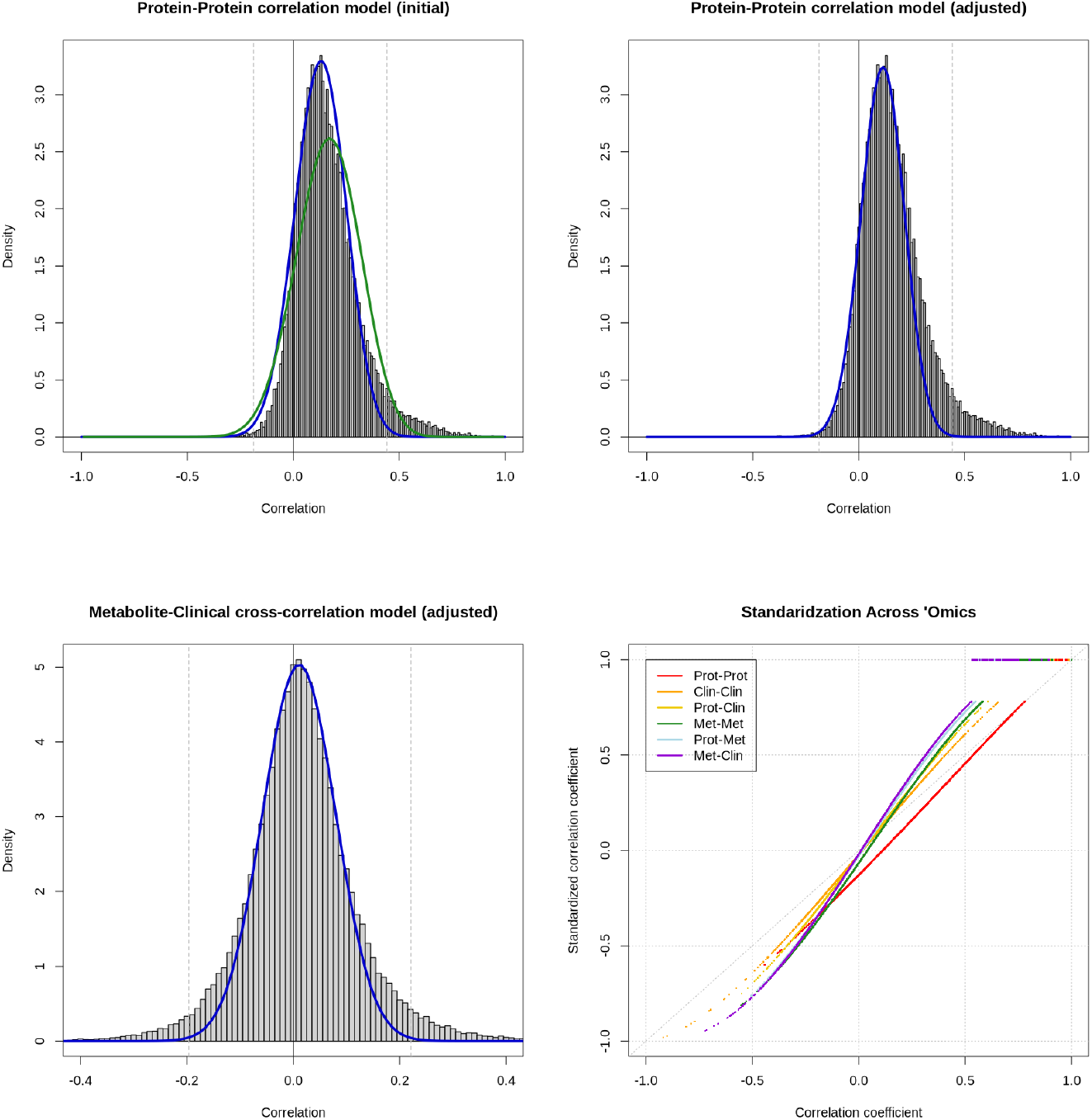
Standardization of Multiomic Correlations in Arivale. The multiomic Arivale dataset includes three ‘omics types and requires fitting of six null models. For correlations between protein analytes (upper left, upper right) the standard method of moments Beta distribution estimate (upper left, green) fits poorly; our heuristic function fits the center of the distribution better (upper left, blue) and can be further optimized by a user (upper right, blue). Correlations between a metabolite and a clinical test (lower left) have a different optimal fit (blue). An optimized fit for the metabolite-clinical test correlations is also shown (lower left, blue). Standardization maps optimized Beta null models for all six combinations of batches (lower right, x-axis) to a single null model (lower right, y-axis).

**Figure 5.**
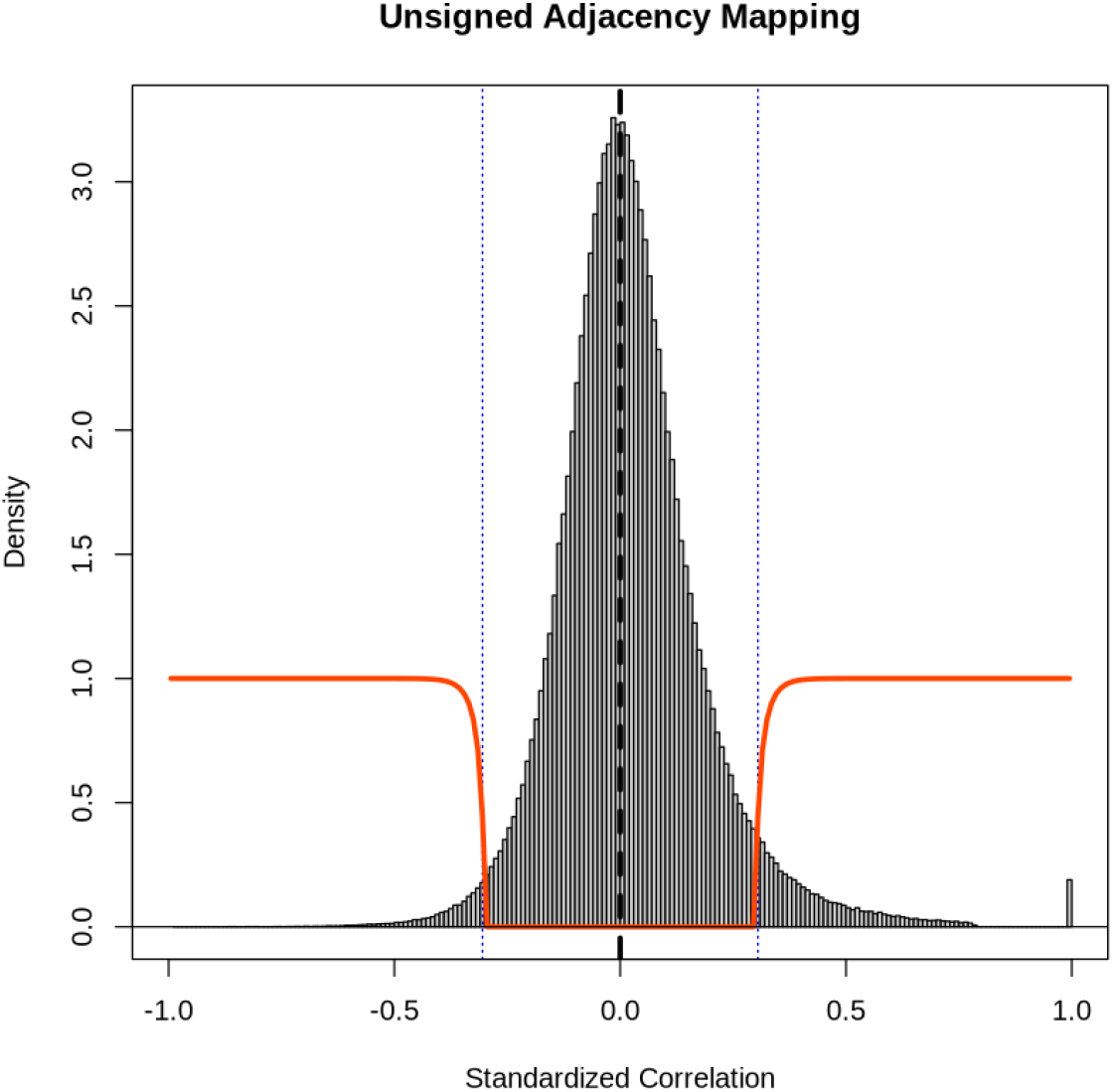
The Adjacency Function. The distribution of standardized correlations from the multiomic Arivale dataset (histogram) is overlaid with WCNA-M’s novel adjacency function (orange),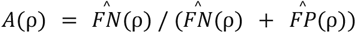, which represents the estimated likelihood that a misclassified correlation ρ should have been classified as an outlier. The null model dominates the central region (dotted blue lines), and the correlations in this region have nearly zero adjacency; the corresponding edges were omitted from the correlation network. The transition between high and low adjacency is not symmetric and sigmoidal, with a sharp decline driven by the rising probability under the null model. The adjacency function and null model are symmetric around 0, but more outliers to the right than to the left cause more positive than negative correlations to achieve high adjacency.

We evaluated predicted cluster separation using heatmaps and silhouette scores, and show here a comparison of WCNA with WCNA-M (Figure 6). The heatmap for WCNA showed low pairwise similarity both within and between most predicted clusters (Fig. 6a, left), and the heatmap for WCNA-M showed greater similarities overall (Fig. 6b, left). The silhouette scores (Figure 6, right) show that WCNA-M generally provides better separation between clusters. We also assessed the success of standardization by computing the entropy of analyte types across clusters (see Methods). A zero clustering entropy (S) indicates analytes within the same cluster were always of the same ‘omics type, with higher values indicating increased mixing across ‘omics. The clustering entropies confirm that WCNA-S and WCNA-M resulted in greater mixing of analytes across ‘omics types (WCNA, *S* = 0. 48; WCNA-S *S* = 0. 79; WCNA-M, *S* = 0. 74).

**Figure 6.**
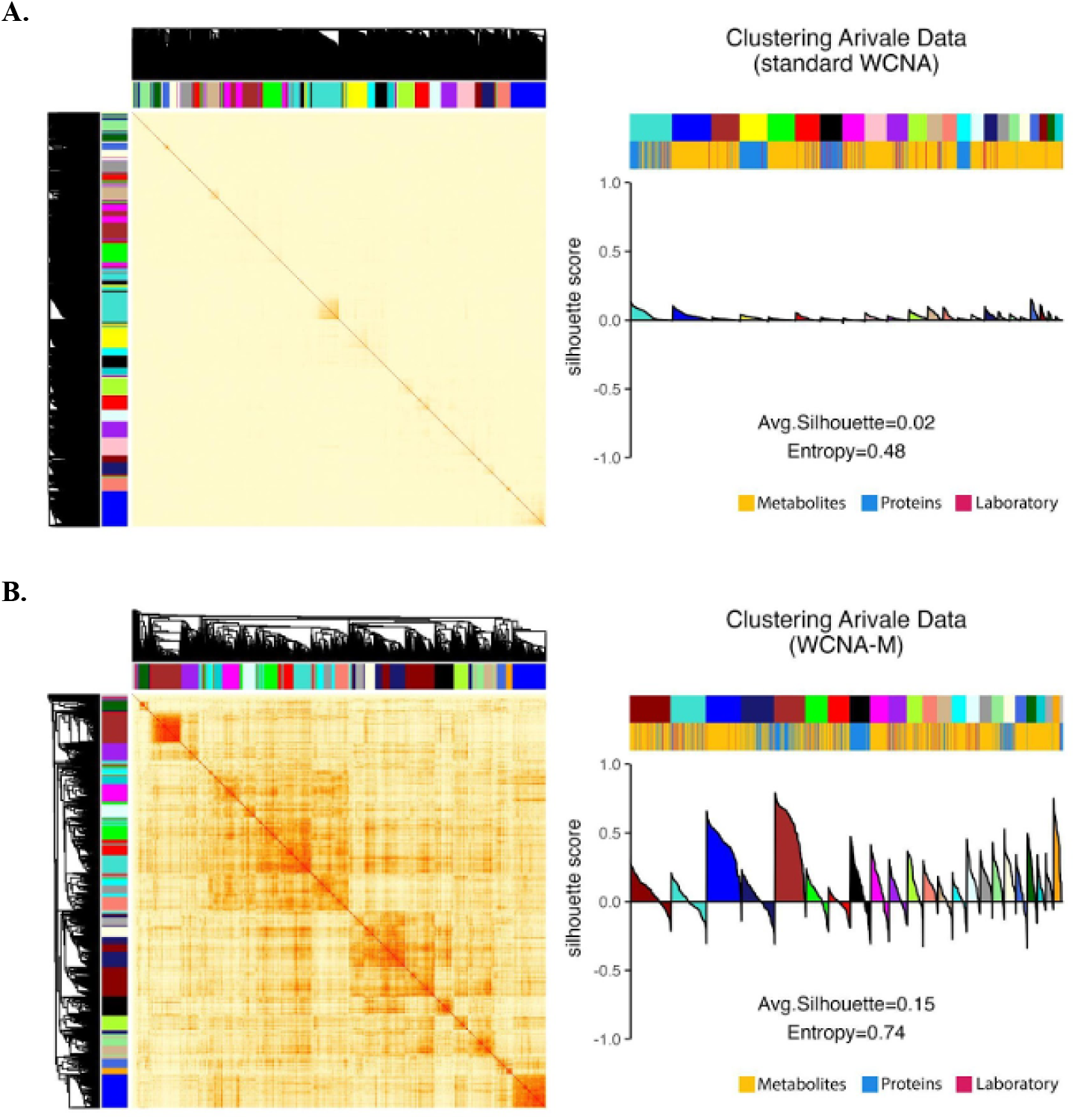
Correlation network construction on the Arivale dataset, using (A) standard WCNA and (B) WCNA-M. The heatmaps (left) show the TOM values used during clustering and dendrograms of the cluster structure; color bars (margins) show cluster membership. Paired colorbars (right, upper) show analyte cluster membership (top bar) versus analyte batch (lower bar: metabolites, yellow; proteins, blue; laboratory tests, pink); average entropies show WCNA-M provides better mixing of batches into clusters. The silhouette plot (right) presents silhouette scores in a decreasing order in each cluster. Deeper branching in the WCNA-M dendograms than the WCNA dendograms (left) show greater separation between clusters and correspond with higher average silhouette scores (right).

### Clusters associated with Age in the Arivale dataset

We then compared the biological relevance of each set of clusters by their ability to predict Age. For each method we predicted Age with a linear regression model, controlling for sex as a covariant and using the principal eigenvectors for all clusters as competing predictors:

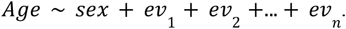

Since every analyte is included in some cluster by each method, the regressions which explained more variance in age identified cluster sets which grouped analytes better for explaining Age. For each cluster with a significant contribution to predicting Age (Table 1) we determined the module membership (MM)(Mao et al. 2020) of each analyte (the correlation of the analyte’s expression pattern with the cluster eigenvector), and the analyte with the highest MM was used to label the cluster (Table 1, Hub Analyte) and to identify similar clusters across methods.

**Table 1.** Regressions to Predict Age. Each cluster identified by any method (columns) as significant in the method’s regression to explain Age is labeled with its hub analyte (see text), and its regression coefficient is shown in years per standard deviation (n/s, not significant). The mean cohort age was 47; positive values indicate the cluster is associated with older participants, negative with younger. The total absolute value of the significant regression coefficients is also shown (Total Magnitude). The values shown are for clusters generated from Spearman coefficients, for which the variance in Age explained by the regression model is shown (Adj. R^2^ (Spearman)). The experiment was repeated using `bicor()`; only the explained variance is shown (Adj. R^2^ (bicor)).

| Hub Analyte | HCLUST | HCLUST-S | HCLUST-ES | WCNA | WCNA-S | WCNA-M |
| --- | --- | --- | --- | --- | --- | --- |
| DHEA-S | -5.1 | -4.7 | -4.4 | -5.1 | **-6.0** | -4.7 |
| UREA NITROGEN | 1.3 | **2.1** | 0.83 | 1.80 | 1.2 | 0.99 |
| Omega-6/Omega-3 Ratio | **2.4** | 2.3 | 2.0 | 2.2 | 2.1 | 2.2 |
| DMTPA | 1.9 | 3.2 | **6.2** | 3.1 | 3.6 | 5.2 |
| PGF | 1.9 |  | n/s | 1.2 |  | n/s |
| gamma-glutamylleucine | n/s | -3.7 | -3.7 | -1.4 | -1.4 | **-1.6** |
| alpha-hydroxyisocaproate |  |  |  |  |  | **-2.8** |
| MMP9 | 1.1 | n/s | n/s | n/s | 2.0 | **-0.96** |
| HOMA-IR | -2.2 | 2.1 | 1.7 | -2.7 |  | **3.3** |
| Triglycerides | -1.7 | n/s |  | -1.4 | -1.1 | **-1.2** |
| SIRT2 | -0.89 | -0.89 | n/s | -0.89 | **-1.1** | -0.89 |
| 1,7-dimethylurate | 0.7 | 0.92 | n/s | **0.98** | 0.96 | 0.81 |
| sphingomyelin | 1.1 | 1.00 | n/s | **1.4** | n/s | 1.0 |
| Shorter FA Carnitines | **1.4** | 1.9 | n/s | n/s | 1.7 | 0.85 |
| Longer FA Carnitines | **1.6** | n/s | n/s | 1.2 |  | 0.95 |
| taurodeoxycholate | **-0.76** | -0.72 | n/s | -0.66 | -0.69 | n/s |
| palmitoylcholine | 0.97 | 1.00 | n/s | n/s | 0.91 | n/s |
| catechol sulfate | 1.3 | n/s | n/s | n/s | n/s | n/s |
| PI3K/AKT Patway | n/s | n/s | n/s | n/s | -1.5 | n/s |
| Glycerophospholipids | n/s | n/s | -1.3 | n/s | n/s | -0.94 |
| Total Magnitude | 26.3 | 24.5 | 20.1 | 24.0 | 24.3 | 28.4 |
| Adj. R<sup>2</sup> (Spearman) | **0.460** | **0.483** | **0.493** | **0.474** | **0.495** | **0.523** |
| Adj. R<sup>2</sup> (bicor) | **0.491** | **0.493** | **0.522** | **0.472** | **0.509** | **0.537** |

**Table 2.** Cluster sizes and overlap. The size of each cluster in Table 1 is shown here along with the number of analytes common to the cluster containing the listed hub analyte across all methods (Every) and the number of analytes associated with the hub analyte by any method (Any). Clusters that were not significantly associated with Age in the regressions are greyed; the smallest cluster for each hub analyte is in bold.

| Hub Analyte | Every | HCLUST | HCLUST-S | HCLUST-ES | WCNA | WCNA-S | WCNA-M | Any |
| --- | --- | --- | --- | --- | --- | --- | --- | --- |
| DHEA-S | 25 | 32 | 34 | **31** | 41 | 71 | 46 | 75 |
| UREA NITROGEN | 24 | 77 | 44 | **33** | 43 | 40 | 42 | 95 |
| Omega-6/Omega-3 Ratio (-) | 24 | 33 | **31** | **31** | 41 | 63 | 37 | 76 |
| DMTPA | 22 | 56 | **52** | 105 | 83 | 107 | 64 | 178 |
| PGF | 13 | 125 | 84 | **29** | 45 |  | 65 | 212 |
| gamma-glutamylleucine | 11 | 64 | 87 | 97 | 105 | 105 | **49** | 185 |
| alpha-hydroxyisocaproate | 5 |  |  |  |  |  | **26** | 169 |
| MMP9 | 6 | 25 | 51 | 78 | 64 | 140 | **22** | 225 |
| HOMA-IR | 17 | 70 | 45 | 93 | 53 |  | 117 | 265 |
| Triglycerides | 22 | 40 | **35** |  | 69 | 73 | 71 | 175 |
| SIRT2 | 55 | 58 | 67 | 60 | 131 | 142 | 96 | 173 |
| 1,7-dimethylurate | 20 | **20** | **20** | 38 | 24 | 28 | 22 | 48 |
| sphingomyelin | 15 | **43** | 44 | 59 | 56 | 61 | 77 | 101 |
| Shorter FA Carnitines | 13 | 56 | 58 | 90 | 113 | **40** | 128 | 160 |
| Longer FA Carnitines | 5 | **18** | 32 | 44 | 45 |  | 30 | 79 |
| taurodeoxycholate | 20 | **22** | 27 | 32 | 28 | 30 | 33 | 35 |
| palmitoylcholine | 13 | **19** | 22 | 67 | 23 | 24 | 41 | 84 |
| catechol sulfate | 12 | 120 | 38 | 47 | 35 | **27** | 33 | 144 |
| PI3K/AKT Patway | 34 | 56 | **42** | 45 | 77 | 51 | 48 | 87 |
| Glycerophospholipids | 11 | 41 | 74 | 52 | 37 | 36 | **26** | 110 |

Of the WCNA strategies, WCNA-M explained the most variance in age (Table 1, Adjusted *R*^2^ = 52. 3%), followed by WCNA-S (49. 5%) and standard WCNA (47. 4%). Of the distance-based strategies, HCLUST-ES explained the most variance in age (49.3%), followed by HCLUST-S (48.3%) and HCLUST (46.0%). In both the distance-based and similarity-based settings, standardization of the correlations improved the biological relevance of the clusters (WCNA-S 2.1% over WCNA; HCLUST-S 1.3% over HCLUST), as did the use of the null model (WCNA-M 2.8% over WCNA-S; HCLUST-ES 1.0% over HCLUST-S). WCNA-M provided substantially more improvement over WCNA (6.3%) than WCNA provided over HCLUST (1.4%), which suggests that the use of the null model. Repeating the entire experiment using `bicor()` in place of Spearman correlations (Table 1, last row) produced similar results (WCNA-M over WCNA, 6.5%; WCNA and HCLUST essentially equal).

## Methods

### Simulating correlated data

For correlation-based clustering, we assumed that analyte-analyate correlations follow a structured relationship: intra-cluster correlations are greater than inter-cluster correlation, and correlations among analytes from the same data type tend to be higher than those between different data types within the same cluster.

We generated 100 replicate test data matrices for each of 6 test scenarios: *C*=5, 10, or 20 “true” clusters intercorrelated with either “high” or “low” targeted signal to noise (SNR), 600 total datasets. Each dataset was a 100 × 800 matrix of expression levels for *m* = 100 samples and *n* = 800 analytes, with *n* = 400 analytes assigned to batch “A” and *n* = 400 analytes assigned to batch “B” to represent two different ‘omics measurement platforms.

Our approach was designed to produce three different kinds of correlations between analytes: (a) within-cluster, within-batch correlations, (b) within-cluster, between-batch correlations, and (c) between-cluster correlations; for each type, the correlations were designed as a covariance matrix and created using the Cholesky decomposition method(Kaiser and Dickman 1962) (see also Supplement S4) on a set of random vectors.

### Within-cluster, within-batch analyte correlations

In the first stage, each cluster and batch was treated as a separate group. Each group represents some number *n*_*c*ω_ of analytes, where 1 ≤ *c* ≤ *C* identifies the cluster and ω ∈ {*A, B*} identifies the batch. We generated a target average correlation μ_*c*ω_ for the group according to the targeted SNR: μ_*c*ω_ ~ *U*[0. 2, 0. 4] for high SNR scenarios, or μ_*c*ω_ ~ *U*[0. 1, 0. 2] for low SNR scenarios. We then generated a group-specific (*n*_*c*ω_ × *n*_*c*ω_)symmetric, positive definite correlation matrix Ω_*c*ω_ = {ρ_*ij*_}, with diagonal 1 (ρ_*ij*_ = 1) and off-diagonal elements ρ_*ij*_ = ρ_*ij*_ ~ *N*(μ_*c*ω_, *s*_*c*ω_), 1 ≤ *i* < *j* ≤ *n*_*c*ω_, where *s*_*c*ω_ = min {0. 5 μ_*c*ω_,0.1}. The Cholesky Decomposition of Ω_*c*ω_ was then multiplied by an iid, {*m* × *n*_*c*_ω} matrix of *iid* standard Normal random numbers to generate *n*_*c*ω_ expression profiles, each containing the designed correlations to each other. When every cluster and batch is finished, an initial expression profile *E*_*i*_ has been produced for every analyte.

### Within-cluster, cross-batch correlations

The designed correlation between analytes in the same cluster *c*, but from different batches, is added using a (2 × 2) correlation matrix Ω_*c*_ = {1, ρ _*AB*_; ρ_*BA*_, 1}, where ρ_*AB*_ = ρ_*BA*_ ~ *U*[0. 1, m*i*n {μ_*cA*_, μ_*cB*_}]. For each pair (*i, j*) of analytes in cluster *c* with analyte *i* from batch A and *j* from batch B, the (2 × 2) Cholesky Decomposition of Ω_*c*_ is multiplied by the (*m* × 2) matrix [*E*_*i*_; *E*_*j*_], producing updated expression profiles *E*_*i*_^’^ and *E* _*j*_^’^. This process is repeated, resampling ρ_*AB*_ each time and updating every analyte’s expression profile to incorporate every within cluster, between batch correlation; this is a linear process and the correlations are additive, so the existing correlations are preserved through each update.

### Cross-cluster correlations

The same strategy just described is used to add cross-cluster correlations using (2 × 2)designed correlation matrix Ω_*_ = {1, ρ_*_; ρ_*_, 1}, where ρ_*_ ~ *N*(0, 0. 05) is resampled for every pair of analytes from different clusters.

### Arivale dataset

The Arivale cohort consists of individuals over 18 years of age who between 2015 and 2019 self-enrolled in a now closed scientific wellness company(Price et al. 2017). Briefly, about 80% of Arivale participants were residents of Washington or California, and about 60% of the cohort is female. For this study, only baseline measurements were considered for each participant, and only fully-characterized metabolites were included. We excluded low-data analytes and samples with goodSamplesGenes()(Horvath 2011); one sample was subsequently found to have zero variance across analytes and removed. After cleaning, 274 proteins from O-link panels CVD2 (92), CVD3 (91), and INF (91) remained, as well as 921 MS-based metabolite measurements and numerical results from 44 clinical laboratory tests. These values were then log-transformed (log_2_ (1 + *x*)) to form Arivale data matrix *M*.

### Correlations

For test datasets or the Arivale data matrix with samples on the rows and analytes on the columns, we computed a matrix of analyte-analyte Spearman correlations *Z* `= SparseSpearmanCor2(M)` from data matrix *M*; correlation coefficients were identical to those computed with `cor(M,method=“spearman”`)(The R Project for Statistical Computing). When repeating the analysis on `bicor()` correlations, we computed the biweight midcorrelation matrix for the Arivale dataset *Z* = `WGCNA::bicor(M)`.

### Null models for correlation coefficients

We used `estimateShape()` to assist in finding a resistant estimate for each correlation type. For the simulated datasets, we set both left and right parameters to k=13, as adjustments to the estimated values of w and v in Beta(w,v). Plots of the initial and final (hand-adjusted) versions of all six Beta null models for the Arivale dataset are available in Supplement S5. The hand-adjusted parameters were (P, proteins; M, metabolites; C, clinical tests): P-P model, *B*(52. 480, 42. 206); M-M, *B*(97. 651, 90. 067); C-C, *B*(63. 289, 61. 146); P-M, *B*(152. 684, 148. 652); P-C, *B*(101. 471, 98. 200); M-C, *B*(107. 892, 105. 589). The mapping of these models to the reference null model are shown in Figure 4, lower right.

### Standardized correlations

Each type of correlation coefficients is mapped from its fitted null model to a single, symmetric Beta null model, Beta(ν_0_,ν_0_), using `standardizeFromModel()`; we refer to the resulting values as standardized correlation coefficients. The reference shape parameter ν_0_ is arbitrary, but we recommend choosing one which is neither too narrow, reducing resolution within the probable range for the null model, nor too wide, reducing resolution in the region containing the outliers of interest. We used ν_0_ = 32. Theoretically, the value of ν_0_ affects neither distance penalties nor adjacencies, so long as the same ν_0_ is used for both purposes. This was verified using an alternative value (ν_0_ = 36).

### Enhanced distances

A correlation coefficient ρ is commonly converted to squared Euclidean distance *d*(ρ) = 1 − ρ for distance-based clustering. We define the corresponding Enhanced Distance *D*_*v,w*_ (ρ), computed by `betaDistance()`, for a correlation coefficient ρ with null distribution Beta(*v,w*) to be

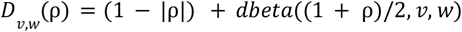

for an unsigned interpretation of ρ, where *dbeta*() is the Beta probability density function(The R Project for Statistical Computing), or

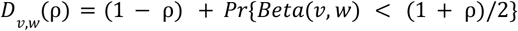

for a signed interpretation of ρ; *Pr*{*Beta*(*v, w*) < *x*} is implemented by the R function *pbeta*(*x*, ν, ω)(The R Project for Statistical Computing).

### Adjacencies from the null model

We define an adjacency function which is the estimated likelihood that a misclassified correlation ρ should have been classified as an outlier:

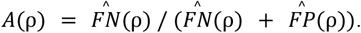

This soft threshold adjacency is centered (*A*(ρ) = 0. 5) where the estimated number of false negatives *FN*(ρ) and false positives *FP*(ρ) is equal. *FN*(ρ) and *FP*(ρ) are estimated from a known null model *M*_0_ as follows (the definition does not require *M*_0_ to be a Beta distribution). Given *K* unique, non-self pairwise correlations, with an unknown number *N* drawn from the null distribution *M*_0_ (negatives) and all others being “outliers”, we estimate *N* as twice the number of observations in the interquartile range of *M*_0_. Then the number of false positives at threshold *t* is

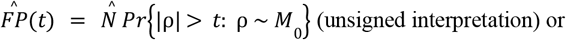

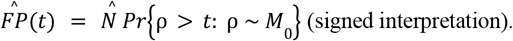

The estimated number of outliers is 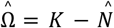, and since every outlier is either a true positive or a false negative,

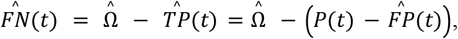

Where 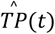 is the estimated number of outliers stronger than the threshold and we can count the number of correlations stronger than the threshold,

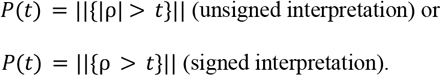

The function `nullModelAdjacencyTable()` performs this estimation, since any outliers in the interquartile range of *M*_0_ will cause underestimation of the number of outliers Ω, a `scale` parameter is provided; by default `scale=1`.

### Clustering

We tested both distance-based and WCNA-type clustering: four methods based on Ward’s minimum variance distance-based clustering (HCLUST, HCLUST-S, HCLUST-ES, and MOFA), and three based on Weighted Correlation Network Analysis (WCNA, WCNA-S, and WCNA-M).

### Distance-based clustering

Spearman correlation coefficients {*Z* = *z*_*ij*_} were computed for all analyte pairs and converted to squared Euclidean distance, *D* = 1 − |*Z*|. Method “HCLUST” used distance matrix *D*.

Method “HCLUST-S” instead computed distance from standardized correlations, *D* = 1 − |*Z*_*c*_ |.

Method “HCLUST-ES” used Enhanced Distance computed as

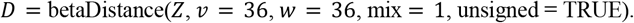

All distance clustering methods used R function `hclust(method=“Ward.D2”)` on *D* to produce a hierarchical clustering tree which was cut into the desired number of clusters *n* using `cutree(k=n)`.

### MOFA analysis

Multi-omic Factor Analysis (MOFA2) is a Bayesian factor analysis method which identifies latent factors of variation across multi-omics datasets(Argelaguet et al. 2018; Gagolewski 2025). We used the `run_mofa()` and `get_weights()` functions on each of our simulated datasets to produce a matrix of coordinates for each analyte on *k* = 30 latent dimensions. We then computed squared Euclidean distances between the analytes in latent space and clustered the analytes using Ward’s method using `hclust()` and `cutree()` in R(The R Project for Statistical Computing), producing as many clusters as there were analyte groups (*n* = 5, 10, 20) in the simulated dataset.

### Weighted Correlation Network Analysis

Our WCNA-based clustering methods are variants of the method described in the standard WGCNA tutorial(Horvath 2011), which we refer to as method “WCNA”:

1. Compute correlation coefficients with `SparseSpearmanCor2()`.
2. Convert correlations to similarities: `1-squaredEuclidean()`.
3. Find a power *k* with `WGCNA::pickSoftThreshold.fromSimilarity()`.
4. Convert similarities to adjacencies using the suggested power function, *x*^*k*^: `WGCNA::adjacency.fromSimilarity()`.
5. Calculate TOM from the adjacencies, `WGCNA::TOMSimilarity()`.
6. Cluster using `hclust()` using `as.distance(1-TOM)` and `“average”` linkage.
7. Cut the clustering tree into initial clusters with `cuttreeDynamic(cutHeight = 1, deepSplit = 4,pamRespectsDendro = FALSE,minClusterSize = 20)`. This function assigns each analyte a cluster, represented by an integer 1, …, *C*, where *C* is the number of clusters.
8. For convenience, we converted integer cluster labels to named colors with `labels2Colors()`.
9. As recommended, we reduced the number of clusters by merging similar clusters, using `moduleEigengenes(nPC=2)` to find representations of the clusters, and `mergeCloseModules(cutHeight=0.25)` to assess the similarities and merge the most similar.
10. The merged clusters are the final clustering, with each analyte assigned to one cluster. For further analysis, a compact representation of each final cluster is their principal eigenvector; these eigenvectors are given by `moduleEigengenes()`$`eigengenes`.

Method “WCNA-S” applies this protocol to standardized correlations (described above) beginning at step 3 (fitting a power parameter). Method WCNA-M uses standardized correlations to tabulate an adjacency function with `nullModelAdjacencyTable()`, then interpolates an adjacency value from the table using `interpolatedAdjacency()` to convert the standardized correlation matrix to an adjacency matrix. Clustering continues following the protocol above starting at step 5 (calculating TOM).

### Clustering assessment

For simulated data, the correct assignment of analytes to groups is known and we evaluated a clustering of *N* analytes assigned to *n* true clusters and *n* predicted clusters using both optimal set-matching accuracy and purity(Berg and Lässig 2006). We first computed the confusion matrix {*C* = *c*_*ij*_}, where *c*_*ij*_ is the count of analytes in predicted cluster *i* and true cluster *j*. Accuracy is the fraction of correctly matched elements under the best one-to-one assignment of predicted to true clusters,

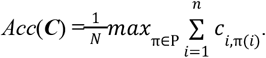

For efficiency, we used `GraphAlignment::LinearAssignment()`(“Finding Groups in Data”: Cluster Ana…) to identify the maximizing permutation π. Purity is the fraction of correctly assigned analytes allowing assignment of multiple clusters to each group,

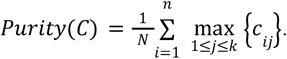

For real datasets, groups are undefined and we evaluated clustering performance using silhouette score (cluster::sihouette()(“Finding Groups in Data”: Cluster Ana…)), averaged over all analytes. The silhouette score (− 1 ≤ *s*(*q*) ≤ 1) of an analyte *q* measures how close the analyte is to its own cluster relative to the nearest other cluster,

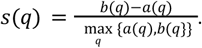

Here *a*(*q*) is the mean distance between *q* and the other analytes in the same cluster, and *b*(*q*) is the mean distance between *q* and the members of the nearest other cluster; for WCNA-based methods, we used distance = 1 - adjacency. A clustering with lower average silhouette score for the same data indicates less separation between clusters.

We computed clustering entropy *S* as the average per-cluster analyte type entropy,

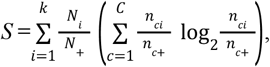

where the analytes are divided into *k* batches (‘omics types), *N*_*i*_ is the number of analytes in batch *i*, the analytes are assigned to *C* nonempty clusters, *n*_*ci*_ is the number of analytes of type *i* in cluster *c*, and a subscript + indicates summation over the corresponding index.

## Discussion

Multi-omic datasets are powerful for deciphering complex biological systems but present unique analytical challenges, particularly for correlation-based clustering methods like Weighted Correlation Network Analysis (WCNA). Our work addresses these challenges by proposing a novel framework for standardizing correlations across batches and converting correlations to distances or adjacencies as required by the clustering method. This framework is based on null models of high-dimensional correlations and is implemented in a new R package, standardcor(standardcor). We have given what is to our knowledge a novel argument, based on the formula for Pearson’s sample correlation coefficient, that the appropriate null model for correlation coefficients is a Beta distribution, whether or not the data is expected to be uncorrelated on average. We have shown (Fig. 1) that the null model accounts for the data’s geometric dimension, whether the weights produced are interpreted as distances or as network adjacencies.

Our findings demonstrate the significant advantages of incorporating null models in multi-omic clustering. In the Arivale dataset, we observe that correlations between analytes measured on the same platform tend to be larger than analytes measured on different platforms. We expect this reflects batch-specific experimental sources of variation, which are shared within a batch but differ between batches. We have provided a means to tightly fit Beta distributions to the bulk of a set of correlations, even in the presence of outliers, to create null models specific to each within-batch or between-batch category of correlation coefficients. These null models allow for the standardization of correlations with different statistical properties onto a single null model, providing cross-batch comparability and mitigating the bias towards single-batch clusters. Importantly, standardization improved batch mixing in clusters for the Arivale dataset, in which analytes are related by biological processes.

Standardized correlations also led to more meaningful clusters for explaining a complex biological phenotype (Age). In our most predictive model (WCNA-M), we observed clusters characterized by many analytes with known associations with age and mortality. For example, 2,3-dihydroxy-5-methylthio-4-pentenoate (DMTPA) is the representative analyte for the cluster with the largest estimated association with age, over 5 years per standard deviation (+5.2 y/sd). This cluster also contains Growth and Differentiation Factor 15 (GDF15), which is recognized as a key player in aging(Conte et al. 2022) and hypertension(Al Ashmar et al. 2024), as well as metabolites kynurenine, N6-carbamoylthreonyladenosine, and dimethylarginine (ADMA and SDMA), all of which (like DMTPA) are linked to hypertension(Nagy et al. 2017; Sökmen et al. 2019). Another large-effect cluster (−4.7 y/sd) is represented by dehydroepiandrosterone sulfate (DHEA-S), which has been suggested as a biomarker for aging(Urbanski 2021; Ohlsson et al. 2010); this cluster includes cortisol and the majority of androgens and estrogens. The decreasing trend in these hormones with age relative to the constant level of cortisol is known as adrenopause(Papadopoulou-Marketou et al. 2000). The clinical test HOMA-IR, which at high levels is an indicator of insulin resistance(Matthews et al. 1985) and is correlated with chronological and biological age(Yang et al. 2023), represents a cluster with an effect of +3.3 y/sd. This cluster also includes insulin, insulin-like growth factor binding proteins 1 and 2, and paraoxonase 3, all of which are linked to insulin biology(Rull et al. 2012).

These examples show that in addition to an association with age or aging, each significant cluster is indicative of a biological pathway associated with age and causes of mortality. The same appears to be true for the other clusters with a significant association with age, represented by alpha-hydroxyisocaproate (−2.8 y/sd), an end product of leucine metabolism and associated with exercise, muscle growth and cartilage growth(Mero et al. 2010; Tran et al. 2015); the Omega-6/Omega-3 ratio clinical test (+2.2 y/sd), which is linked to increasing telomere length(Seo et al. 2022) and increased risk of metabolic syndrome(Jang and Park 2020); gamma-glutamyl leucine (−1.6 y/sd), which is linked to higher cardiometabolic disease risk factors(Wu et al. 2022); the clinical test for Triglycerides (−1.2 y/sd), which is involved in both lipid metabolism and in energy metabolism(Spitler and Davies 2020) (this cluster also includes the low density lipoprotein receptor(Lee et al. 1982; Ericsson et al. 1991)); as well as 8 other statistically significant clusters with an estimated effect of 1 y/sd or less (Table 1) represented by the Urea Nitrogen (BUN) test(Liu et al. 2021), Matrix Metalloproteinase 9 (MMP-9)(Cancemi et al. 2020), Sirtuin 2 (SIRT2)(Garmendia-Berges et al. 2023), 1,7-dimethylurate (a marker of caffeine consumption), sphingomyelin(Mielke et al. 2015), shorter fatty-acylcarnitines(Jarrell et al. 2020), longer fatty-acylcarnitines(Jarrell et al. 2020; Caballero et al. 2021), and glycerophospholipids(Mutlu et al. 2021; Possik et al. 2023). It is striking that a systems biology approach including multi-omic measurements is able to connect proteins, metabolites, and clinical test results as multifaceted views into such a variety of aging-related biology.

Our null model-based approach offers a rigorous statistical model for separating signal from noise in correlation-based clustering. This contrasts with the power law transformation proposed with WCNA, which is motivated by the observation that natural networks are often scale-free and does not specifically address the significance of the correlations on which the network model is constructed. While our translation of the null model into distance-based clustering was less successful, particularly on simulated test data, we note that both distance-based and similarity-based uses of the null model framework provided measurable improvements over other methods on actual experimental data in both of the ways the null models were used (for standardization of correlations and for providing distances or adjacencies, as required by the clustering method).

Interestingly, the enhanced distance approach provided improved clustering on the Arivale dataset, but not on the simulated data; in contrast, our adjacency function and WCNA provided an improvement in clustering on both simulated and actual data. We note two important differences which together explain this phenomenon, and add to our understanding of these results.

The Arivale dataset and our simulated data differ in the correlation structure of the clusters. Briefly, we simulated test data with correlations between each pair of analytes in the same cluster; there was no mechanism which caused the correlations between different analyte pairs to involve a particular subset of samples, or to resemble each other in any way. We should not expect a single expression pattern (eigenvector) to represent a cluster in our test data. In contrast, the clusters identified in the Arivale data are each characterized by an eigenvector with which most analytes in the cluster are strongly correlated (or anti-correlated). The clusters in the Arivale data are “simpler” than the clusters in the test data, in the sense that less information is required to describe the correlation structure. In other words, the Arivale clusters are lower-dimensional than the clusters in the simulated data. Our enhanced distance and adjacency function are based on the distributional properties of the correlations, which is sensitive to dimensionality but not to individual, pairwise correlations. Thus, we expect the null model-based approach to be sensitive to correlations centered on a single pattern, but not necessarily to a dense set of correlations with no pattern in common.

WCNA and distance-based clustering differ primarily by WCNA’s use of Topological Overlap Measure (TOM). TOM uses pairwise correlations to identify densely connected regions of the adjacency graph; however it does not depend on whether these pairwise correlations involve similar sets of samples, or are associated in any way other than involving a restricted set of analytes. TOM is therefore sensitive to both the independent correlations we designed into our test data, and the correlations of multiple analytes with a common eigenvalue present in the Arivale data. Our results correspond precisely to our expectations, based on our understanding of the two methods used to incorporate non-pairwise information. In fact, our results show that our methods and TOM identify different non-pairwise information, and both contribute to identifying meaningful clusters in the Arivale data.

While our null model-based framework offers substantial improvements, certain limitations warrant consideration. Fitting Beta null models accurately relies on the assumption that the bulk of analyte pairs exhibit spurious correlation, and the parameter estimation method, though outlier-resistant, still requires careful evaluation and supervision. Further validation across a broader range of multi-omic data types and biological contexts will strengthen the generalizability of our findings; we expect such validation to occur rapidly, as we are aware of a number of projects that are already using package `standardcor`. A quantitative measure of the fit of the Beta model to the targeted middle half of an observed set of correlation coefficients would assist the user in refining the null model, and might make the fitting process more automatic. A more complete investigation of null model-based adjacencies within the rich WCNA ecosystem is also desirable. We look forward to these extensions of this work.

## Supporting information

Guide to Supplemental Materials

## Acknowledgements

We thank C. Funk, G. Glusman, J. Roach, A. Baumgardner, and N. Price for helpful discussions throughout the course of this project. We thank A. Magis for assistance obtaining Arivale data. We also thank M. Brunkow for her coordination efforts.

## Author contributions

M.R. conceived of the approach and was primary method developer. L.P., H.N., and N.R. performed method review and suggested improvements. H.R. generated simulated data and managed data handling. M.R., L.P., H.N., and N.R. participated in study design. M.R. and H.N performed data analysis and figure generation. M.R. and N.R. interpreted results. M.R. and H.N. were the primary writers of the paper, with contributions from all authors. All authors read and approved the final manuscript.

## Competing interests

The authors declare no competing interests.

### IRB Approval

Arivale Cohort: Procedures for the current study were run under the Western Institutional Review Board (WIRB) with Institutional Review Board (IRB) Study Number 20170658 at the Institute for Systems Biology and 1178906 at Arivale (both in Seattle, WA).

### Cohorts

The Arivale cohort consists of individuals over 18 years of age who between 2015 and 2019 self-enrolled in a now closed scientific wellness company. Briefly, the majority of Arivale participants (~80%) were residents of Washington or California when in the program. While the cohort tends to be healthier than the general US population, it is more reflective of populations in these two states^40^. The cohort is also predominantly female (~60%), which may be due to women being more likely to join these types of programs. For this study, only baseline measurements were considered for each participant, and individuals who provided a stool sample were included in the analysis.

## Statistical Analysis

## Code Availability

Code used for statistical analysis is available through GitHub (https://github.com/longevity-consortium/standardcor).

## Data availability

Qualified researchers can access the full Arivale deidentified dataset supporting the findings in this study for research purposes. Requests should be sent to.

## Footnotes

1 We anticipate changing the definition of Extended Distance to use the more successful strategy used in WCNA-M (below) in the near future.

