## Supplementary material for "Model-based Standardization of Correlation Coefficients Improves Multi-Omic Clustering and Biological Signal Discovery": Guide to Supplemental Materials

### Guide to Supplementary Materials

for the article

*Correlation Null Models Provide Improved Cluster Separation and Reduce Biases in Concatenated Clustering of Multiomic Data*

by M. Robinson, L. Pflieger, H. Noh, and N. Rappoport

September 2025

- **Supplementary Figure 1**
  Provides a comparison of the accuracy results presented in Figure 3 with two other metrics: purity, which uses a different assignment of predicted to true clusters, and average silhouette score (see Figure 2).
- **Supplement S1**
  Discusses the geometry of correlation coefficients, and the advantage provided by the Topological Overlap Measure used in WCNA. We provide a derivation of the form for our Beta null model for correlation coefficients, and comment on why we believe TOM and the Beta null model should be used together.
- **Supplement S2**
  Student's $t$ test is an exact test of statistical significance for correlation coefficients which applies when $E[\rho]=0$. When $-1<E[\rho]<1$, the standard approach is to use Fisher's transformation to provide a $z$-statistic; this statistic is approximately Gaussian, and converges asymptotically to a Gaussian distribution as the sample size increases. We show that our Beta null model provides an exact test that is identical up to rounding error with the $t$-test when it applies. When $-1<E[\rho]<1$ and the marginal distribution is Gaussian, he Beta test epending on the marginal distribution,  there is good agreement between the Beta test and the approximate test using Fisher's transformation is approximately Gaussian. We illustrate the differences between Fisher's transformation and the Beta test when Fisher's transformation is not expected to be accurate.
- **Supplement S3**
  Student's *t* and Fisher's transformation are applied under the assumption that the data has iid Gaussian marginal distributions. We illustrate by simulation whether the Beta null model appears to be exact, asymptotically correct, or incorrect for a variety of *iid* marginal distributions.
- **Supplement S4**
  We explain our use of the Cholesky Decomposition.
- **Supplement S5**
  For each of the six null models generated for the multi-omic Arivale dataset, we show the fit of two models to the observed correlation coefficients to the unique, non-self analyte pairs (including only one of $(a,b)$ or $(b,a)$ for analytes $a \neq b$ and excluding pairs $(a,a)$): the model fitted with `estimateShape()` with default parameters, and the model fitted by hand using nonzero values for the parameters `left` and `right`.
