## Supplementary Figure 1 for "Model-based Standardization of Correlation Coefficients Improves Multi-Omic Clustering and Biological Signal Discovery"

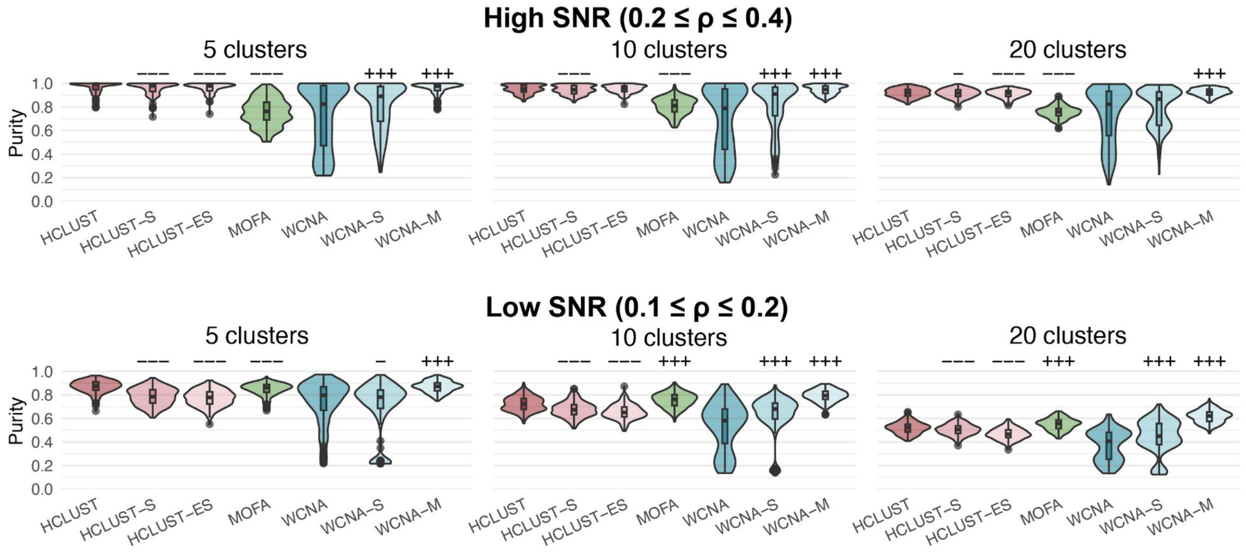


**Supplementary Figure 1.** Comparison of the same clustering methods shown in Figure 3 (main paper) using the "purity" metric, an alternative to the "accuracy" metric used in Figure 3. Purity is calculated with the same predicted cluster permitted to match more than one "true" cluster, while Accuracy requires a 1-1 matching between predicted and "true" clusters (see Methods). The organization and color of the violin plots here is identical to Figure 3. As in Figure 3, each method was annotated as having significantly higher/lower median value than the reference method (HCLUST or WCNA) at the $p<0.05$ (+/-), $p<0.01$ (++/--), or $p<0.001$ (+++/---) significance level, without correction for multiple testing (30 tests). The results with purity and accuracy are consistent, except for HCLUST-ES with 5 clusters, High SNR.
