## Supplement S1 for "Model-based Standardization of Correlation Coefficients Improves Multi-Omic Clustering and Biological Signal Discovery"

### *The geometry of correlation coefficients*

A correlation coefficient characterizes the relationship between two variables in a paired (statistical) sample, $(X,Y)=\{(x_i,y_i): i=1..n\}$. In our case the variables $X$ and $Y$ represent different analytes measured over the same set of $n$ (biological) samples, and we will consider $X$ and $Y$ to be points in the same $n$-dimensional space to discuss pairwise correlation from a geometric perspective.

Pearson's correlation coefficient is defined on the paired sample as a sum of products,

$$\rho(X,Y)=\frac{1}{n-1}\sum_{i}\left(\frac{x_i-\underline{x}}{s_X}\right)\left(\frac{y_i-\underline{y}}{s_Y}\right)=\frac{1}{n-1}\sum_{i} z_X(i)\,z_Y(i)=\hat{z}_X\cdot\hat{z}_Y, \qquad [1]$$

where $\underline{x}$, $s_X$, and $z_X$ are the (statistical) sample mean, sample standard deviation, and (unbiased) z-score vector for $X$ and $\underline{y}$, $s_Y$, and $z_Y$ are the corresponding statistics for $Y$, and the factor of $\frac{1}{n-1}$ adjusts for the difference in scale between $z$-score vectors computed with the unbiased sample standard deviation ($z_X$ and $z_Y$, which are not quite unit length vectors) and those computed with the biased sample standard deviation ($\hat{z}_X$ and $\hat{z}_Y$, which are). This dot product provides a geometric interpretation of Pearson's correlation coefficient, because the dot product of unit vectors is the cosine of the angle $A_{\hat{z}_X\hat{z}_Y}$ between them, the same angle $A_{XY}$ formed by points $X$, $Y$, and the $n$-dimensional origin:

$$\rho(X,Y)=\hat{z}_X\cdot\hat{z}_Y=\cos(A_{\hat{z}_X\hat{z}_Y})=\cos(A_{XY}).$$

### *Argument for a Beta null model for correlation coefficients*

The Quadrant Count Ratio (QCR)(Kader and Franklin 2008) is used as an educational stepping stone to motivate the formula for Pearson's Correlation Coefficient. For a set of $n$ pairs $\{(x_i, y_i) : 1 \le i \le n\}$, a vertical line at $\underline{x}$ and a horizontal line at $\underline{y}$ divide the pairs into four quadrants referred to as Quadrant I (containing the points with $x_i > \underline{x}$ and $y_i > \underline{y}$), Quadrant II ($x_i < \underline{x}$, $y_i > y$), Quadrant III (containing the points with $x_i < \underline{x}$ and $y_i < \underline{y}$), Quadrant IV ($x_i > \underline{x}$, $y_i < y$). Note that some points (the points for which $x_i = \underline{x}$ or $y_i = \underline{y}$) don't lie in any quadrant; for these points, the slope $m_i = \frac{y_i - \underline{y}}{x_i - \underline{x}}$ of the line through $(\underline{x}, \underline{y})$ and $(x_i, y_i)$ is either 0 or undefined. Letting $n_Q$ represent the number of points in Quadrant Q, the numbers of points corresponding to a line with $m_i > 0$ and $m_i < 0$ are $n_I + n_{III}$ and $n_{II} + n_{IV}$, respectively. If we score each point as "voting" for either positive correlation (+1) or negative correlation (-1) and divide by the total number of votes, we obtain the formula for the QCR:

$$QCR = \frac{(n_I + n_{III}) - (n_{II} + n_{IV})}{n_I + n_{II} + n_{III} + n_{IV}},$$

and if we define $f = \frac{QCR + 1}{2}$, we find that $f$ is the fraction of votes for positive correlation:

$$f = \frac{QCR + 1}{2} = \frac{1}{2}(QCR + 1)$$

$$¿\frac{1}{2}\left(\frac{(n_I + n_{III}) - (n_{II} + n_{IV})}{n_I + n_{II} + n_{III} + n_{IV}} + \frac{n_I + n_{II} + n_{III} + n_{IV}}{n_I + n_{II} + n_{III} + n_{IV}}\right)$$

$$¿\frac{1}{2}\left(\frac{2(n_I + n_{III})}{n_I + n_{II} + n_{III} + n_{IV}}\right)$$

$$f = \frac{n_I + n_{III}}{n_I + n_{II} + n_{III} + n_{IV}}.$$

If we consider $n^{¿}=n_I+n_{II}+n_{III}+n_{IV}$ votes for positive or negative correlation as a Binomial process with a success a vote for positive correlation, the vote result $V$ is a Binomial random variable with $n^{¿}$ trials and probability of success $f$, $V\sim Binomial(n^{¿},f)$. The distribution for estimating $f$ from a vote sample of size $n^{¿}$ (the sampling distribution for $f$) is known(Hastings et al. 2000) to be the Beta distribution, $f\sim Beta(v,\omega)$, where the shape parameters are one less than the number of successes and one less than the number of failures:

$$v=n_I+n_{III}-1=n^{¿}f-1$$
$$\omega=n_{II}+n_{IV}-1=n^{¿}(1-f)-1.$$

Thus, the Quadrant Count Ratio is distributed as a linear function of a Beta distribution,

$$QCR=2f-1\sim 2\,Beta(v,\omega)-1.$$

Returning to the Pearson correlation coefficient, we see the formula is a weighted version of the formula for the QCR; and separating the sum into positive and negative "votes":

$$\rho=\sum_{i}^{❑}\widehat{z_X}(i)\widehat{z_Y}(i)=\sum_{w_i>0}^{❑}|w_i|-\sum_{w_i<0}^{❑}|w_i|, \qquad [1]$$

where $w_i=\widehat{z_X}(i)\widehat{z_Y}(i)$. Since the parameters of the Beta distribution depend only on the vote totals and need not be integers, we deduce that

$$\rho\sim 2\,Beta(v,\omega)-1$$

For some shape parameters $v$ and $\omega$. In this case we are not trying to estimate $f$ from a fixed value for $\rho$; we are interested in identifying the family of distributions which can model an observed set of correlation coefficients, most of which are assumed to be from the null model.

***Weighted Correlation Network Analysis and higher-order information***

If $W$ is a third analyte measured on the same samples, the pairwise correlations $\rho(X, Y)$, $\rho(X, W)$, and $\rho(Y, W)$ are not mutually statistically independent, as the equations for $\rho(X, W)$, and $\rho(Y, W)$ constrain $\rho(X, Y)$ through $W$; and every additional measured analyte likewise introduces additional constraints. Exploiting such additional, higher-order constraints can be considered the motivation for constructing and analyzing an explicit model of the correlation network in WCNA. WCNA uses Topological Overlap Measure (TOM) to turn local information about the correlation structure of the network in the vicinity of a particular edge (analyte pair), which is then used in the clustering step. While TOM exploits the specific correlations between associated analytes–which presumably represent at least in part correlations created by a phenomenon of interest–we note that the Beta null model above is not influenced by the structure of the correlation network itself, it is purely a function of the number of pairs and the expected correlation coefficient (see [1] above). This too is higher-order information, but appears to provide qualitatively different higher-order information than the local information that TOM accesses. We therefore suggest TOM and the Beta null model are complementary approaches, and should provide the best results when used together in constructing the correlation network.
