## Supplement S2 for "Model-based Standardization of Correlation Coefficients Improves Multi-Omic Clustering and Biological Signal Discovery"

### Preliminaries

In [1]:
```
options(jupyter.plot_scale=1,
        width=200,
        repr.matrix.max.cols=200,
        repr.matrix.max.rows=Inf)
```

### Well-known t-test for non-zero correlation

The standard (two-sided) t-test of the null hypothesis $\rho = 0$ for $n$ dimensions with $E[\rho] = 0$ uses the ratio of the correlation coefficient with its standard error $\mathrm{SE}(\rho) = \sqrt{\frac{1-\rho^2}{n-2}}$, which has a $t$-distribution with $n-2$ degrees of freedom:

$$\frac{\rho\sqrt{n-2}}{\sqrt{1-\rho^2}} \sim t_{n-2}$$

$$\Pr(|\rho| > 0) = 2\Pr(t_{n-2} > \left|\frac{\rho\sqrt{n-2}}{\sqrt{1-\rho^2}}\right|)$$

In [2]:
```
correlation.t.test <- function(rho, n) {
  t <- abs(rho)*sqrt(n-2)/sqrt(1-rho*rho)
  return(2*pt(t,df=n-2,lower.tail=FALSE))
}
```

### Proposed Beta test for non-zero correlation

We have proposed a null model for correlation coefficients in the form of a Beta distribution,

$$\frac{(1+\rho)}{2} \sim \mathrm{Beta}(\nu, \omega),$$

which for $n$ independent and identically distributed bivariate Normal dimensions and $E[\rho] = 0$ has parameters $\nu = \omega = \frac{n}{2} - 1$. The corresponding two-sided significance test exploits the symmetry of the Beta distribution with equal shape parameters:

$$\begin{aligned} \Pr\Big(|\rho| > 0\Big) &= \Pr\Big(\big|\mathrm{Beta}(\nu,\nu)\big| > \frac{1+|\rho|}{2}\Big) && (1) \\ &= 2\Pr\Big(\mathrm{Beta}(\nu,\nu) > \frac{1+|\rho|}{2}\Big). && (2) \end{aligned}$$

In [3]:
```
correlation.beta.test <- function(rho, n) {
  v <- n/2-1
  return(2*pbeta((1+abs(rho))/2,v,v,lower.tail=FALSE))
}
```

In [ ]:
```
#
# m = (u-1) / (u+v-2)
# k = u + v = n - 2
#
# u = 1 +     m * (k-2)
# v = 1 + (1-m) * (k-2)
```

In [411…
```
###
# when u = v = n/2-1, u + v = 2(n/2 - 1) = n-2.
###
noncentral.correlation.beta.test <- function(rho, u, v) {
  x <- (1.0+rho)/2.0
  p.lo <- pbeta(x,u,v,lower.tail=TRUE)
  p.hi <- pbeta(x,u,v,lower.tail=FALSE)
  return(2*pmin(p.lo,p.hi))
}
```

In [412…
```
correlation.Fisher.test <- function(rho, rho0, n) {
  z <- (atanh(rho) - atanh(rho0))*sqrt(n-3)
  return(2*pnorm(abs(z),lower.tail=FALSE))
}
```

In [316…
```
fine <- 500
rho <- c(-fine:fine)/fine ; rho <- ifelse(rho > -1, ifelse(rho < 1, rho,
```

### Equivalence of the $t$ and Beta tests when $\mathrm{E}[\rho] = 0$

Since the Beta null model is new, we demonstrate here for the case $\mathrm{E}[\rho] = 0$ that the two-sided significance test against the Beta null model is identical to the standard t-test for this case for $n \geq 3$ measured analytes (equivalently, paired samples), with numerical differences between the computed p-values on the order of floating point errors.

In [428…
```
par(mfrow=c(2,2))
for (n in c(3:6)) {
  plot(rho,correlation.t.test(rho,n),type='l',xlab="Correlation",
       ylab="p-value",main=paste("n =",n),lwd=3,col='dodgerblue')
  abline(h=c(0:2)/2,lty=3,col='gray')
  abline(v=c(-2:2)/2,lty=3,col='gray')
  lines(rho,correlation.beta.test(rho,n),lwd=1, col='black')
}
legend(-1,1,legend=c("t-test","Beta"),lwd=c(1,3),col=c('black','dodgerblu
```

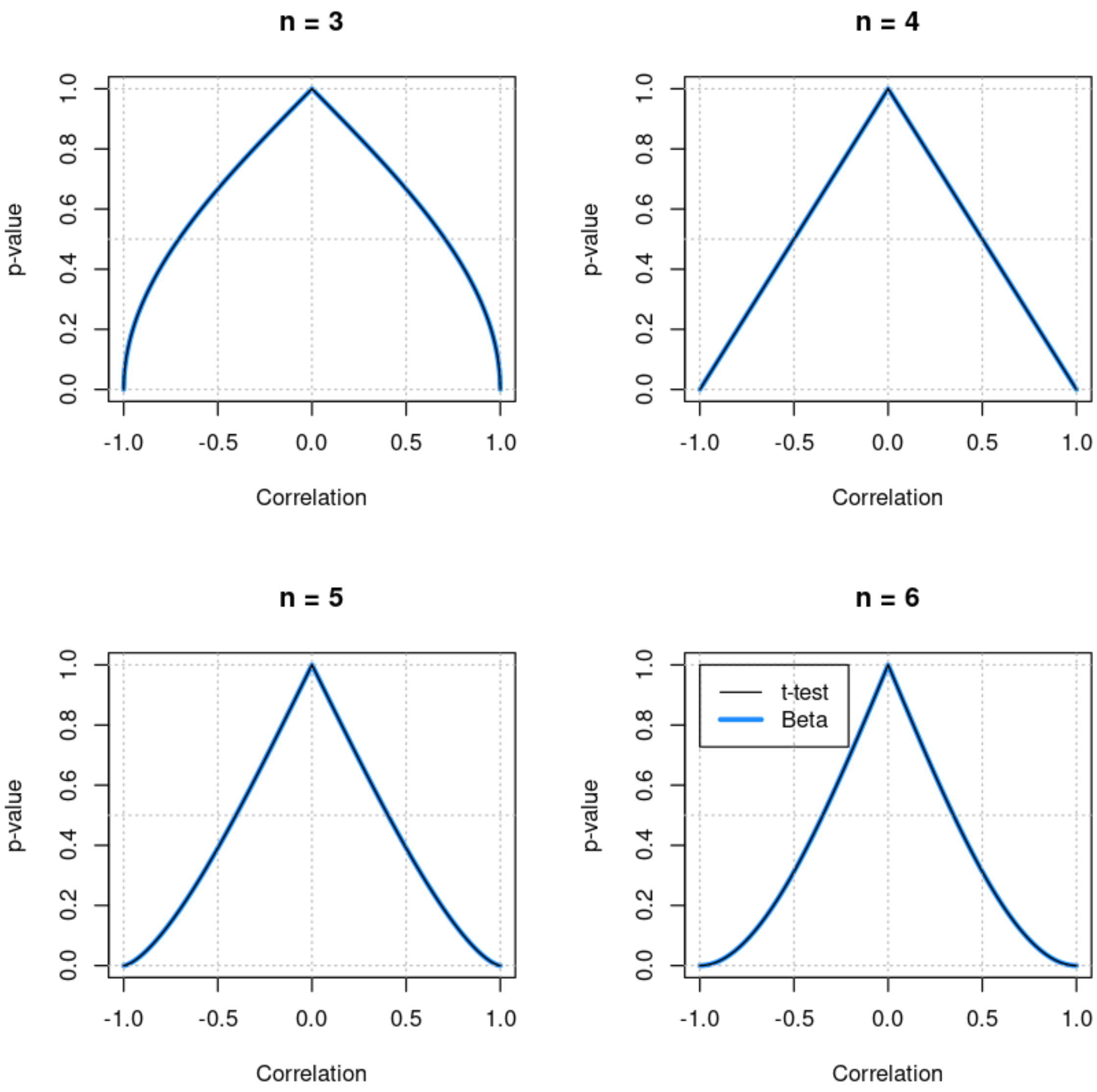


In [434…

```r
par(mfrow=c(2,2))
for (n in c(3:6)) {
  dp <- correlation.t.test(rho,n)-correlation.beta.test(rho,n)
  plot(rho, dp*10^15, type='l',xlab="Correlation", ylim=c(-5,5),
       ylab="p_t - p_beta (x 10^-15)",main=paste("n =",n),pch=1)
  abline(h=0,col='gray')
}
```

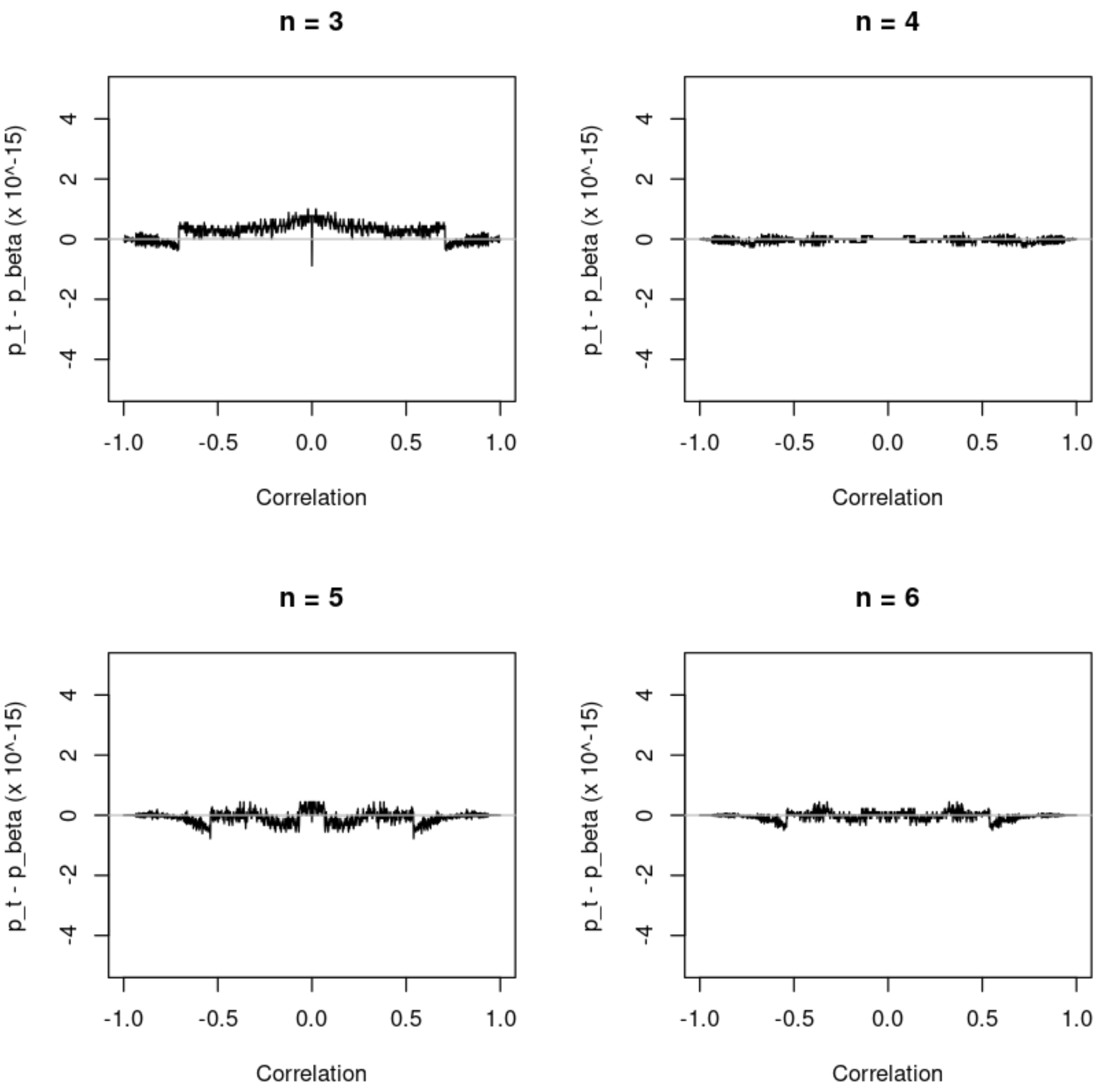


In [430…

```
par(mfrow=c(2,2))
for (n in c(10,20,25,30)) {
  plot(rho,correlation.t.test(rho,n),type='l',xlab="Correlation",
       ylab="p-value",main=paste("n =",n),lwd=3,col='dodgerblue')
  abline(h=c(0:2)/2,lty=3,col='gray')
  abline(v=c(-2:2)/2,lty=3,col='gray')
  lines(rho,correlation.beta.test(rho,n),lwd=1, col='black')
}
legend(-1,1,legend=c("t-test","Beta"),lwd=c(1,3),col=c('black','dodgerblu
```

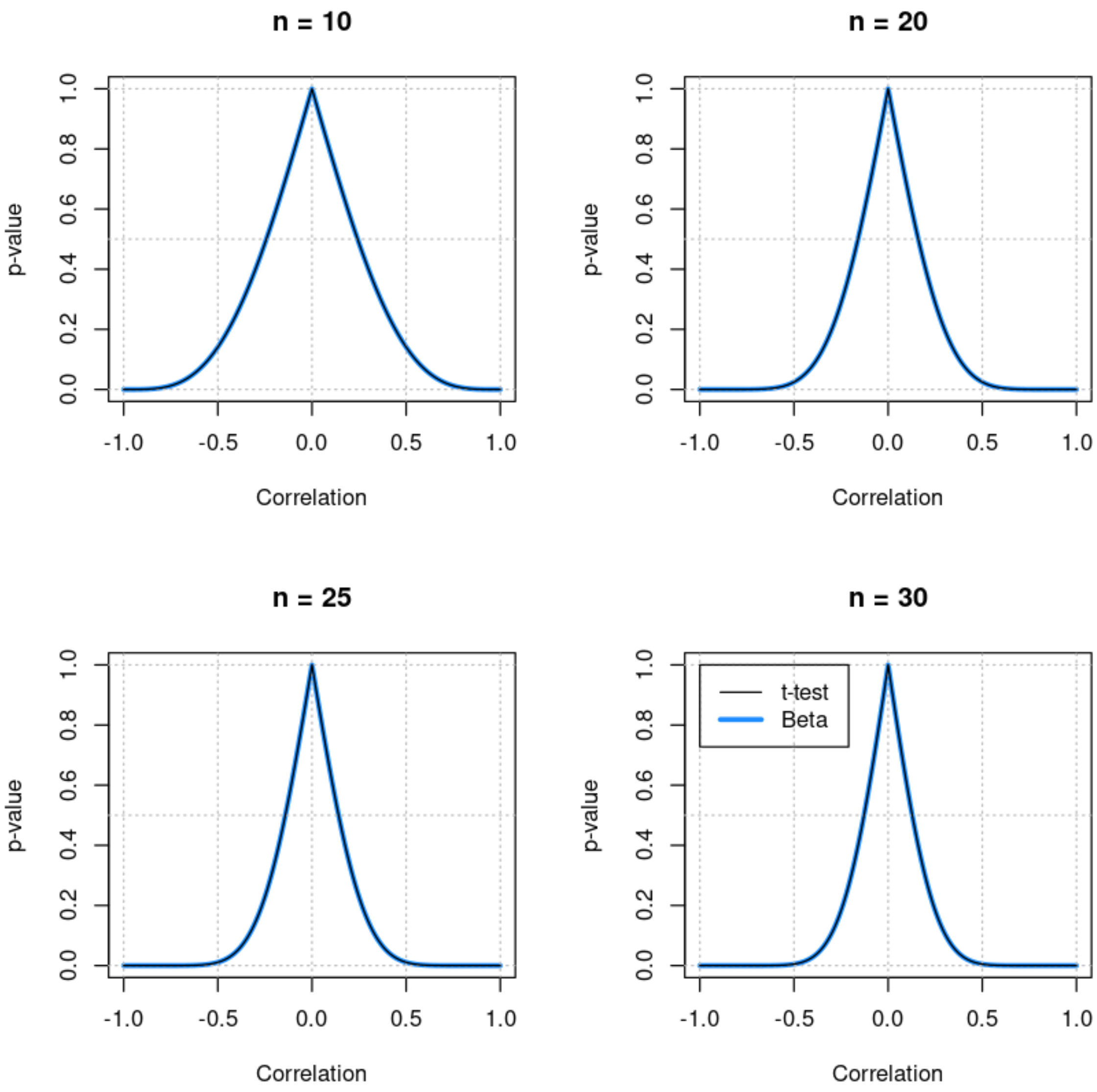


In [433…

```
par(mfrow=c(2,2))
for (n in c(10,20,25,30)) {
  dp <- correlation.t.test(rho,n)-correlation.beta.test(rho,n)
  plot(rho, dp*10^15, type='l',xlab="Correlation", ylim=c(-5,5),
       ylab="p_t - p_beta (x 10^-15)",main=paste("n =",n),pch=1)
  abline(h=0,col='gray')
}
```

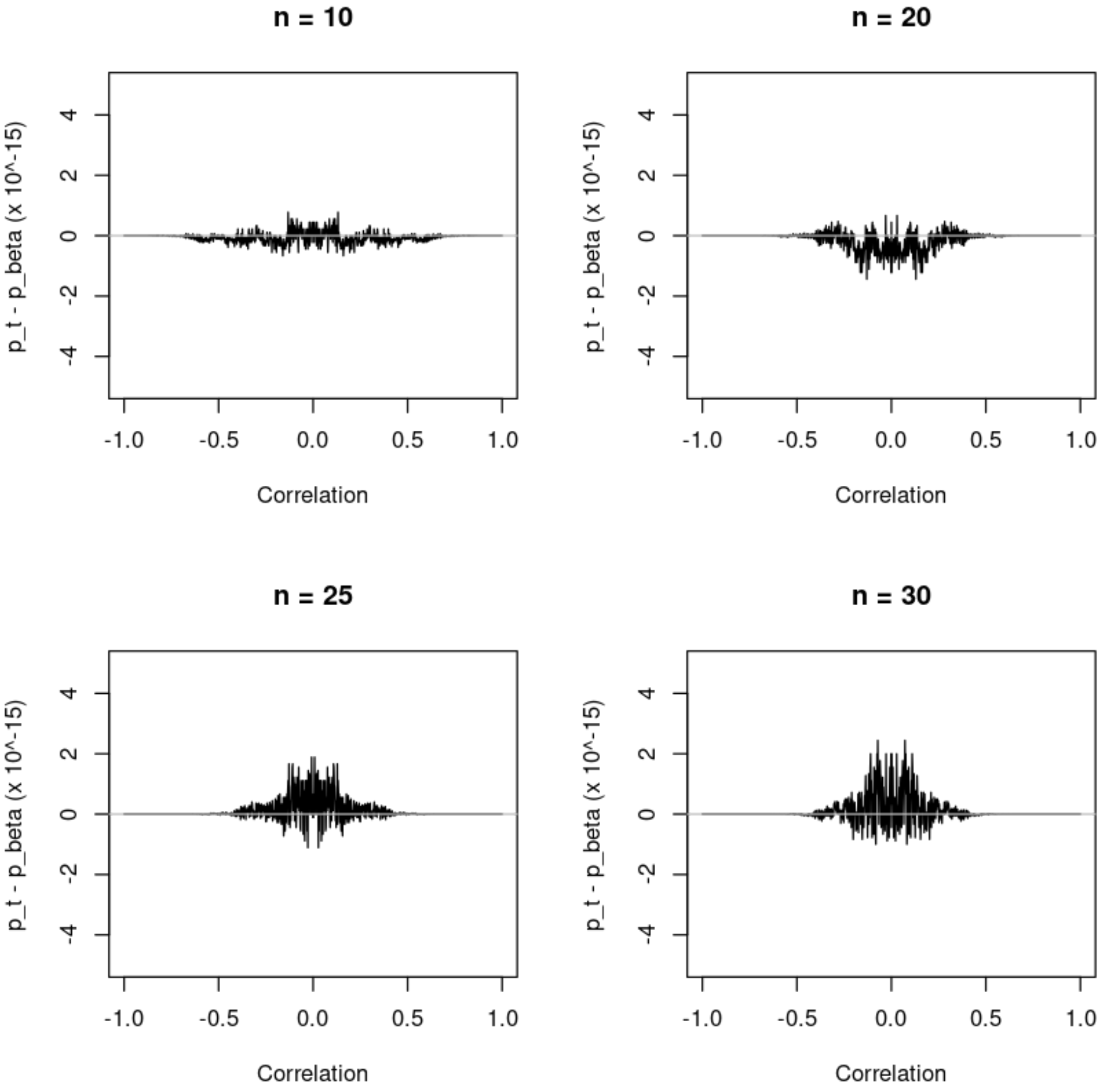


In [435…

```
par(mfrow=c(2,2))
for (n in c(50,100,200,1000)) {
  plot(rho,correlation.t.test(rho,n),type='l',xlab="Correlation",
       ylab="p-value",main=paste("n =",n),lwd=3,col='dodgerblue')
  abline(h=c(0:2)/2,lty=3,col='gray')
  abline(v=c(-2:2)/2,lty=3,col='gray')
  lines(rho,correlation.beta.test(rho,n),lwd=1, col='black')
}
legend(-1,1,legend=c("t-test","Beta"),lwd=c(1,3),col=c('black','dodgerblu
```

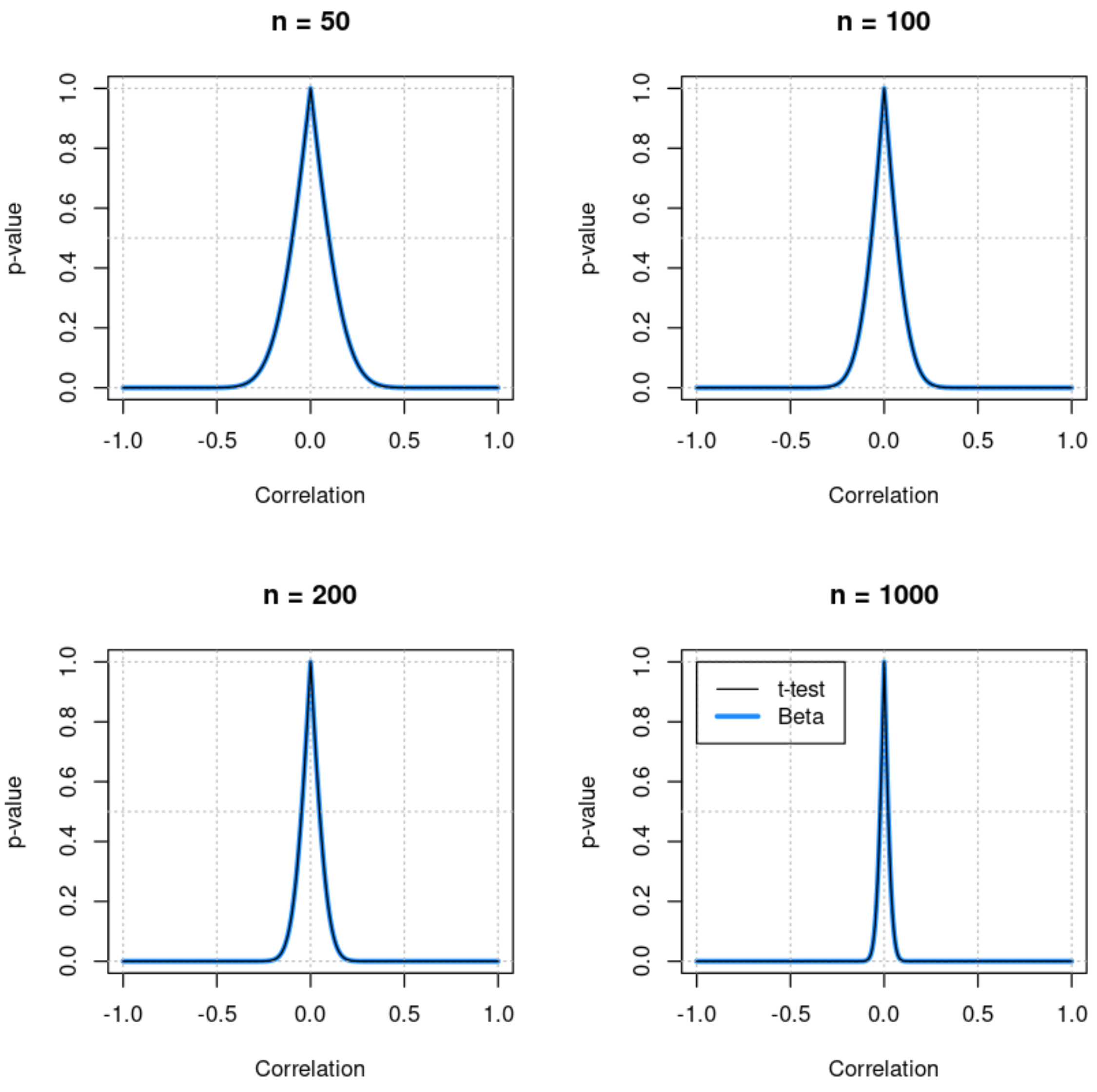


In [436…

```
par(mfrow=c(2,2))
for (n in c(50,100,200,1000)) {
  dp <- correlation.t.test(rho,n)-correlation.beta.test(rho,n)
  plot(rho, dp*10^15, type='l',xlab="Correlation", ylim=c(-5,5),
       ylab="p_t - p_beta (x 10^-15)",main=paste("n =",n),pch=1)
  abline(h=0,col='gray')
}
```

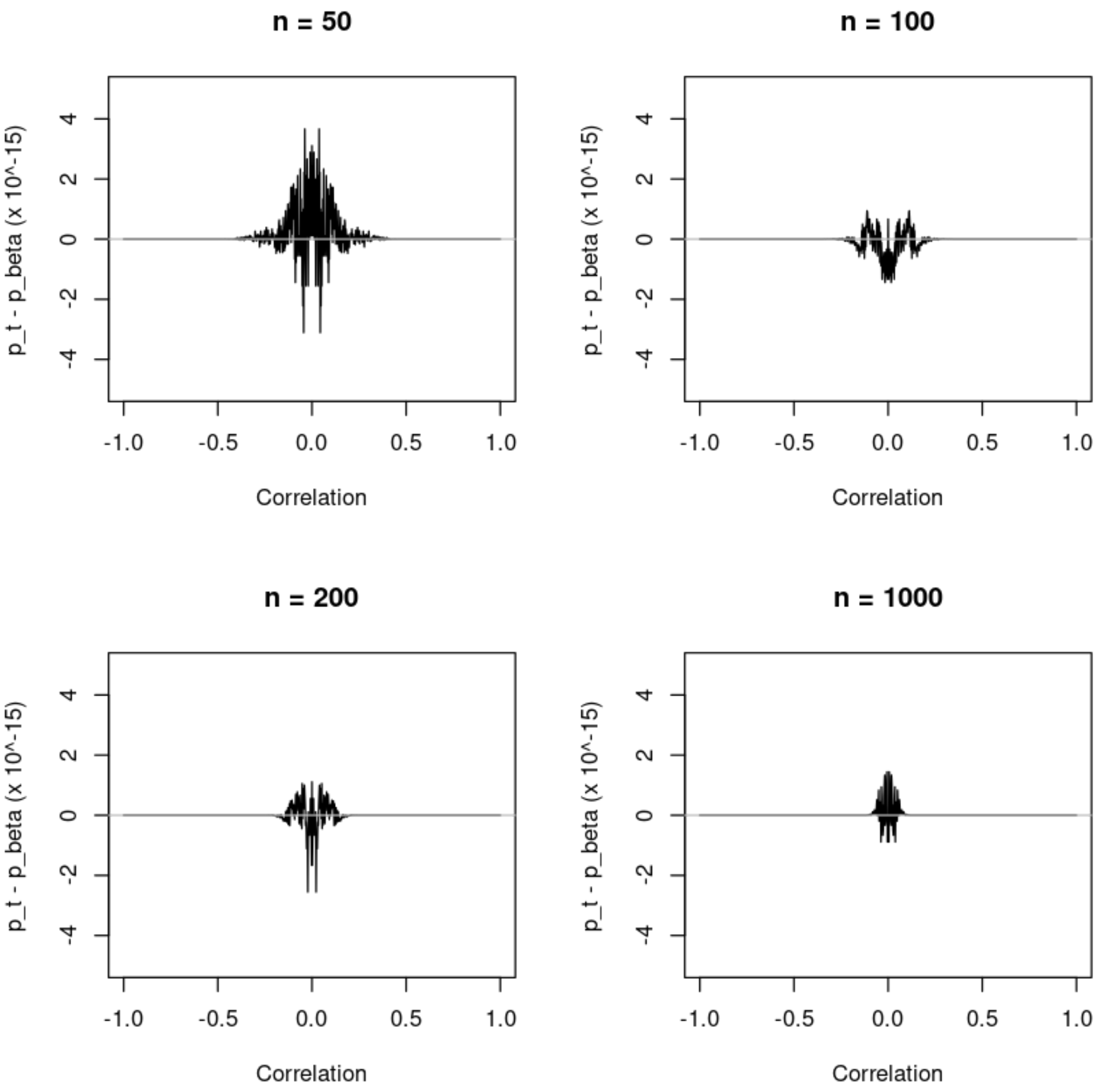


### Comparison with Fisher transformation, $\mathrm{E}[\rho] = 0$

The Fisher transformation provides an approximate $z$-score, $z = F(\rho)$, which is has a Normal distribution. Here we show how similar this approach is in the same special case considered above.

In [401…

```
par(mfrow=c(2,2))
for (n in c(4:7)) {
  plot(rho,correlation.beta.test(rho,n),type='l',xlab="Correlation (R)",y
       main=paste("n =",n),lwd=3,col="dodgerblue")
  abline(h=c(0:2)/2,lty=3,col='gray')
  abline(v=c(-2:2)/2,lty=3,col='gray')
  lines(rho,correlation.Fisher.test(rho,0,n),lwd=1)
}
legend(-1,1,legend=c("Beta","Fisher"),lwd=c(3,1),col=c('dodgerblue','blac
```

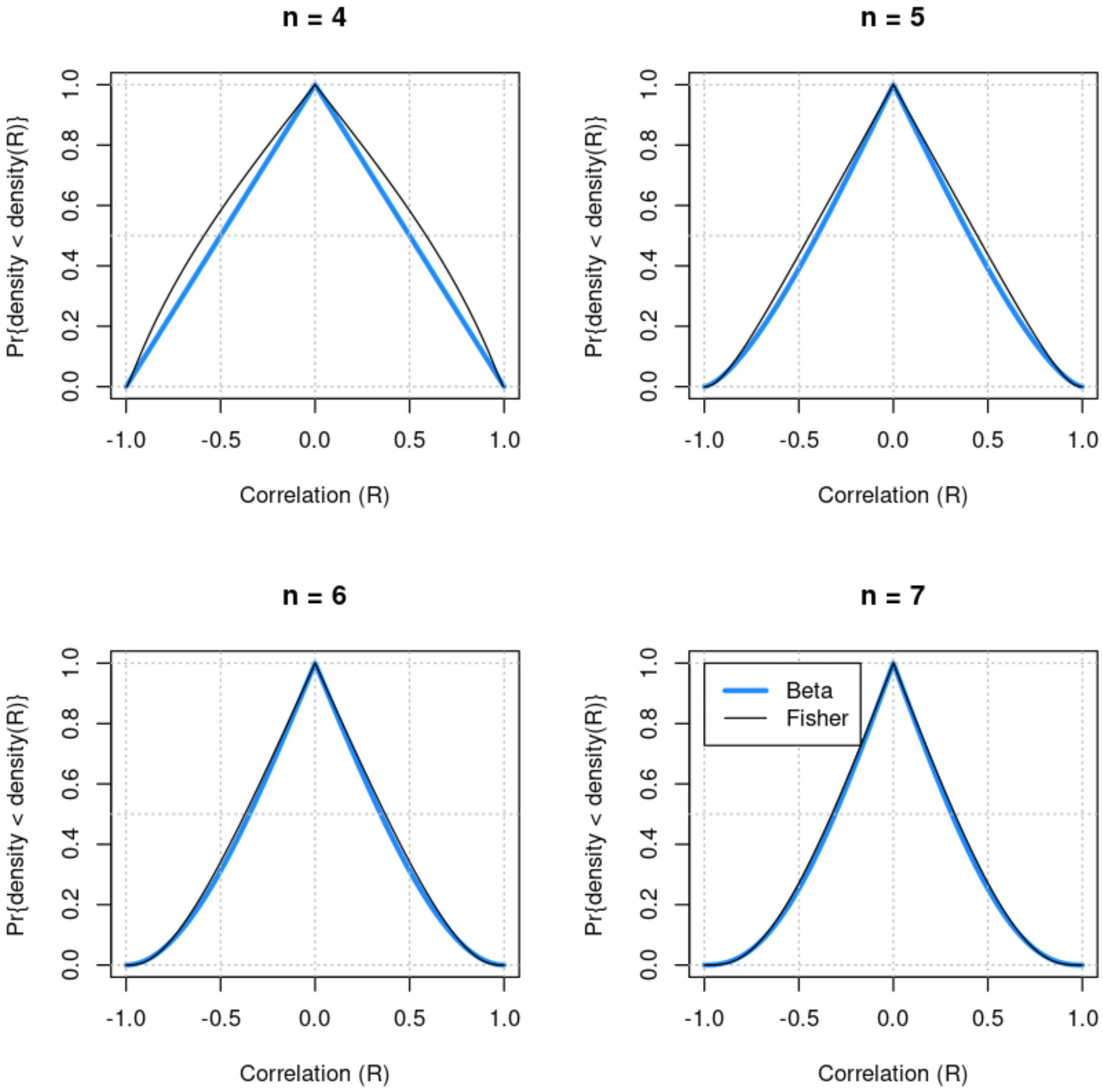


In [402…

```
par(mfrow=c(2,2))
for (n in c(10,15,20,30)) {
  plot(rho,correlation.beta.test(rho,n),type='l',xlab="Correlation",ylab=
       main=paste("n =",n),lwd=3,col="dodgerblue")
  abline(h=c(0:2)/2,lty=3,col='gray')
  abline(v=c(-2:2)/2,lty=3,col='gray')
  lines(rho,correlation.Fisher.test(rho,0,n),lwd=1)
}
legend(-1,1,legend=c("Beta","Fisher"),lwd=c(3,1),col=c('dodgerblue','blac
```

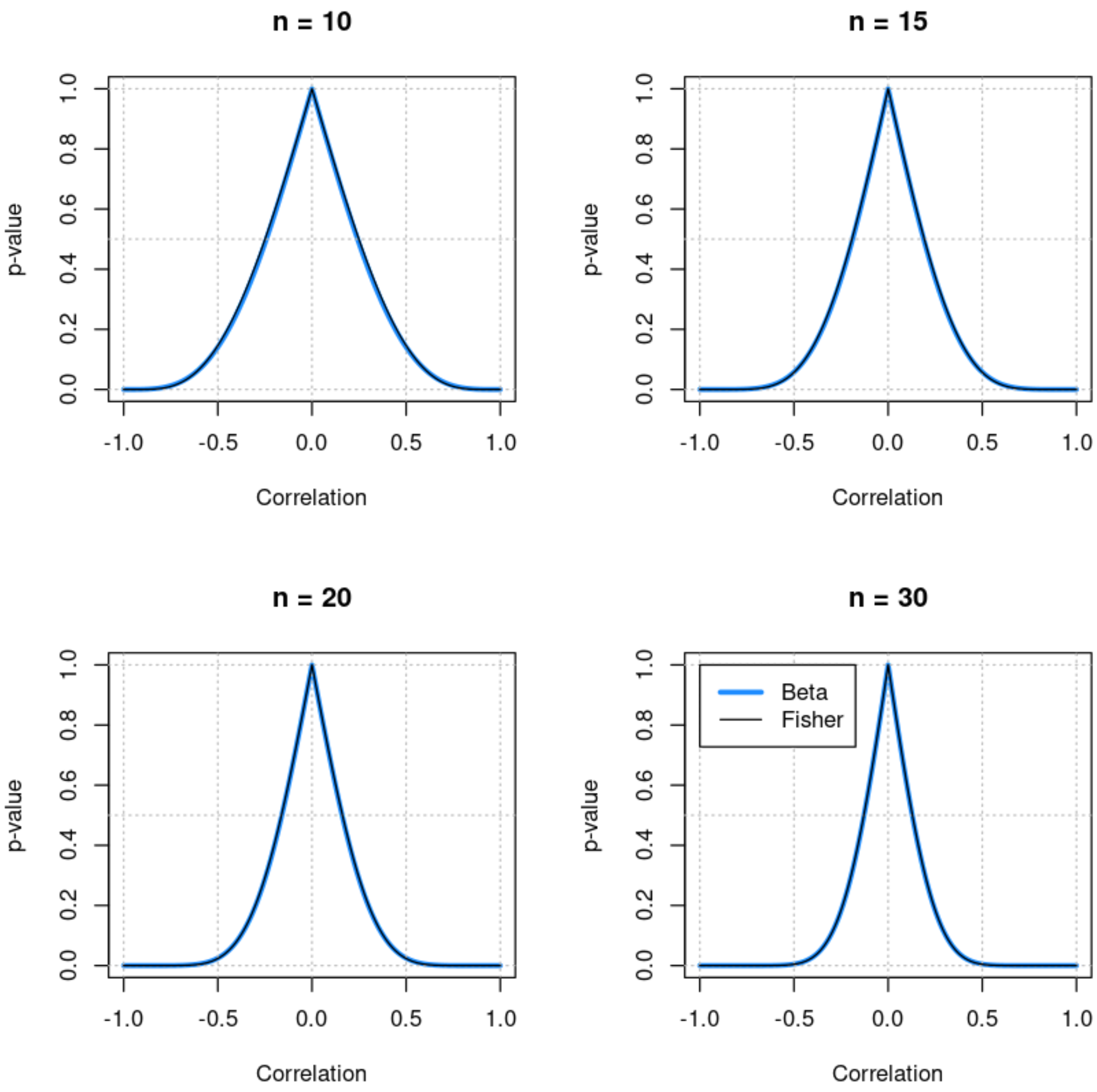


### Comparison with Fisher transformation, $-0.9 \leq E[\rho] \leq 0.9$

The Fisher transformation is the standard approach when $\mathrm{E} \neq 0$, and here we consider how the Beta null model compares in this more general case, and how the correspondence to the Fisher transformation changes as the sample size decreases.

### # u = v = n/2 - 1 # n=3 -> u = v = 1/2 # n=4 -> u = v = 1 # n >= 5: # mean.beta = u / (u+v) # u = mean.beta (n+2) # v = 2 + n - u # n = u + v - 2 # # if (n >= 5) { # u <- 1 + (n-4)*mean.beta # v <- 1 + (n-4)*(1-mean.beta) # } else { # u <- (n-2)*(1+mean.beta)/3 # v <- (n-2)*(2-mean.beta)/3 # } # rho2 <- rho[abs(rho) + 1e-7 < 1] f <- dbeta((1+rho2)/2,u,v) f <- f / max(f) lines(rho2,f,col='orangered')

In [500…

```
par(mfcol=c(2,2))
n <- 1000
min.logp <- -6
for (rho0 in c(-0.6,-0.1,0.3,0.9)) {
  mean.beta <- (1 + rho0)/2
  u <- (n - 2) *      mean.beta
  v <- (n - 2) * (1 - mean.beta)
  beta.p <- noncentral.correlation.beta.test(rho, u, v)
  f <- dbeta((1+rho)/2,u,v)
```

```
  plot(rho, f, type='l', lwd=3, xlab="Correlation (R)", ylab="Density", c
       main=c(paste("R ~ Beta(",round(u,2),",",round(v,2),")",sep=''),
              paste("n =",n,"    E[R] =",round(rho0,3))))
  abline(h=c(0,1), lty=3, col='grey')
  abline(v=c(-2:2)/2, lty=3, col='grey')
  abline(v=rho0, lty=2, lwd=3, col='grey60')

  suppressWarnings(plot(rho, log10(beta.p), type='l', lwd=3, xlab="Correl
                        ylim=c(min.logp,0), col='dodgerblue', main="Pr{de
  abline(h=c(min.logp:0), lty=3, col='grey')
  abline(v=c(-2:2)/2, lty=3, col='grey')
  abline(v=rho0, lty=2, lwd=3, col='grey60')
  suppressWarnings(lines(rho,log10(correlation.Fisher.test(rho,rho0,n)),l
}
legend(-1,0,legend=c("Fisher","Beta"),lwd=c(1,3),col=c('black','dodgerblu
```

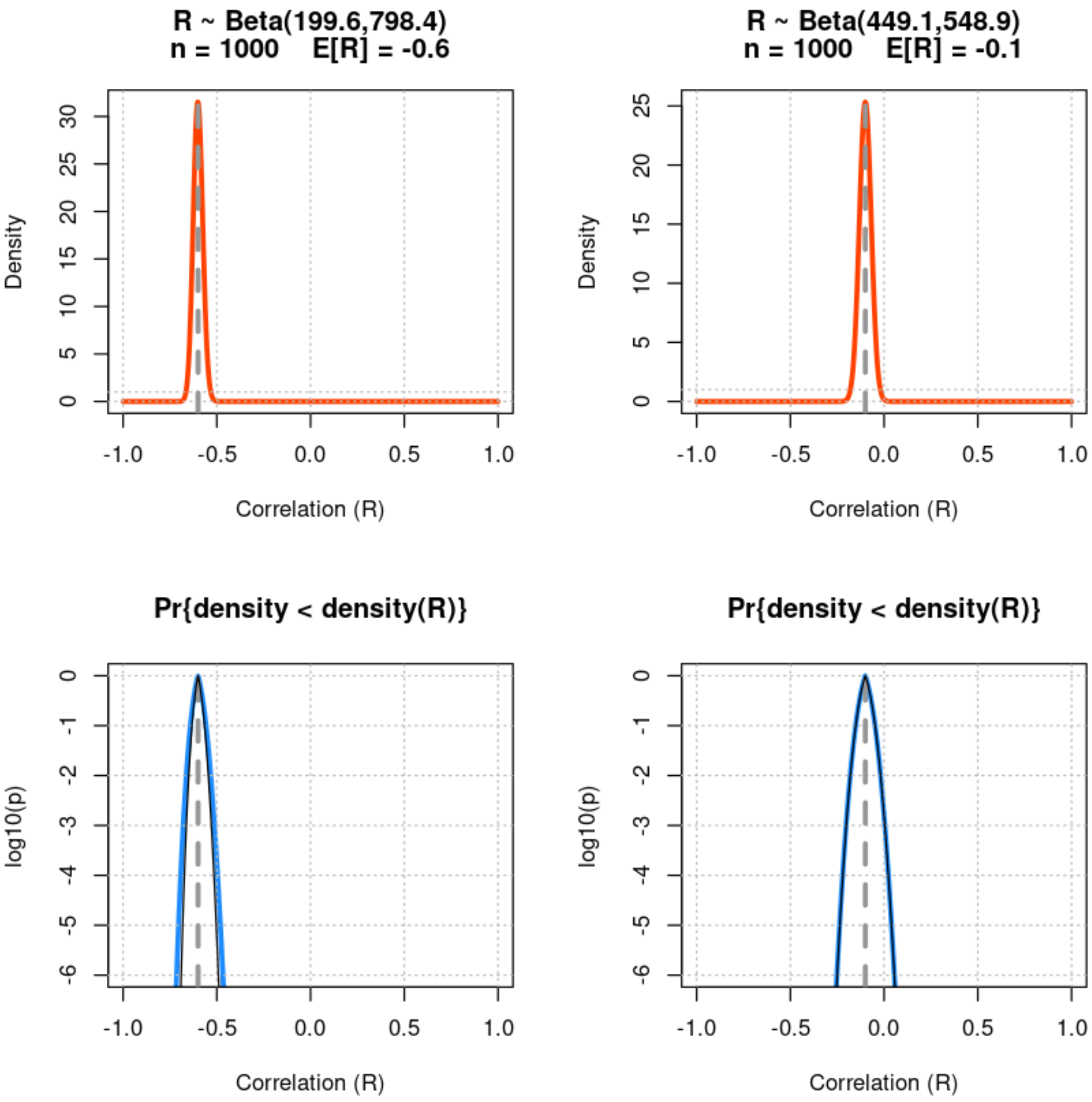

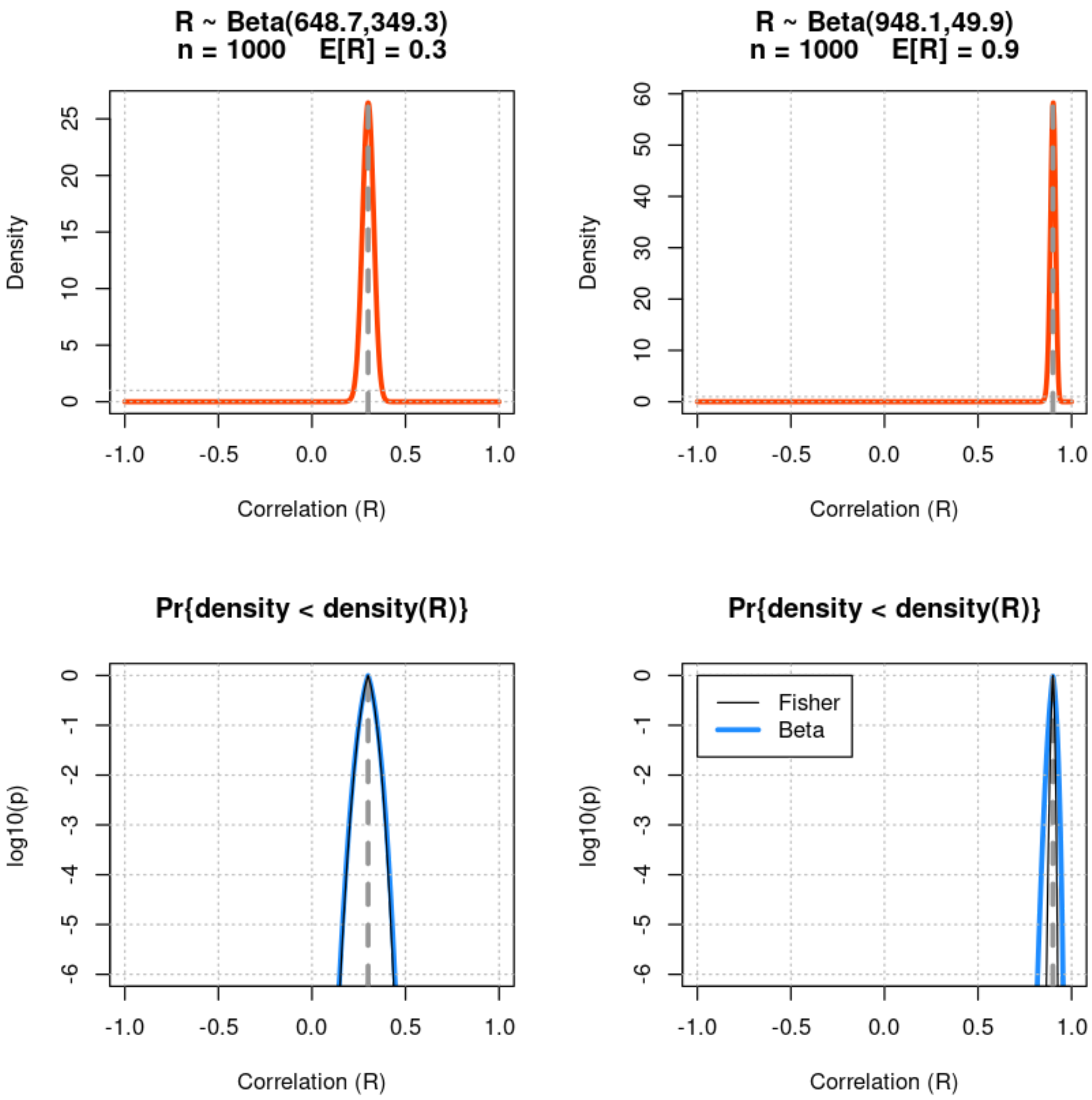


In [502…

```
par(mfcol=c(2,2))
n <- 100
min.logp <- -6
for (rho0 in c(-0.6,-0.1,0.3,0.9)) {
  mean.beta <- (1 + rho0)/2
  u <- (n - 2) *      mean.beta
  v <- (n - 2) * (1 - mean.beta)
  beta.p <- noncentral.correlation.beta.test(rho, u, v)
  f <- dbeta((1+rho)/2,u,v)
  plot(rho, f, type='l', lwd=3, xlab="Correlation (R)", ylab="Density", c
       main=c(paste("R ~ Beta(",round(u,2),",",round(v,2),")",sep=''),
              paste("n =",n,"    E[R] =",round(rho0,3))))
  abline(h=c(0,1), lty=3, col='grey')
  abline(v=c(-2:2)/2, lty=3, col='grey')
  abline(v=rho0, lty=2, lwd=3, col='grey60')

  suppressWarnings(plot(rho, log10(beta.p), type='l', lwd=3, xlab="Correl
                        ylim=c(min.logp,0), col='dodgerblue', main="Pr{de
  abline(h=c(min.logp:0), lty=3, col='grey')
  abline(v=c(-2:2)/2, lty=3, col='grey')
  abline(v=rho0, lty=2, lwd=3, col='grey60')
  suppressWarnings(lines(rho,log10(correlation.Fisher.test(rho,rho0,n)),l
}
legend(-1,0,legend=c("Fisher","Beta"),lwd=c(1,3),col=c('black','dodgerblu
```

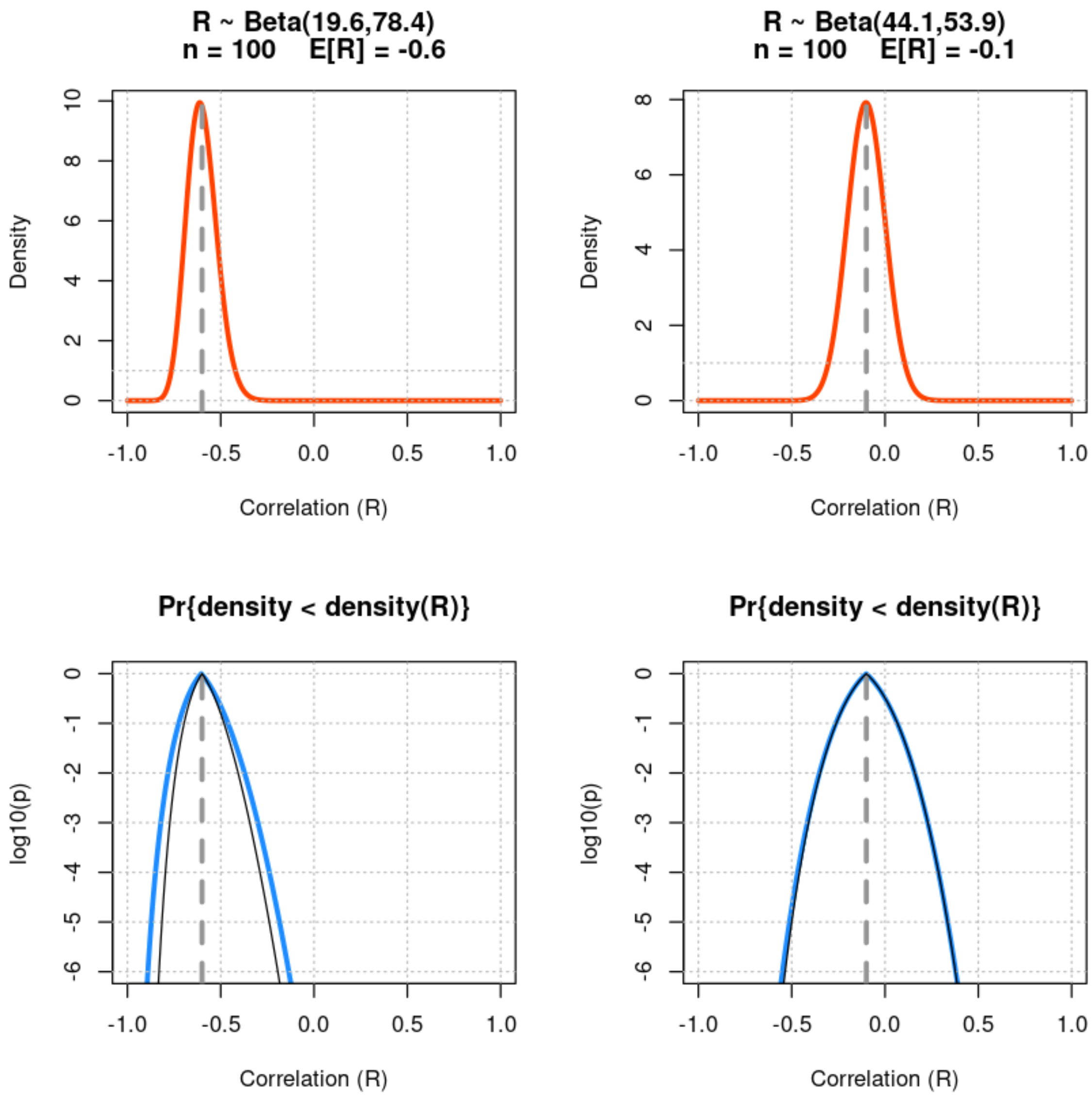
R ~ Beta(19.6,78.4)
n = 100 E[R] = -0.6
R ~ Beta(44.1,53.9)
n = 100 E[R] = -0.1
Density
Correlation (R)
Pr{density < density(R)}
log10(p)

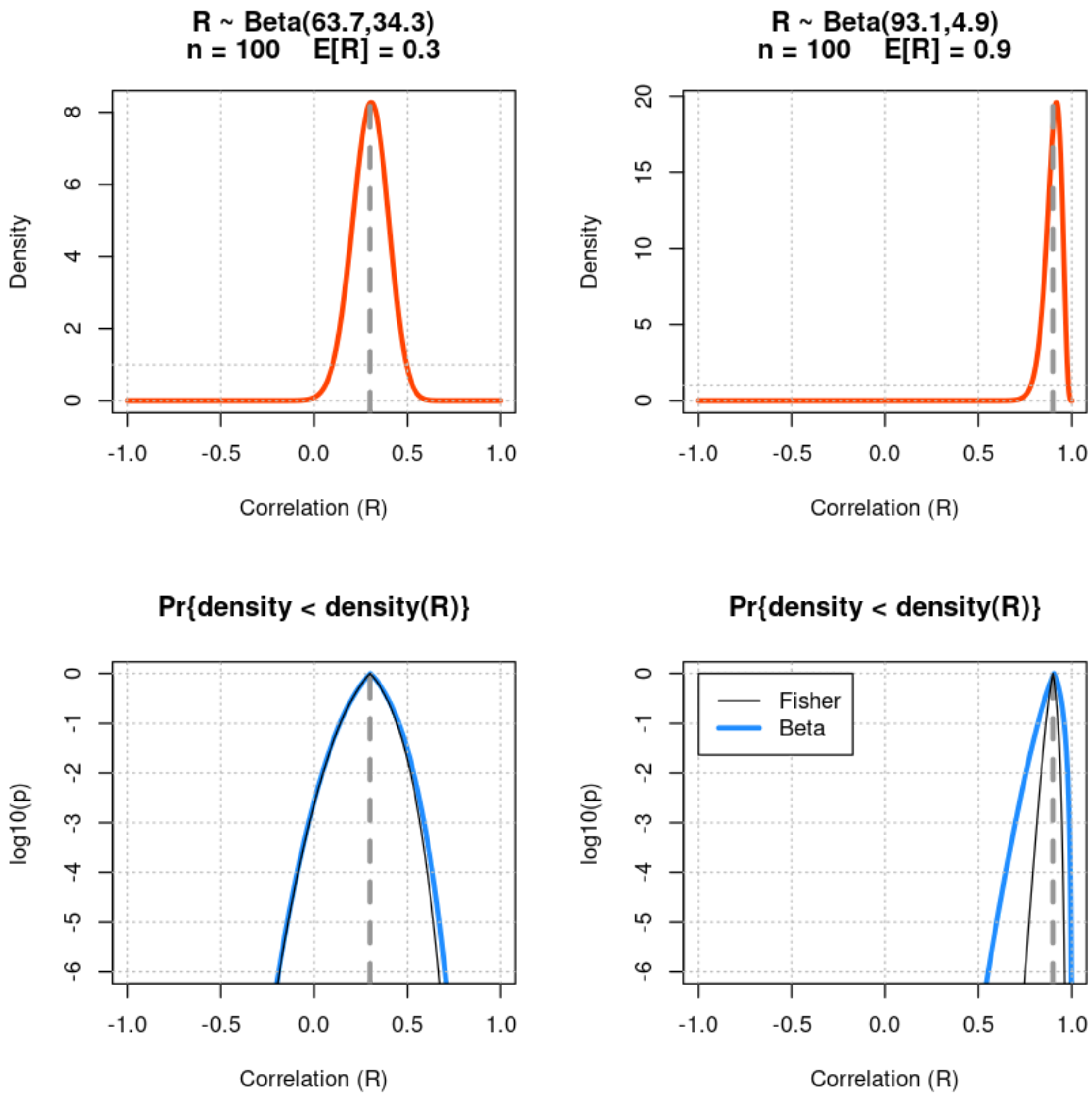


In [503…

```
par(mfcol=c(2,2))
n <- 30
min.logp <- -6
for (rho0 in c(-0.6,-0.1,0.3,0.9)) {
  mean.beta <- (1 + rho0)/2
  u <- (n - 2) *      mean.beta
  v <- (n - 2) * (1 - mean.beta)
  beta.p <- noncentral.correlation.beta.test(rho, u, v)
  f <- dbeta((1+rho)/2,u,v)
  plot(rho, f, type='l', lwd=3, xlab="Correlation (R)", ylab="Density", c
       main=c(paste("R ~ Beta(",round(u,2),",",round(v,2),")",sep=''),
              paste("n =",n,"   E[R] =",round(rho0,3))))
  abline(h=c(0,1), lty=3, col='grey')
  abline(v=c(-2:2)/2, lty=3, col='grey')
  abline(v=rho0, lty=2, lwd=3, col='grey60')

  suppressWarnings(plot(rho, log10(beta.p), type='l', lwd=3, xlab="Correl
                        ylim=c(min.logp,0), col='dodgerblue', main="Pr{de
  abline(h=c(min.logp:0), lty=3, col='grey')
  abline(v=c(-2:2)/2, lty=3, col='grey')
  abline(v=rho0, lty=2, lwd=3, col='grey60')
  suppressWarnings(lines(rho,log10(correlation.Fisher.test(rho,rho0,n)),l
}
legend(-1,0,legend=c("Fisher","Beta"),lwd=c(1,3),col=c('black','dodgerblu
```

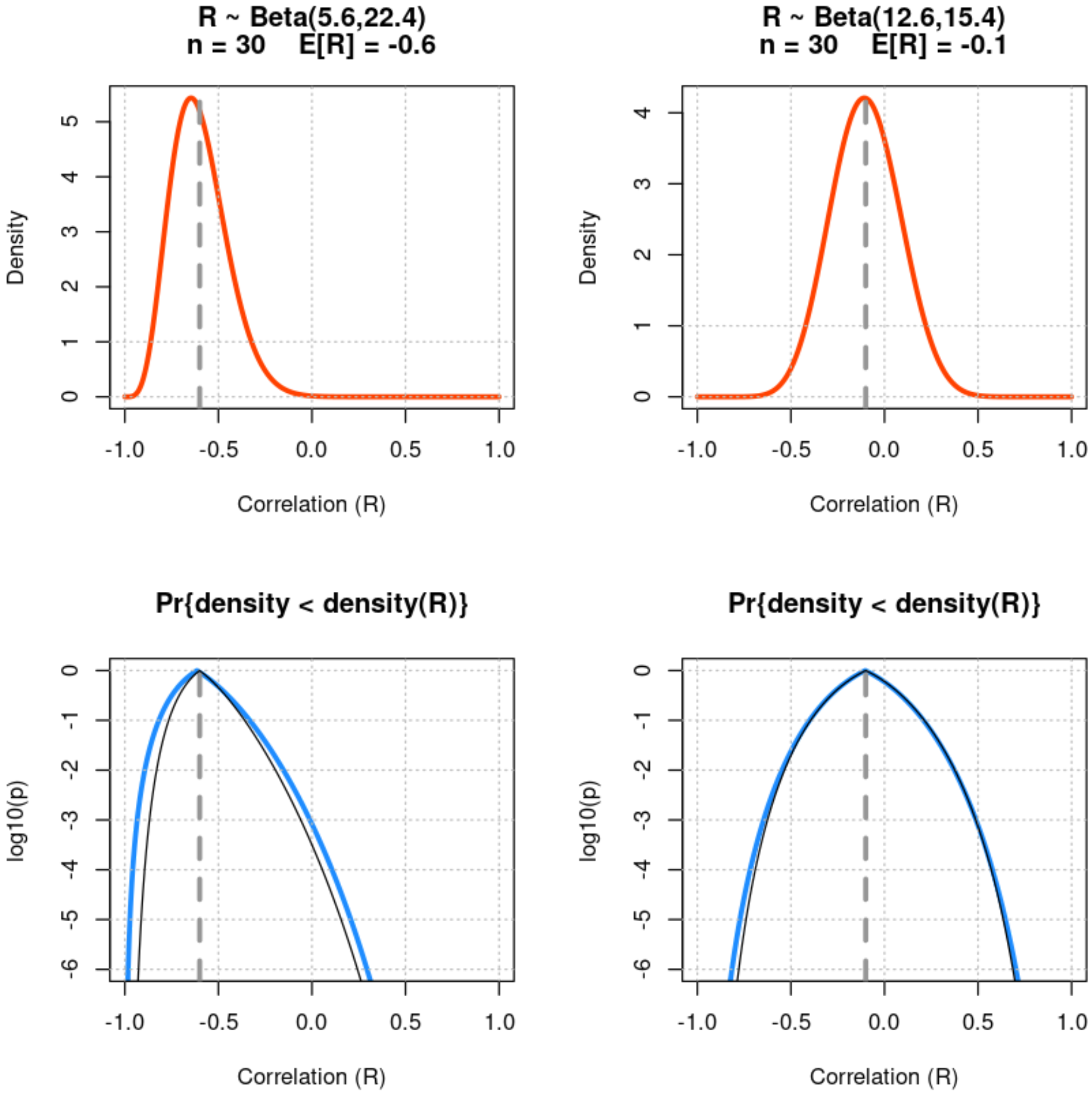

R ~ Beta(5.6,22.4)
n = 30    E[R] = -0.6
Density
Correlation (R)
R ~ Beta(12.6,15.4)
n = 30    E[R] = -0.1
Density
Correlation (R)
Pr{density < density(R)}
log10(p)
Correlation (R)
Pr{density < density(R)}
log10(p)
Correlation (R)

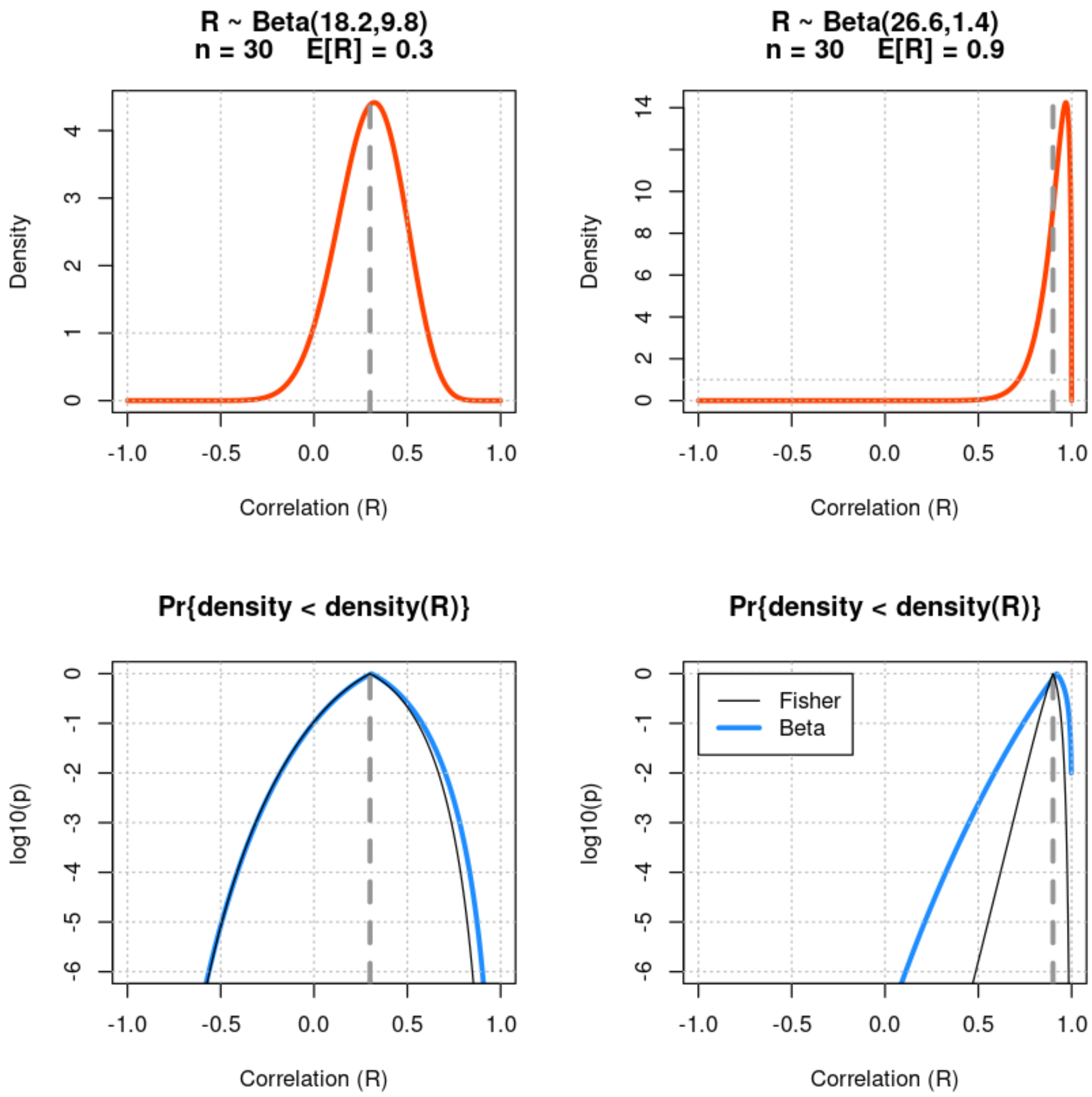


In [504…

```
par(mfcol=c(2,2))
n <- 20
min.logp <- -6
for (rho0 in c(-0.6,-0.1,0.3,0.9)) {
  mean.beta <- (1 + rho0)/2
  u <- (n - 2) *       mean.beta
  v <- (n - 2) * (1 - mean.beta)
  beta.p <- noncentral.correlation.beta.test(rho, u, v)
  f <- dbeta((1+rho)/2,u,v)
  plot(rho, f, type='l', lwd=3, xlab="Correlation (R)", ylab="Density", c
       main=c(paste("R ~ Beta(",round(u,2),",",round(v,2),")",sep=''),
              paste("n =",n,"    E[R] =",round(rho0,3))))
  abline(h=c(0,1), lty=3, col='grey')
  abline(v=c(-2:2)/2, lty=3, col='grey')
  abline(v=rho0, lty=2, lwd=3, col='grey60')

  suppressWarnings(plot(rho, log10(beta.p), type='l', lwd=3, xlab="Correl
                        ylim=c(min.logp,0), col='dodgerblue', main="Pr{de
  abline(h=c(min.logp:0), lty=3, col='grey')
  abline(v=c(-2:2)/2, lty=3, col='grey')
  abline(v=rho0, lty=2, lwd=3, col='grey60')
  suppressWarnings(lines(rho,log10(correlation.Fisher.test(rho,rho0,n)),l
}
legend(-1,0,legend=c("Fisher","Beta"),lwd=c(1,3),col=c('black','dodgerblu
```

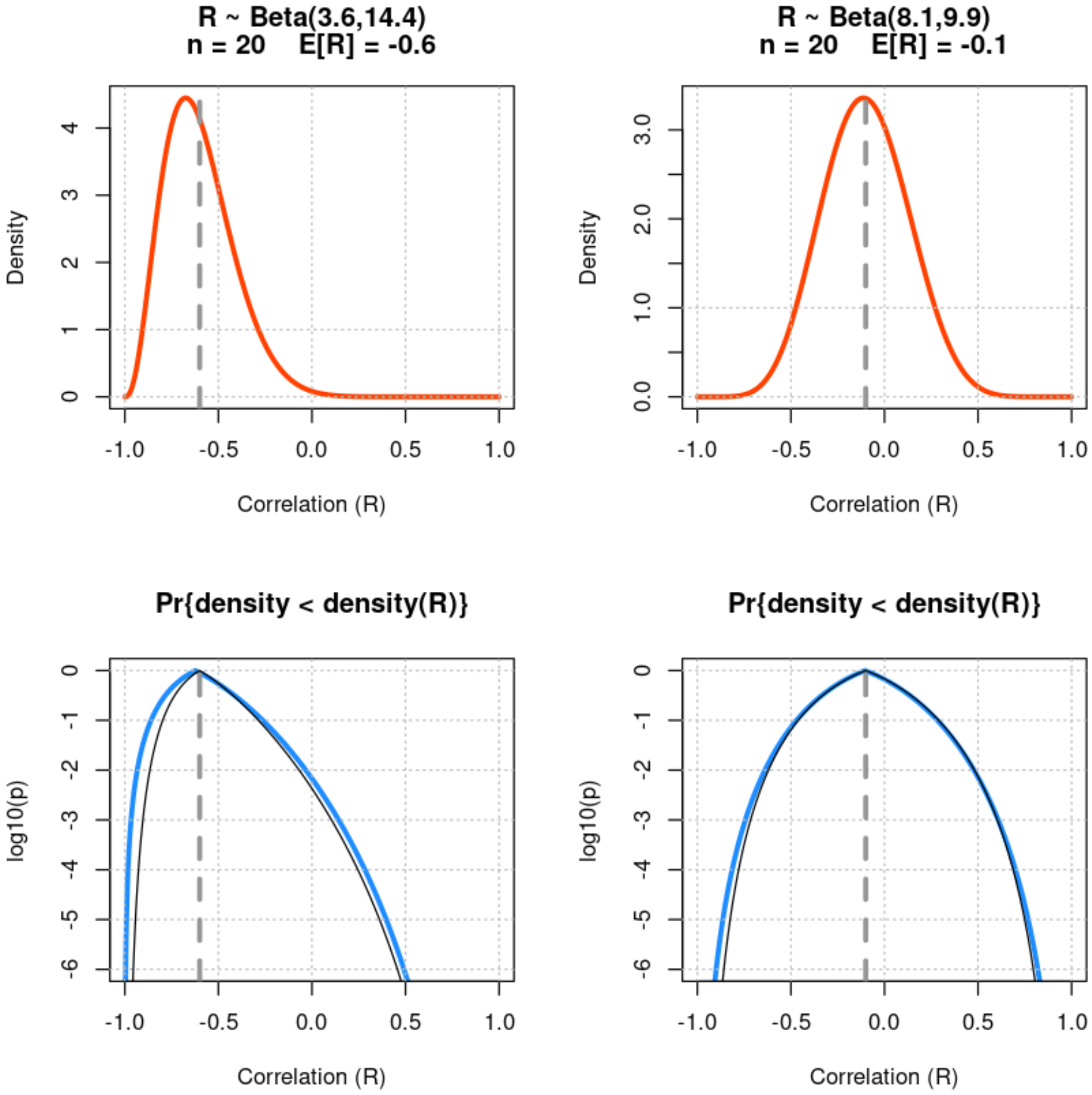

R ~ Beta(3.6,14.4)
n = 20 E[R] = -0.6
Density
Correlation (R)
R ~ Beta(8.1,9.9)
n = 20 E[R] = -0.1
Density
Correlation (R)
Pr{density < density(R)}
log10(p)
Correlation (R)
Pr{density < density(R)}
log10(p)
Correlation (R)

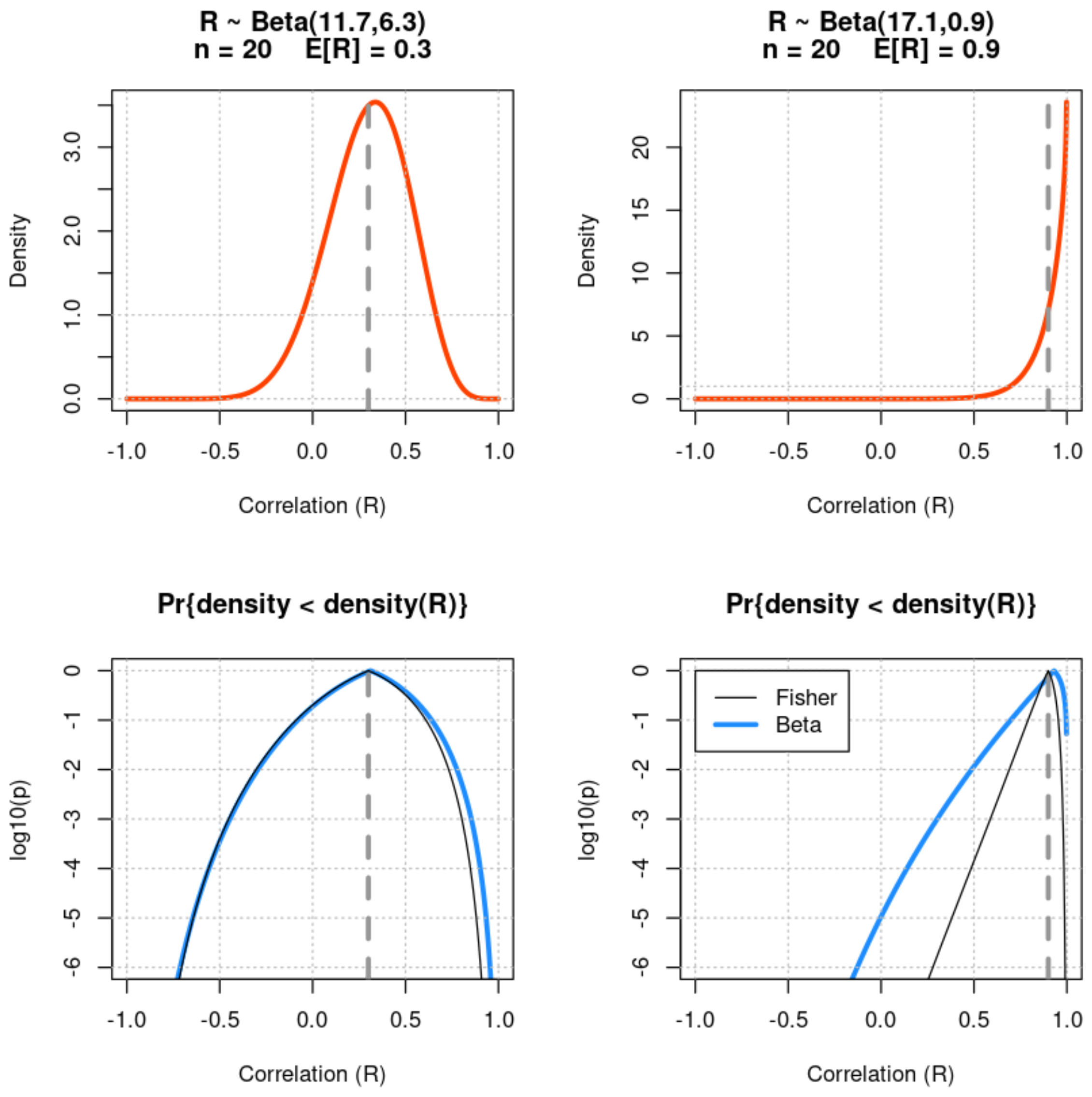


In [505…

```
par(mfcol=c(2,2))
n <- 10
min.logp <- -6
for (rho0 in c(-0.6,-0.1,0.3,0.9)) {
  mean.beta <- (1 + rho0)/2
  u <- (n - 2) *       mean.beta
  v <- (n - 2) * (1 - mean.beta)
  beta.p <- noncentral.correlation.beta.test(rho, u, v)
  f <- dbeta((1+rho)/2,u,v)
  plot(rho, f, type='l', lwd=3, xlab="Correlation (R)", ylab="Density", c
       main=c(paste("R ~ Beta(",round(u,2),",",round(v,2),")",sep=''),
              paste("n =",n,"    E[R] =",round(rho0,3))))
  abline(h=c(0,1), lty=3, col='grey')
  abline(v=c(-2:2)/2, lty=3, col='grey')
  abline(v=rho0, lty=2, lwd=3, col='grey60')

  suppressWarnings(plot(rho, log10(beta.p), type='l', lwd=3, xlab="Correl
                        ylim=c(min.logp,0), col='dodgerblue', main="Pr{de
  abline(h=c(min.logp:0), lty=3, col='grey')
  abline(v=c(-2:2)/2, lty=3, col='grey')
  abline(v=rho0, lty=2, lwd=3, col='grey60')
  suppressWarnings(lines(rho,log10(correlation.Fisher.test(rho,rho0,n)),l
}
legend(-1,0,legend=c("Fisher","Beta"),lwd=c(1,3),col=c('black','dodgerblu
```

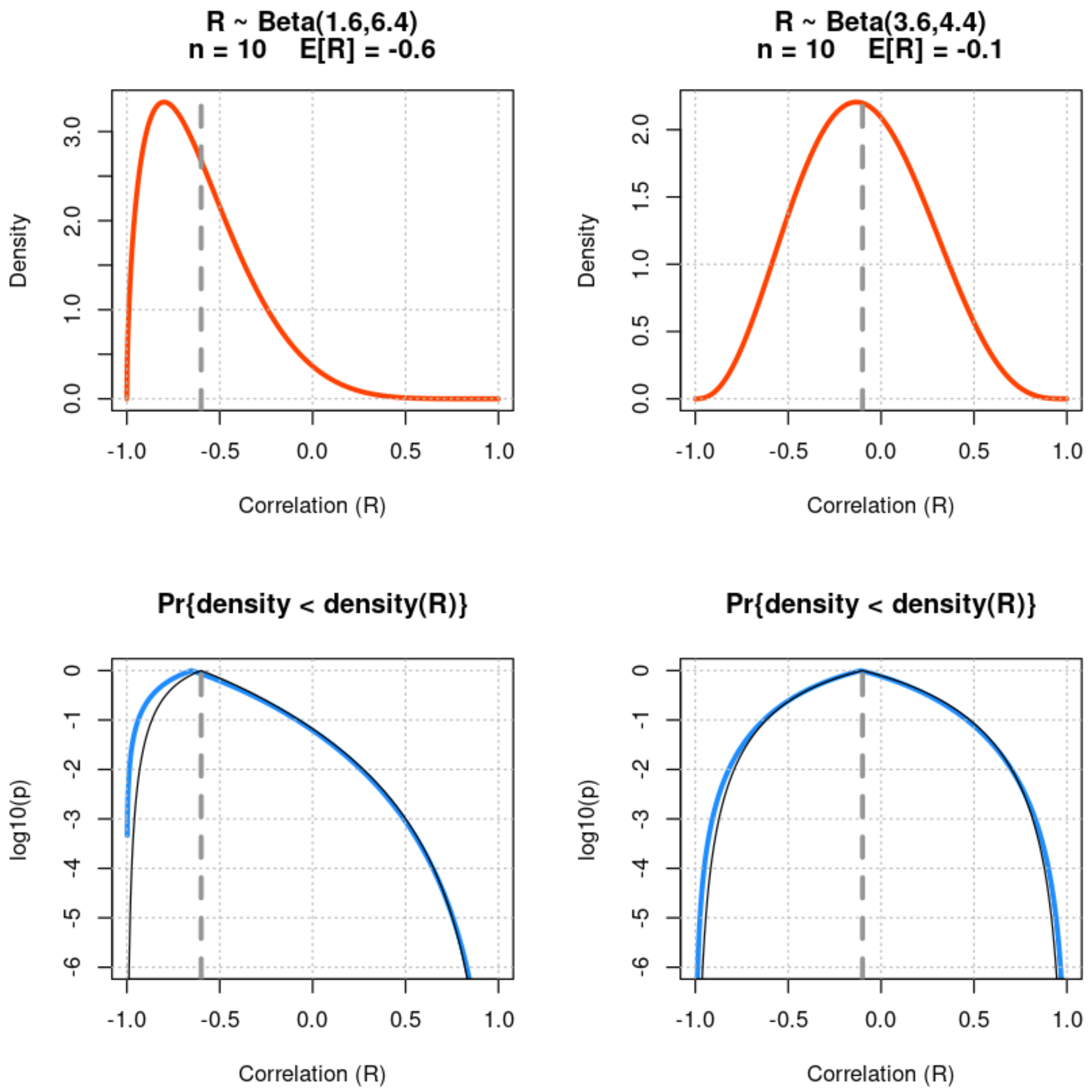
R ~ Beta(1.6,6.4)
n = 10 E[R] = -0.6
Density
Correlation (R)
R ~ Beta(3.6,4.4)
n = 10 E[R] = -0.1
Density
Correlation (R)
Pr{density < density(R)}
log10(p)
Correlation (R)
Pr{density < density(R)}
log10(p)
Correlation (R)

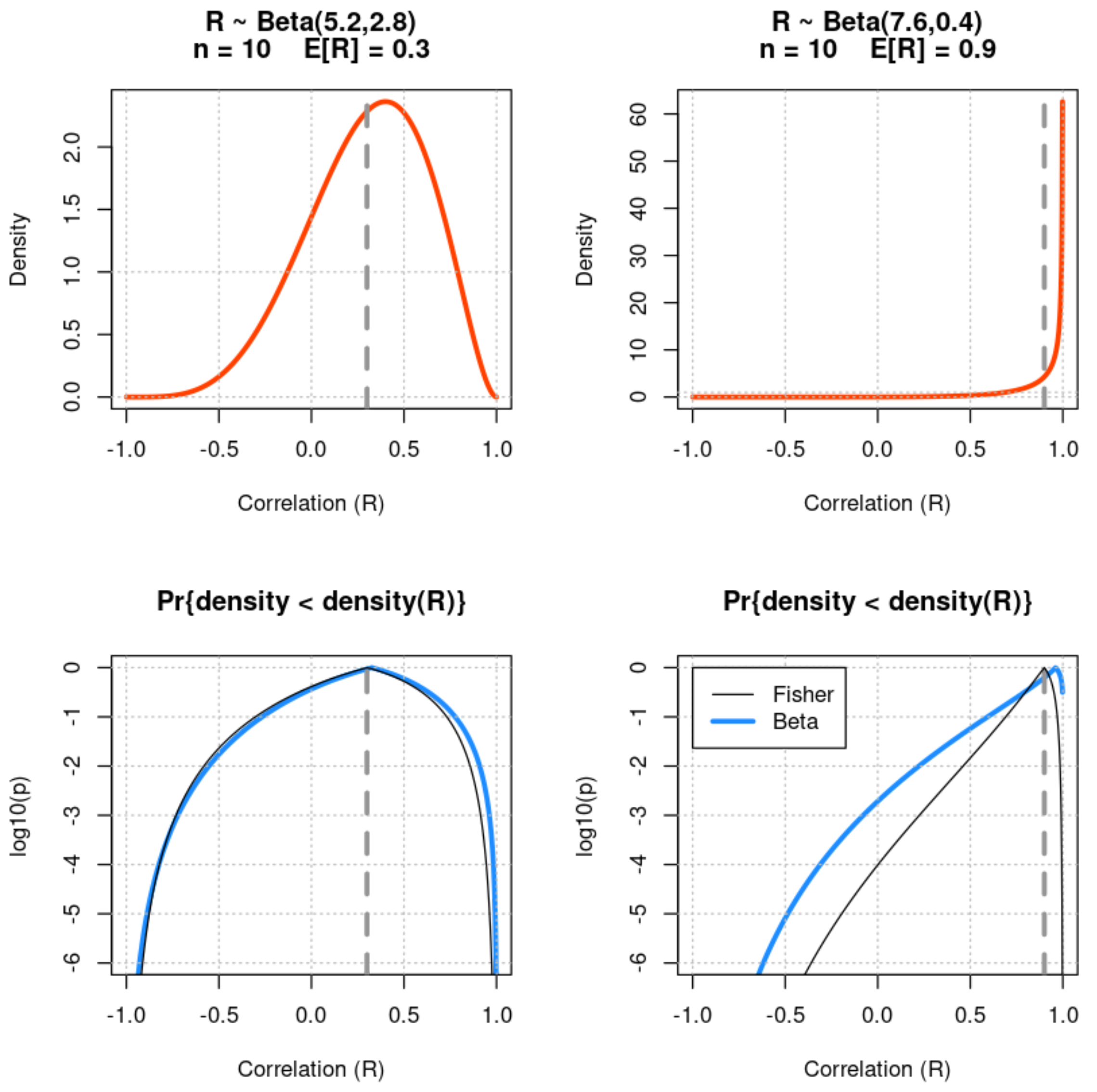


In [506…

```
par(mfcol=c(2,2))
n <- 7
min.logp <- -6
for (rho0 in c(-0.6,-0.1,0.3,0.9)) {
  mean.beta <- (1 + rho0)/2
  u <- (n - 2) *      mean.beta
  v <- (n - 2) * (1 - mean.beta)
  beta.p <- noncentral.correlation.beta.test(rho, u, v)
  f <- dbeta((1+rho)/2,u,v)
  plot(rho, f, type='l', lwd=3, xlab="Correlation (R)", ylab="Density", c
       main=c(paste("R ~ Beta(",round(u,2),",",round(v,2),")",sep=''),
              paste("n =",n,"   E[R] =",round(rho0,3))))
  abline(h=c(0,1), lty=3, col='grey')
  abline(v=c(-2:2)/2, lty=3, col='grey')
  abline(v=rho0, lty=2, lwd=3, col='grey60')

  suppressWarnings(plot(rho, log10(beta.p), type='l', lwd=3, xlab="Correl
                        ylim=c(min.logp,0), col='dodgerblue', main="Pr{de
  abline(h=c(min.logp:0), lty=3, col='grey')
  abline(v=c(-2:2)/2, lty=3, col='grey')
  abline(v=rho0, lty=2, lwd=3, col='grey60')
  suppressWarnings(lines(rho,log10(correlation.Fisher.test(rho,rho0,n)),l
}
legend(-1,0,legend=c("Fisher","Beta"),lwd=c(1,3),col=c('black','dodgerblu
```

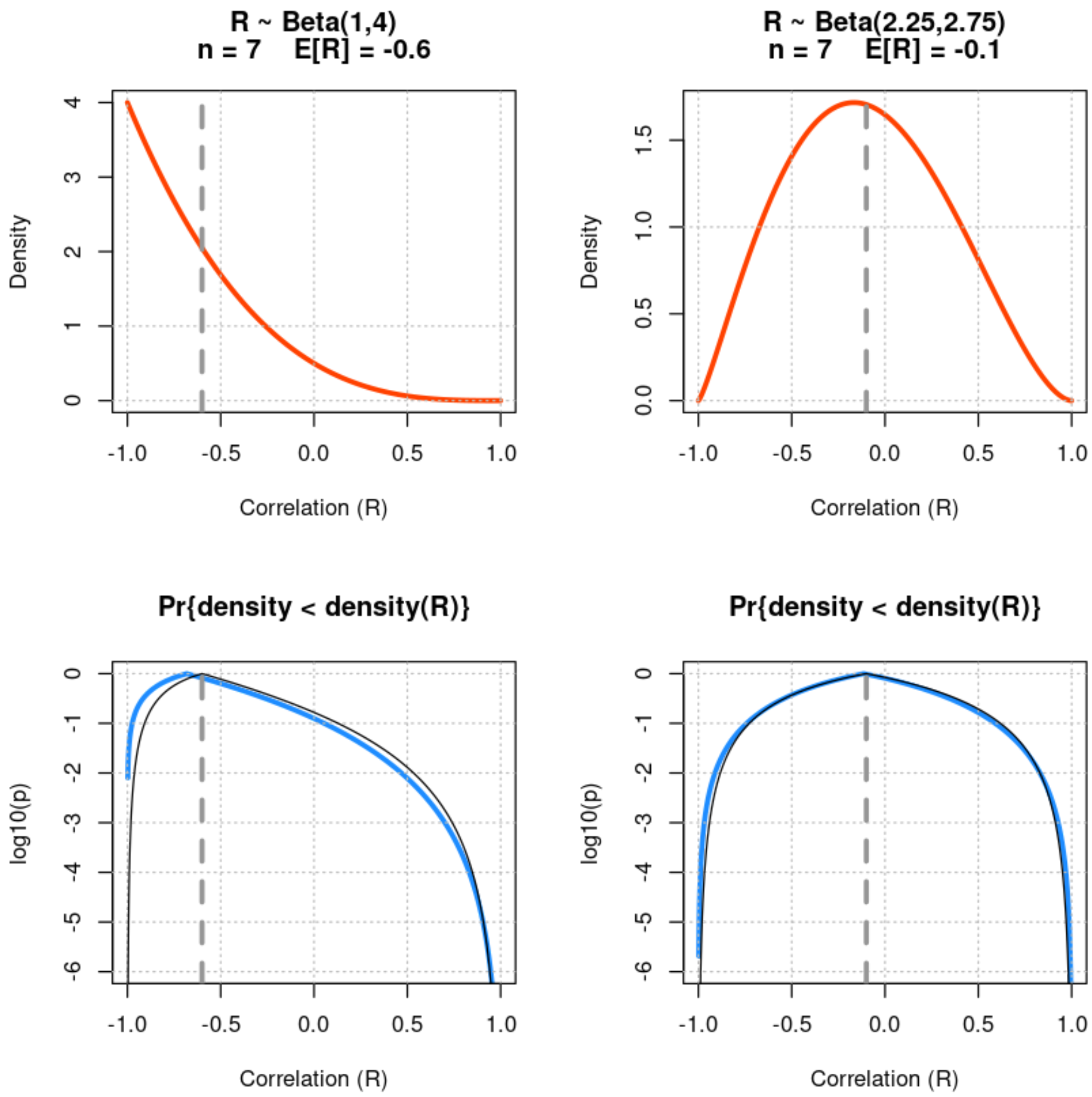
R ~ Beta(1,4)
n = 7 E[R] = -0.6
R ~ Beta(2.25,2.75)
n = 7 E[R] = -0.1
Density
Correlation (R)
Pr{density < density(R)}
log10(p)
-1.0
-0.5
0.0
0.5
1.0

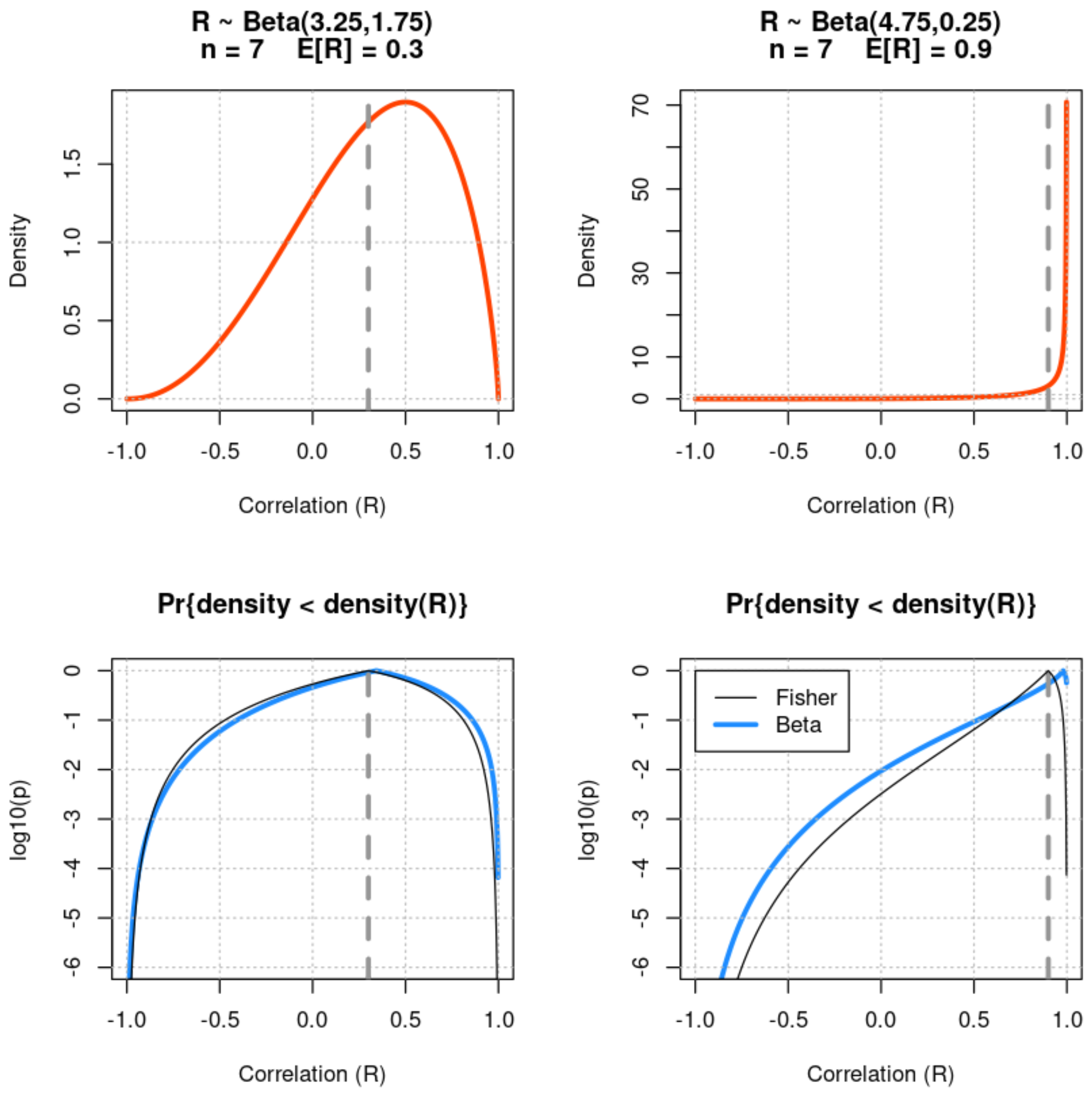


In [507…

```
par(mfcol=c(2,2))
n <- 6
min.logp <- -6
for (rho0 in c(-0.6,-0.1,0.3,0.9)) {
  mean.beta <- (1 + rho0)/2
  u <- (n - 2) *       mean.beta
  v <- (n - 2) * (1 - mean.beta)
  beta.p <- noncentral.correlation.beta.test(rho, u, v)
  f <- dbeta((1+rho)/2,u,v)
  plot(rho, f, type='l', lwd=3, xlab="Correlation (R)", ylab="Density", c
       main=c(paste("R ~ Beta(",round(u,2),",",round(v,2),")",sep=''),
              paste("n =",n,"    E[R] =",round(rho0,3))))
  abline(h=c(0,1), lty=3, col='grey')
  abline(v=c(-2:2)/2, lty=3, col='grey')
  abline(v=rho0, lty=2, lwd=3, col='grey60')

  suppressWarnings(plot(rho, log10(beta.p), type='l', lwd=3, xlab="Correl
                        ylim=c(min.logp,0), col='dodgerblue', main="Pr{de
  abline(h=c(min.logp:0), lty=3, col='grey')
  abline(v=c(-2:2)/2, lty=3, col='grey')
  abline(v=rho0, lty=2, lwd=3, col='grey60')
  suppressWarnings(lines(rho,log10(correlation.Fisher.test(rho,rho0,n)),l
}
legend(-1,0,legend=c("Fisher","Beta"),lwd=c(1,3),col=c('black','dodgerblu
```

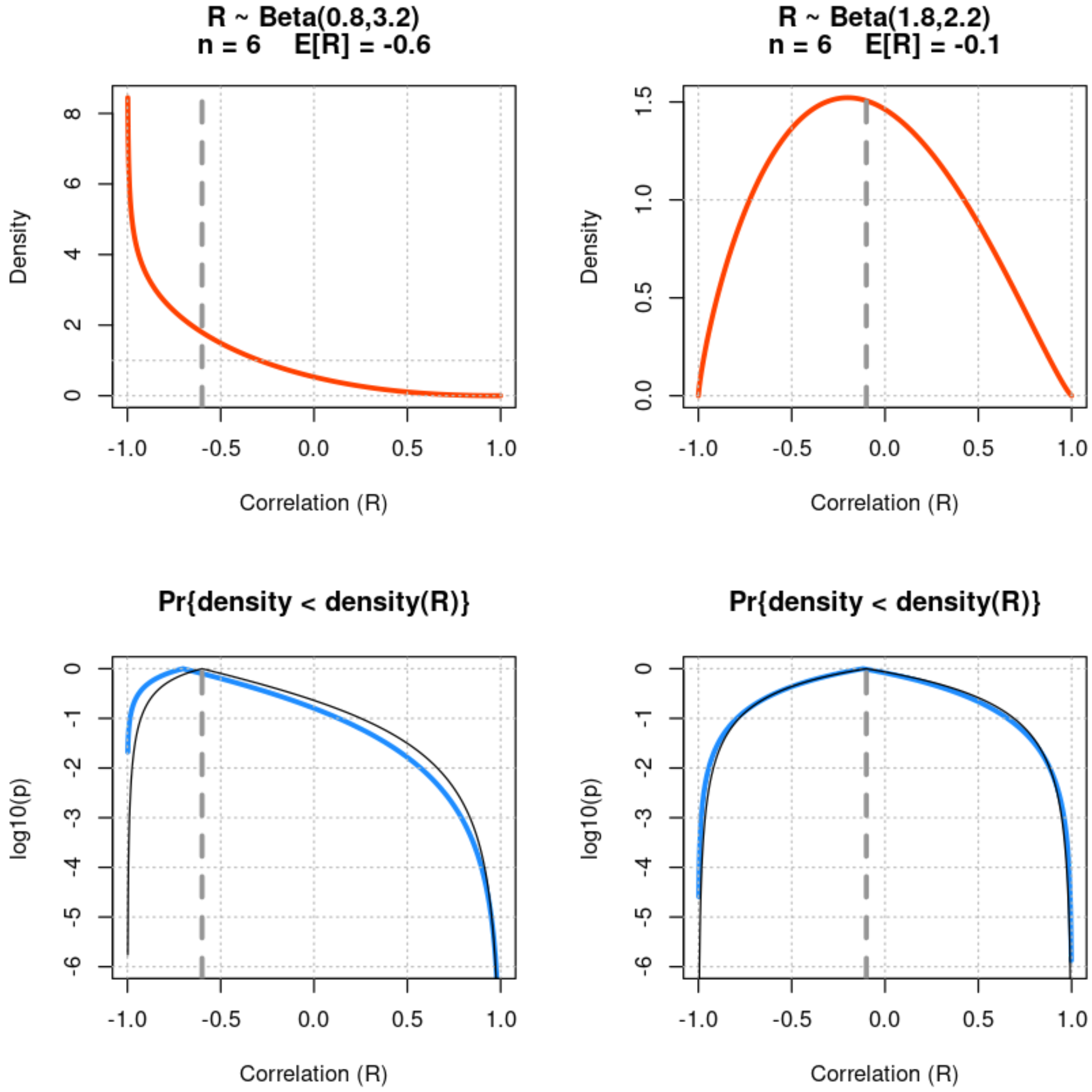
R ~ Beta(0.8,3.2)
n = 6    E[R] = -0.6
R ~ Beta(1.8,2.2)
n = 6    E[R] = -0.1
Density
Correlation (R)
Pr{density < density(R)}
log10(p)

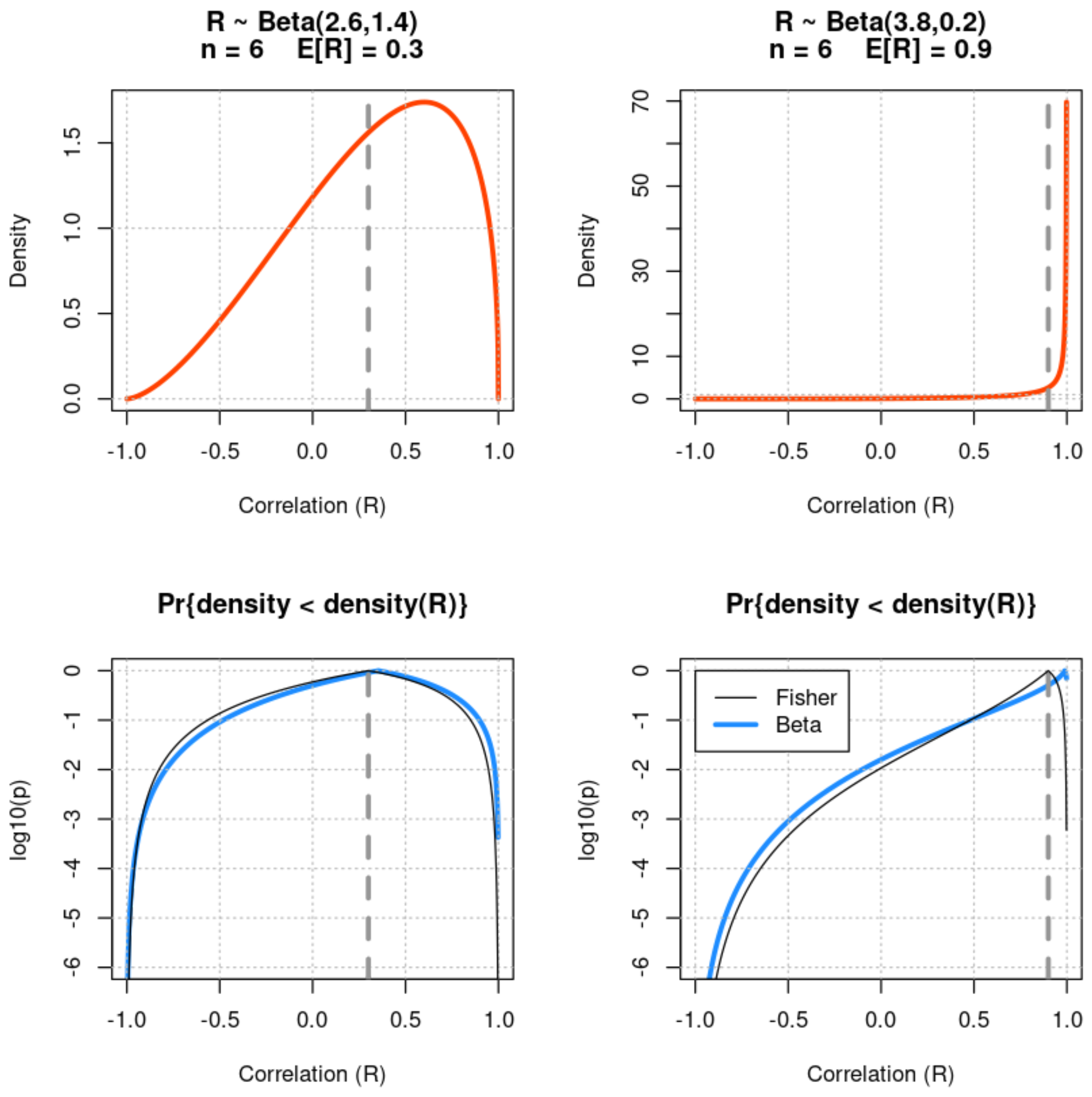


In [508…

```
par(mfcol=c(2,2))
n <- 5
min.logp <- -6
for (rho0 in c(-0.6,-0.1,0.3,0.9)) {
  mean.beta <- (1 + rho0)/2
  u <- (n - 2) *      mean.beta
  v <- (n - 2) * (1 - mean.beta)
  beta.p <- noncentral.correlation.beta.test(rho, u, v)
  f <- dbeta((1+rho)/2,u,v)
  plot(rho, f, type='l', lwd=3, xlab="Correlation (R)", ylab="Density", c
       main=c(paste("R ~ Beta(",round(u,2),",",round(v,2),")",sep=''),
              paste("n =",n,"    E[R] =",round(rho0,3))))
  abline(h=c(0,1), lty=3, col='grey')
  abline(v=c(-2:2)/2, lty=3, col='grey')
  abline(v=rho0, lty=2, lwd=3, col='grey60')

  suppressWarnings(plot(rho, log10(beta.p), type='l', lwd=3, xlab="Correl
                        ylim=c(min.logp,0), col='dodgerblue', main="Pr{de
  abline(h=c(min.logp:0), lty=3, col='grey')
  abline(v=c(-2:2)/2, lty=3, col='grey')
  abline(v=rho0, lty=2, lwd=3, col='grey60')
  suppressWarnings(lines(rho,log10(correlation.Fisher.test(rho,rho0,n)),l
}
legend(-1,0,legend=c("Fisher","Beta"),lwd=c(1,3),col=c('black','dodgerblu
```

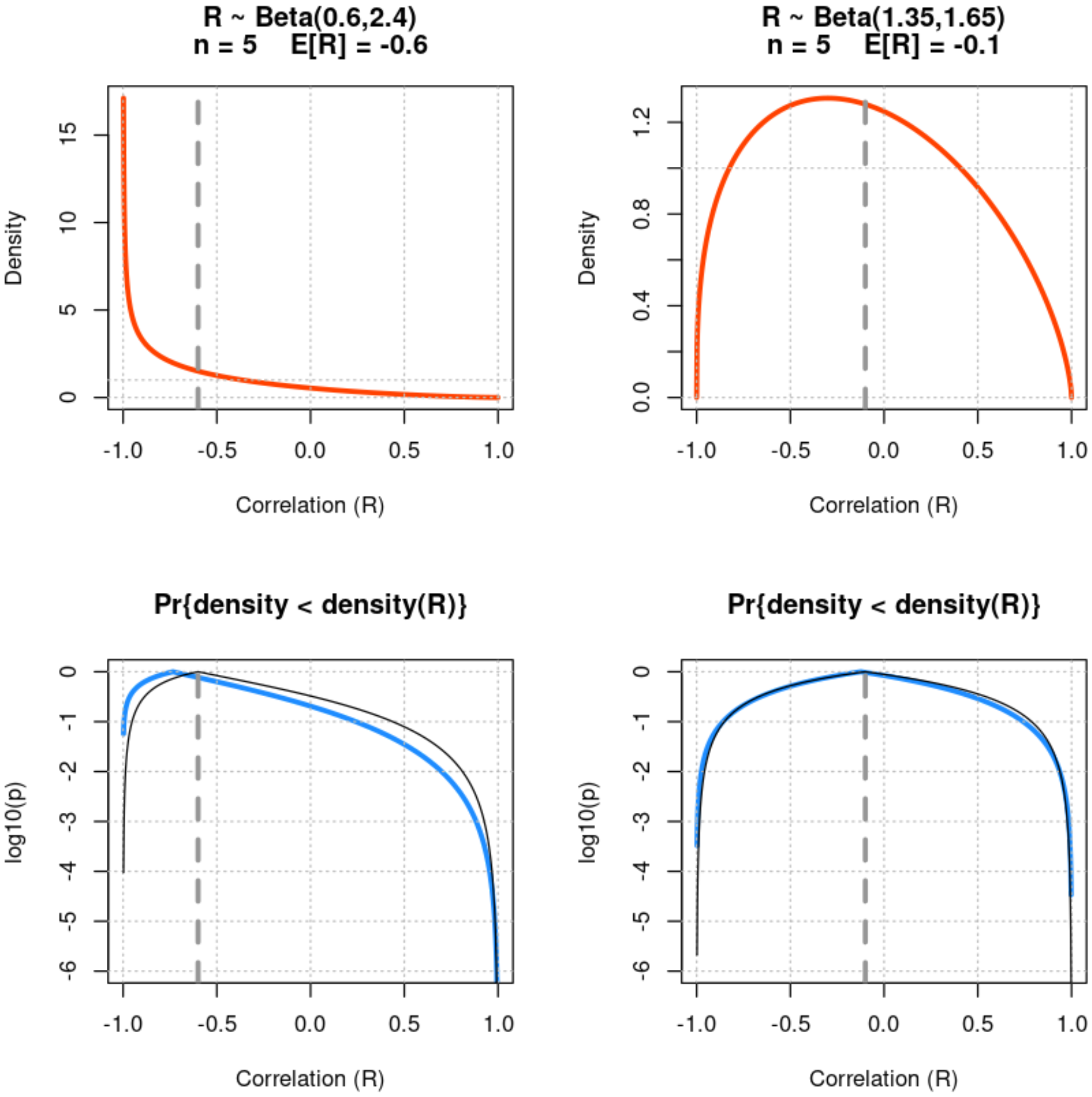
R ~ Beta(0.6,2.4)
n = 5 E[R] = -0.6
R ~ Beta(1.35,1.65)
n = 5 E[R] = -0.1
Density
Correlation (R)
Pr{density < density(R)}
Pr{density < density(R)}
log10(p)
Correlation (R)

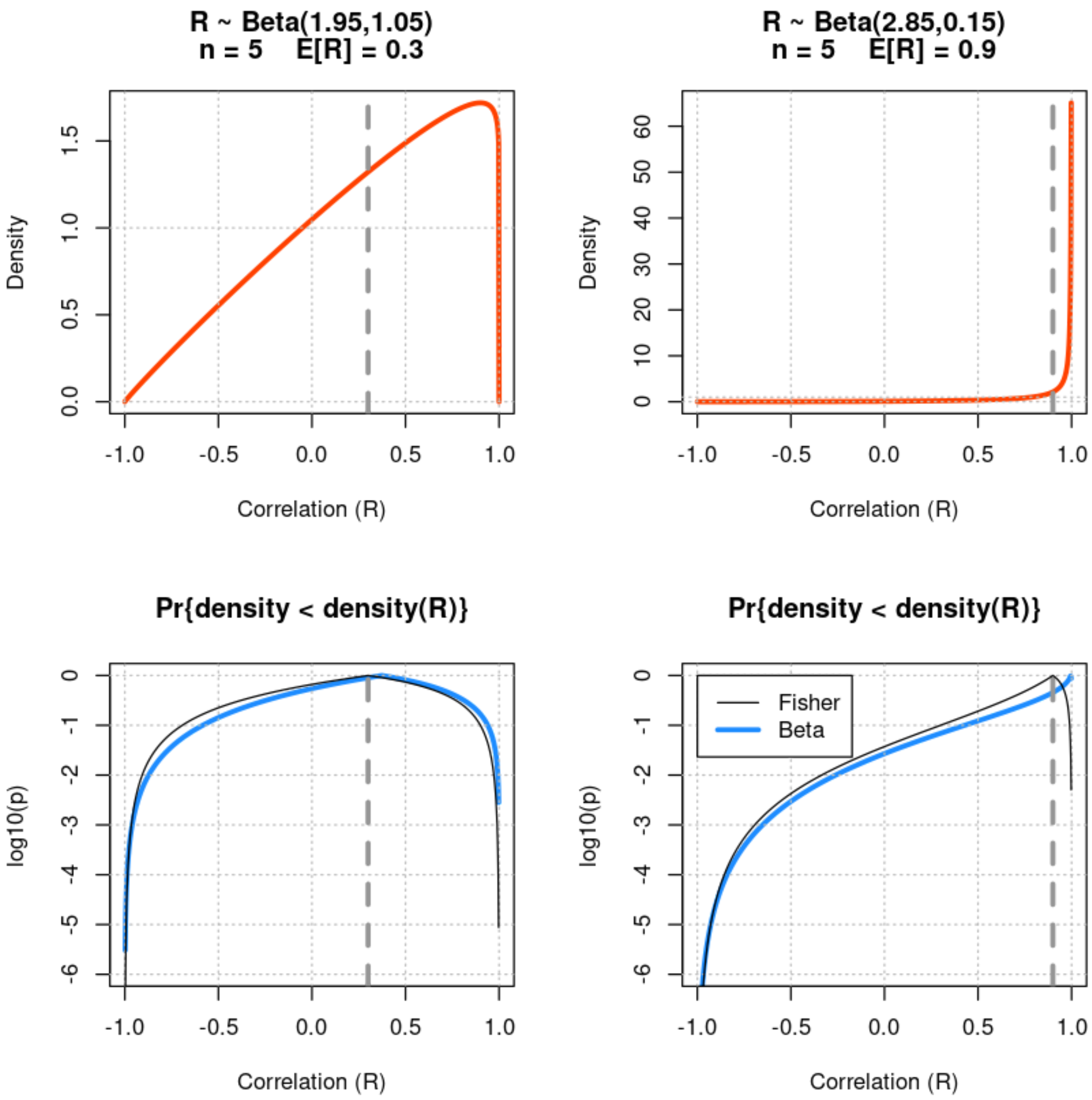


In [509…

```
par(mfcol=c(2,2))
n <- 4
min.logp <- -6
for (rho0 in c(-0.6,-0.1,0.3,0.9)) {
  mean.beta <- (1 + rho0)/2
  u <- (n - 2) *       mean.beta
  v <- (n - 2) * (1 - mean.beta)
  beta.p <- noncentral.correlation.beta.test(rho, u, v)
  f <- dbeta((1+rho)/2,u,v)
  plot(rho, f, type='l', lwd=3, xlab="Correlation (R)", ylab="Density", c
       main=c(paste("R ~ Beta(",round(u,2),",",round(v,2),")",sep=''),
              paste("n =",n,"    E[R] =",round(rho0,3))))
  abline(h=c(0,1), lty=3, col='grey')
  abline(v=c(-2:2)/2, lty=3, col='grey')
  abline(v=rho0, lty=2, lwd=3, col='grey60')

  suppressWarnings(plot(rho, log10(beta.p), type='l', lwd=3, xlab="Correl
                        ylim=c(min.logp,0), col='dodgerblue', main="Pr{de
  abline(h=c(min.logp:0), lty=3, col='grey')
  abline(v=c(-2:2)/2, lty=3, col='grey')
  abline(v=rho0, lty=2, lwd=3, col='grey60')
  suppressWarnings(lines(rho,log10(correlation.Fisher.test(rho,rho0,n)),l
}
legend(-1,0,legend=c("Fisher","Beta"),lwd=c(1,3),col=c('black','dodgerblu
```

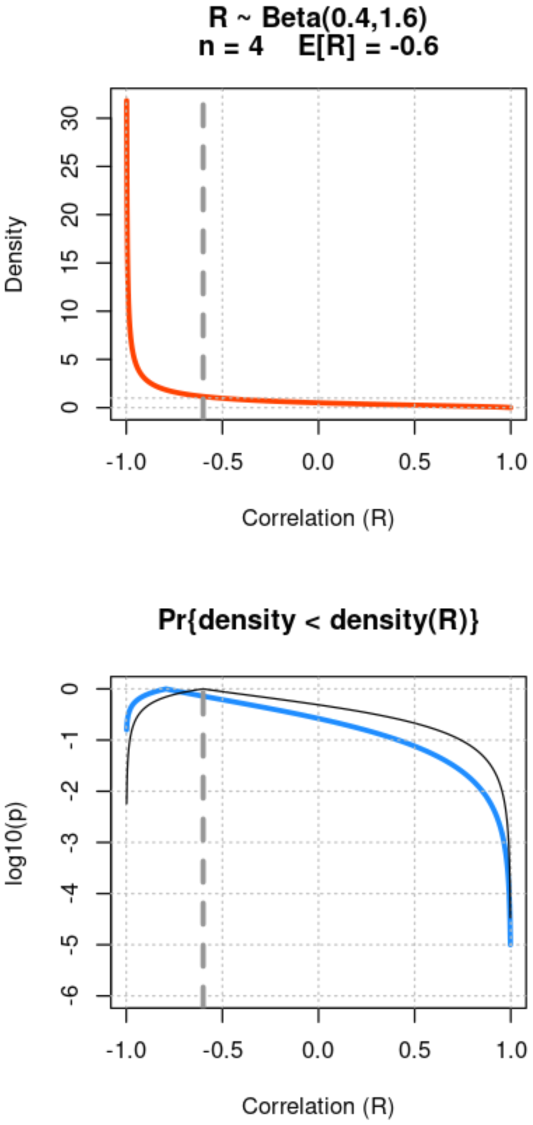
R ~ Beta(0.4,1.6)
n = 4 E[R] = -0.6
R ~ Beta(0.9,1.1)
n = 4 E[R] = -0.1
Density
Correlation (R)
Pr{density < density(R)}
Pr{density < density(R)}
log10(p)
Correlation (R)

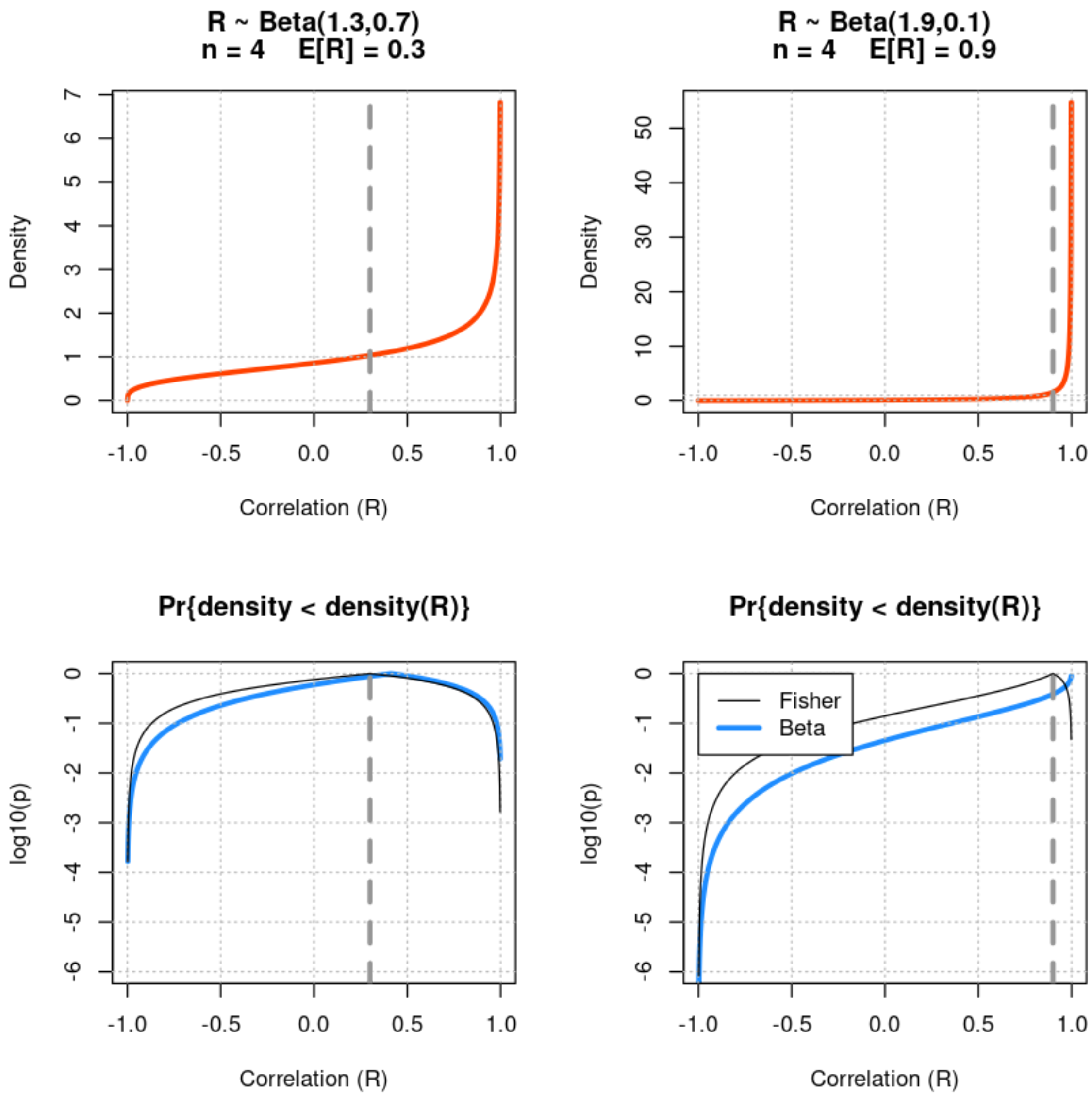


#### Conclusions

Student's *t*-test when the data is expected to be uncorrelated and the corresponding Beta null model test are identical, which provides strong confirmation that the Beta null model's dependence on the dimensionality of the data is correct. When nonzero correlation is expected, however, Fisher's transformation provides a `z` statistic that is known to be approximately Normally distributed, and whose distribution is known to converge to a Normal distribution as the number of dimensions increases without bound. When the sample size is small (`n < 30`) and the expected association between the points is moderate (`|E[r]| << 0.9`), when the Fisher transformation is expected to be reasonably accurate, the simulations above show that the test based on a Beta null model is approximated by the test based on the Fisher transformation. The Beta distribution is also known to converge to a Normal distribution as the number of dimensions (and therefore its shape parameters) increases without bound, but its mean remains constant. All of these results appear to be consistent with the Beta null model being an appropriate candidate for the exact distribution of spurious correlations, even when `E[r] != 0`, and even potentially for `|E[r]| >= 0.9`. The Beta null model presented here therefore represents a single approach to significance testing for correlation coefficients that as general, and potentially more accurate than the Fisher transformation approach, that is equivalent to the exact *t*-test approach when it applies. Whether or not the Beta null model is exact regardless of the expected correlation remains to be formally proven, however none of our results, or the results of our other simulations (see Supplement S2), are inconsistent with that hypothesis.
