## Supplement S3 for "Model-based Standardization of Correlation Coefficients Improves Multi-Omic Clustering and Biological Signal Discovery"

### Preliminaries

In [1]:
```
options(jupyter.plot_scale=1,
        width=200,
        repr.matrix.max.cols=200,
        repr.matrix.max.rows=Inf)
```

### Simulations of spurious correlations: dimension and marginal distribution

We have simulated vectors from several iid marginal distributions, all with zero mean, to identify potential requirements to produce a $\rho \sim \mathrm{Beta}(n/2-1, n/2-1)$ null distribution for correlations among $n$-dimensional points (vectors). Briefly we find:

1. The marginal distributions we simulated which were symmetric around $0$ with finite variance appear to produce the expected null model.
2. An asymmetric distribution with finite variance appears to approach the expected null model asymptotically in $n$.
3. Symmetric distributions with nonfinite variance did not appear to produce the expected null model.

We conclude that while some marginal distributions do not produce the proposed null model, some "well-behaved" distributions do.

In [2]:
```
###
# Demonstration that n-D correlations are distributed as
#      r ~ 2 Beta(n/2-1, n/2-1) - 1.
# We therefore can use the null model
#     density(r | n dimensions) = dbeta((1+r)/2, n/2-1, n/2-1)
###
# Estimating the effective dimensionality of the data
###
# Since there is a straightforward relationship between
# the expected distribution and the data's dimensionality,
# we explore how to estimate the effective dimensionality
# of a dataset from the observed distribution of
# correlations among the observations.
###
# The variance of Beta(v,w) is
#     var(B) = vw / ((v+w)^2 (v+w+1))
# Substituting w = v,
#     var(B) = v^2 / (4v^2 (2v+1))
#     var(B) = 1 / 4 (2v+1)
# Solving for v,
#     4 (2v + 1) = 1/var(B)
#        2v      = 1/(4var(B)) - 1
#         v      = (1/(4 var(B)) - 1)/2
# Substituting v = n/2 - 1,
#     n/2 - 1    = (1/(4 var(B)) - 1)/2
#     n - 2      =  1/(4 var(B)) - 1
```

```
#     n          =  1/(4 var(B)) + 1
# Thus we can estimate the dimensionality of (mostly random)
# correlation data as
#     d = 1 + 1 /(4 var(B))
# However,
#     var(r) = var(2 B - 1) = var(2 B) = 4 var(B),
# so
#     d = 1 + 1 / var(r)
#
# This estimate is NOT expected to be robust to outliers.
# Instead, we suggest a robust estimate of the dimensionality
# by fitting a beta distribution B* to the center of the observed
# distribution of (1+r)/2 (by "center" we mean the region with
# density at least half the density at the mode), and using
# the estimate
#     d = 1 + 1 / (4 var(B*)).
###
```

In [3]:

```
# M random vectors generate C = M(M-1) / 2 non-self unique spurious corre
# When M = 10000, C = 49,995,000.

M <- 10000
Ns <- c(3,4,5,6,
        7,8,9,10,
        20,30,40,50,
        100,200,300,400)
```

In [4]:

```
marginal <- function(...) { rnorm(...) }

hist(marginal(100000),main="Gaussian marginal", xlab='z',
     col='gray', border='gray', prob=TRUE, xlim=c(-4,4), nclass=101)
xx <- c(-400:400)/100
abline(h=0); abline(v=c(-4:4),lty=3)
lines(xx,dnorm(xx),lwd=3,col='orangered')
legend(3,0.35,legend = c('sample','model'),col=c('gray','orangered'),bg='
box()
```

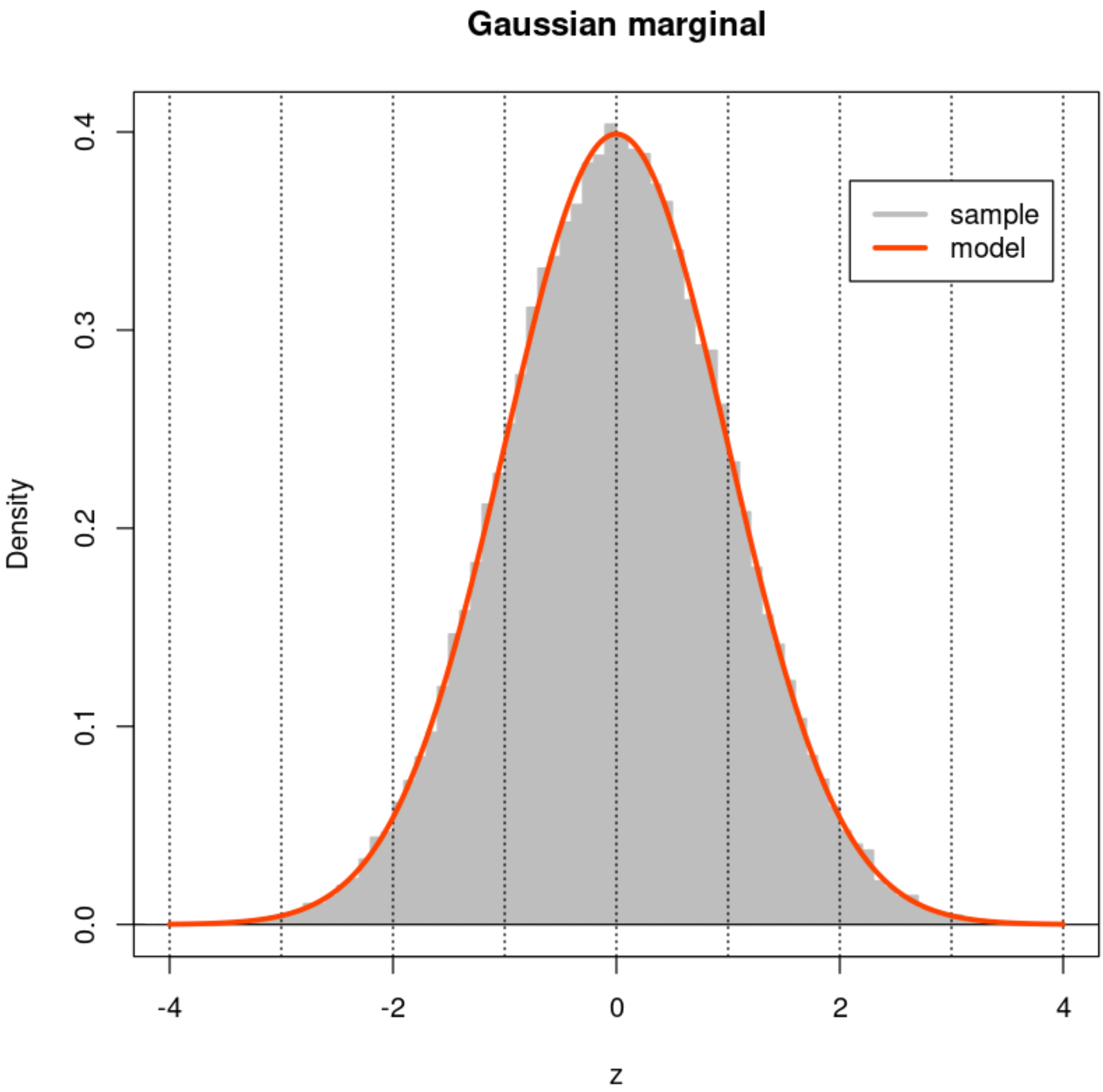


In [5]:
```
fine <- 101
par(mfrow=c(2,2))
xx <- (c(-50:49) + 0.05)/50
for (n in Ns) {
  X <- matrix(marginal(n*M),ncol=M,nrow=n)
  Z <- cor(X,method='p')
  hist(Z[row(Z) < col(Z)],nclass=fine,xlab="Correlation (r)",
       main=paste(n,"-vectors",sep=''),
       col='gray', border='gray', prob=TRUE, xlim=c(-1,1))
  lines(xx, dbeta((1+xx)/2,n/2-1,n/2-1)/2, col='orangered',lwd=3)
  abline(h=0) ; abline(v=c(-1:1),lty=3)
  box()
}
```

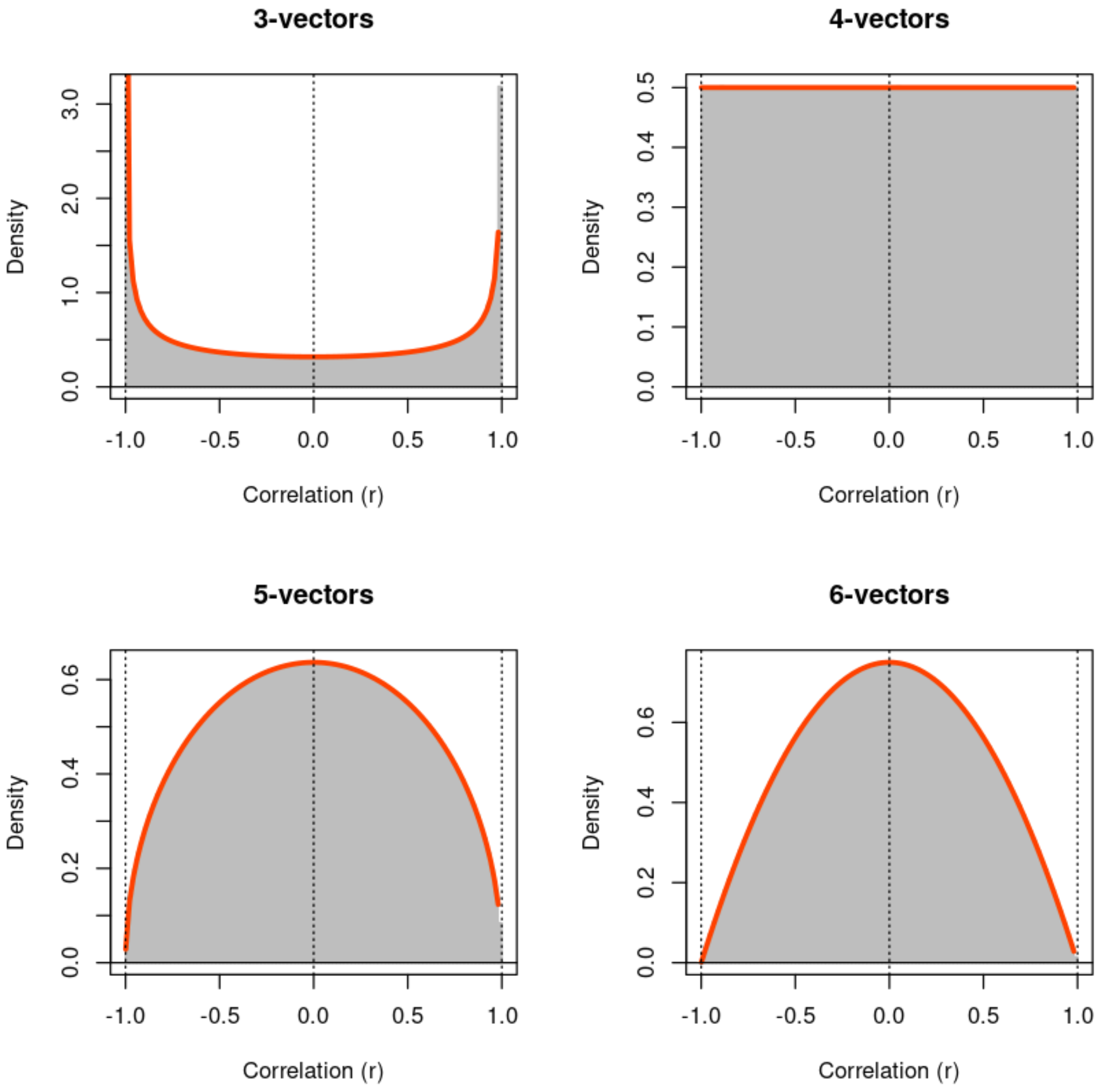
3-vectors
4-vectors
5-vectors
6-vectors
Density
Correlation (r)
-1.0
-0.5
0.0
0.5
1.0
3.0
2.0
1.0
0.5
0.4
0.3
0.2
0.1
0.6

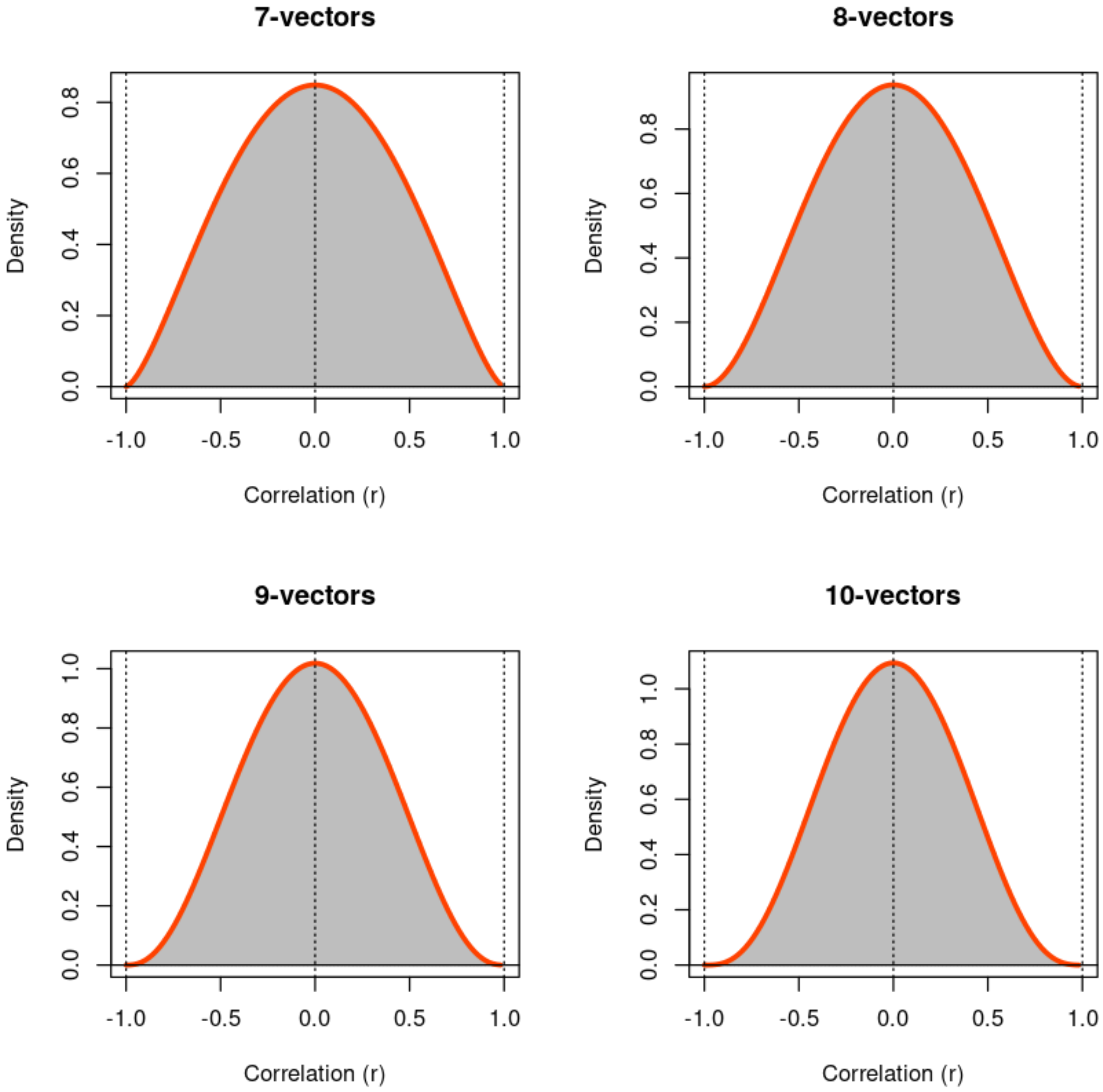
7-vectors
8-vectors
9-vectors
10-vectors
Density
Correlation (r)
-1.0
-0.5
0.0
0.5
1.0
0.2
0.4
0.6
0.8
1.0

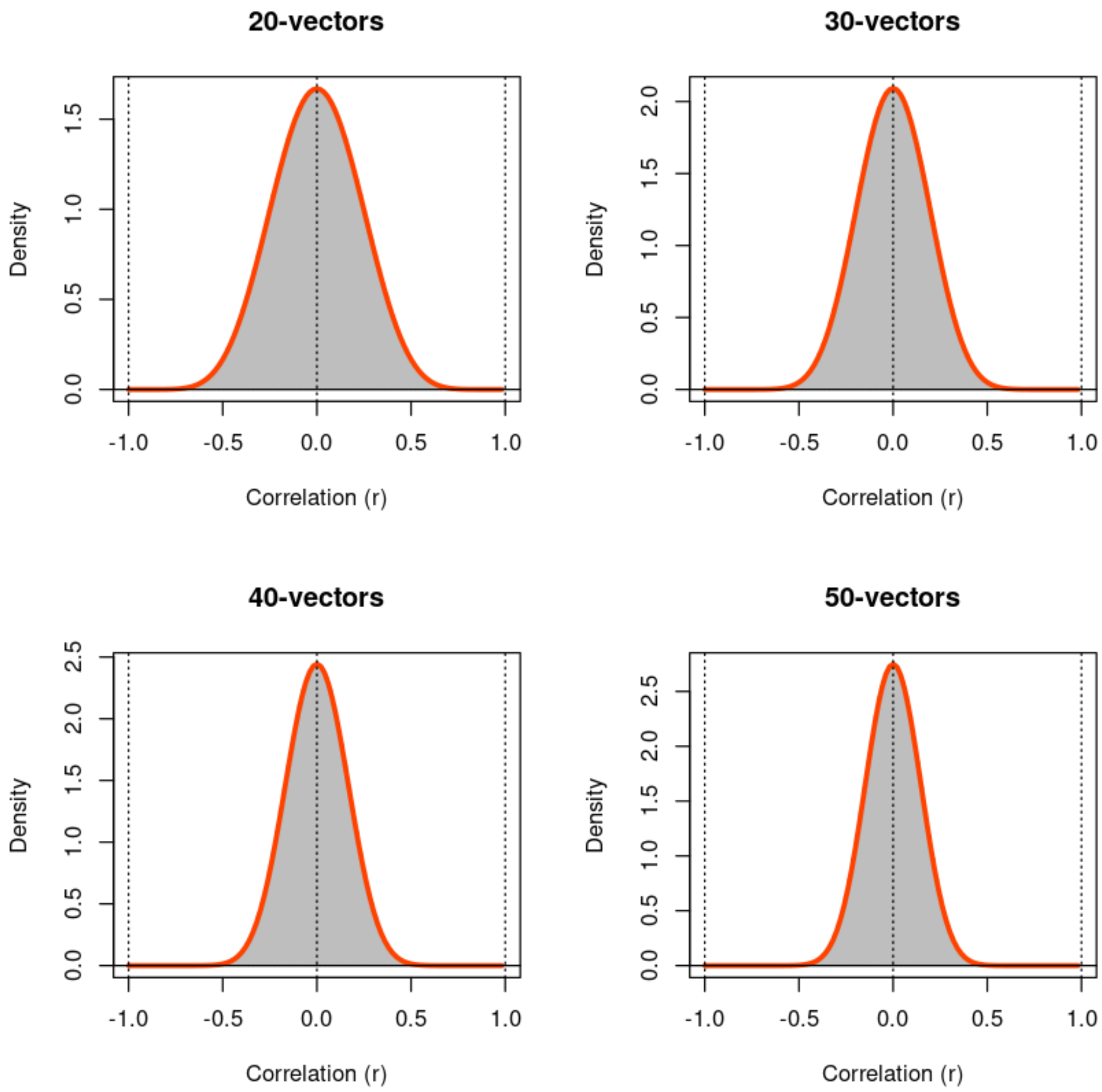
20-vectors
30-vectors
40-vectors
50-vectors
Density
Correlation (r)
-1.0
-0.5
0.0
0.5
1.0
1.5
2.0
2.5

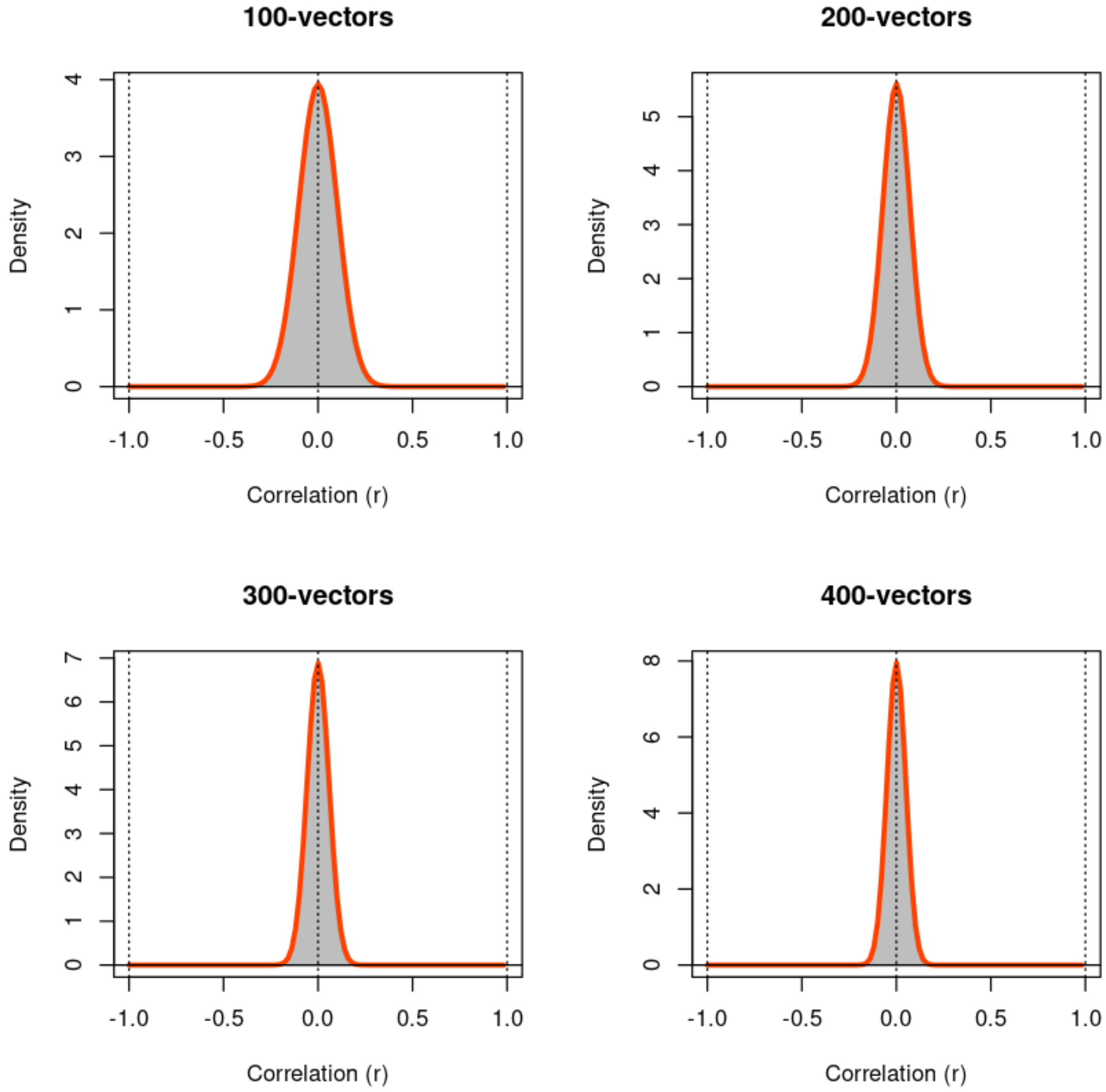


In [6]:

```
marginal <- function(...) { 2*runif(...)-1 }

hist(marginal(100000),main="Uniform marginal", xlab='U',
     col='gray', border='gray', prob=TRUE, xlim=c(-1,1), nclass=101)
xx <- c(-100:100)/100
abline(h=0); abline(v=c(-1:1),lty=3)
lines(xx,dunif((1+xx)/2)/2,lwd=3,col='orangered')
legend(0.75,0.35,legend = c('sample','model'),col=c('gray','orangered'),b
box()
```

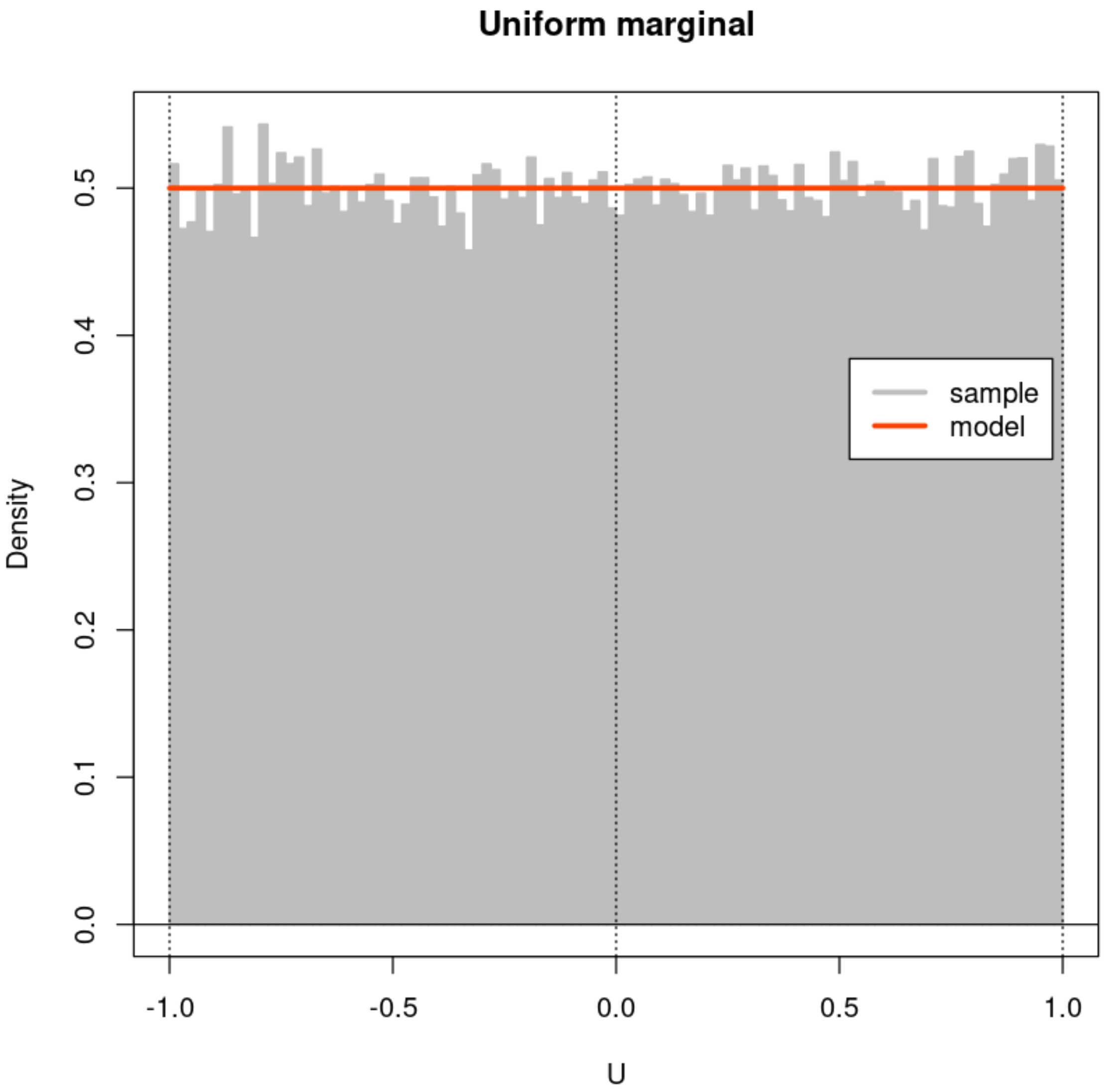


In [7]:

```
fine <- 101
par(mfrow=c(2,2))
xx <- (c(-50:49) + 0.05)/50
for (n in Ns) {
  X <- matrix(marginal(n*M),ncol=M,nrow=n)
  Z <- cor(X,method='p')
  hist(Z[row(Z) < col(Z)],nclass=fine,xlab="Correlation (r)",
       main=paste(n,"-vectors",sep=''),
       col='gray', border='gray', prob=TRUE, xlim=c(-1,1))
  lines(xx, dbeta((1+xx)/2,n/2-1,n/2-1)/2, col='orangered',lwd=3)
  abline(h=0) ; abline(v=c(-1:1),lty=3)
  box()
}
```

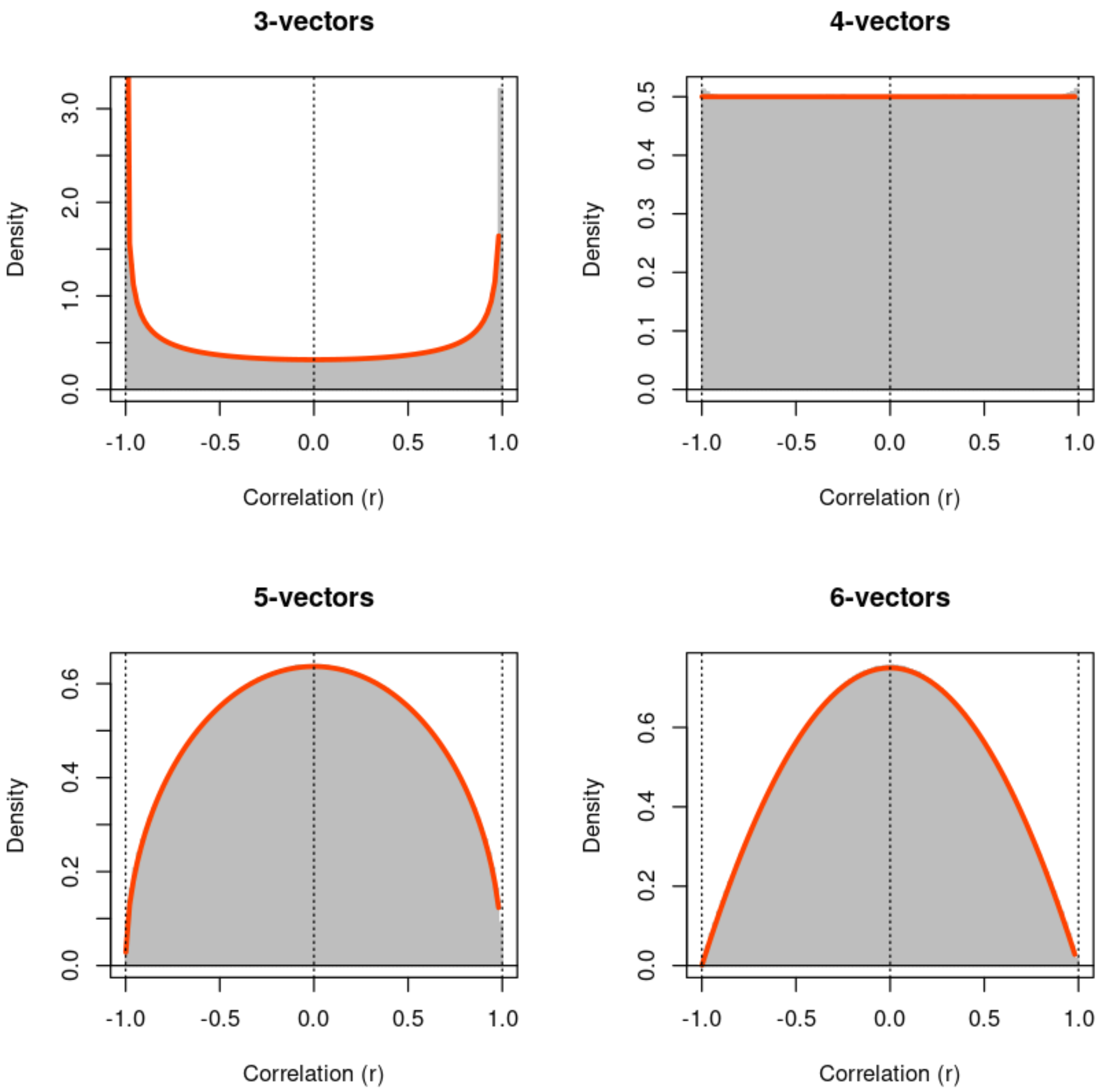

3-vectors
4-vectors
5-vectors
6-vectors
Density
Correlation (r)
-1.0
-0.5
0.0
0.5
1.0

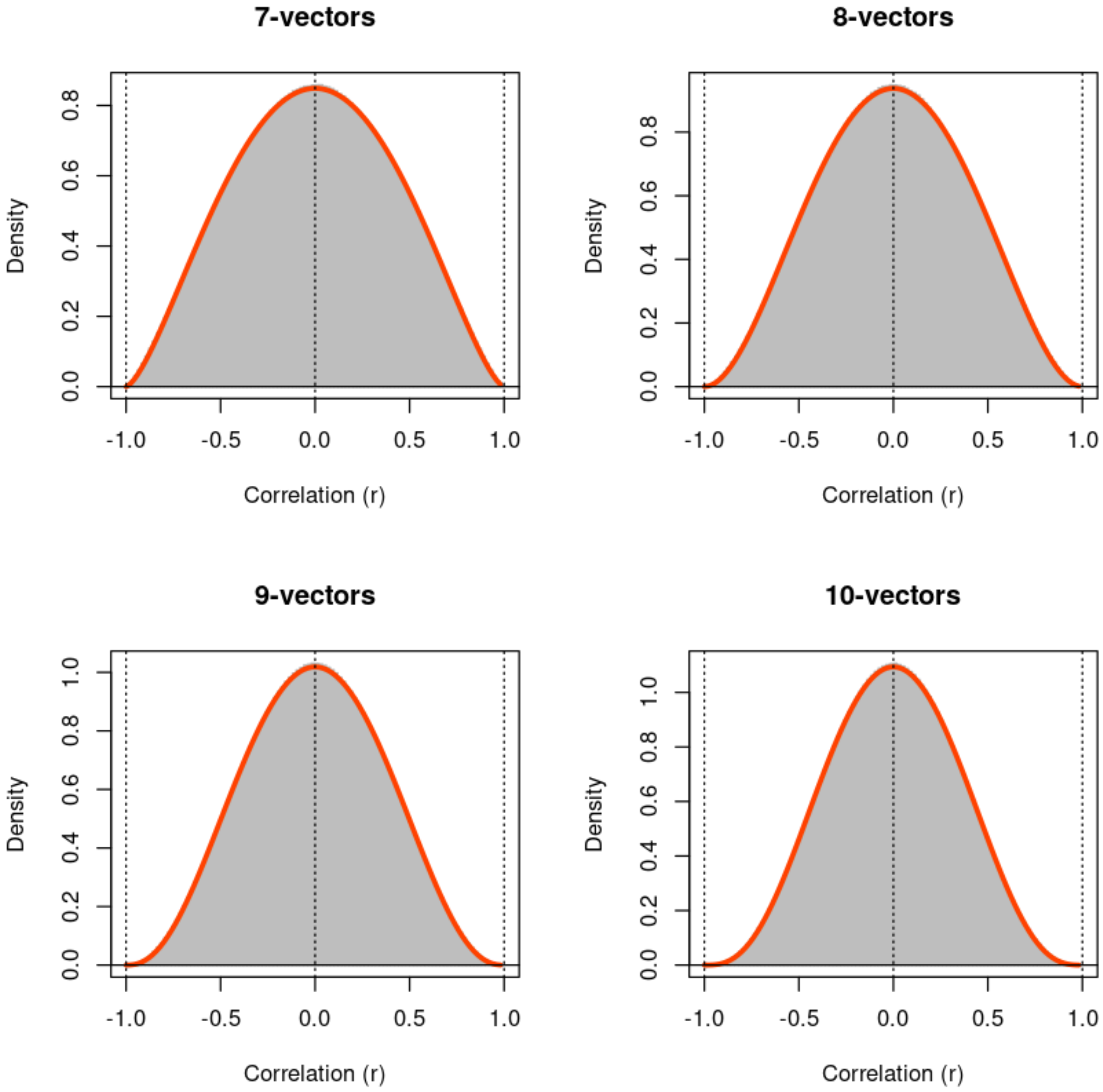
7-vectors
8-vectors
9-vectors
10-vectors
Density
Correlation (r)
-1.0
-0.5
0.0
0.5
1.0
0.2
0.4
0.6
0.8

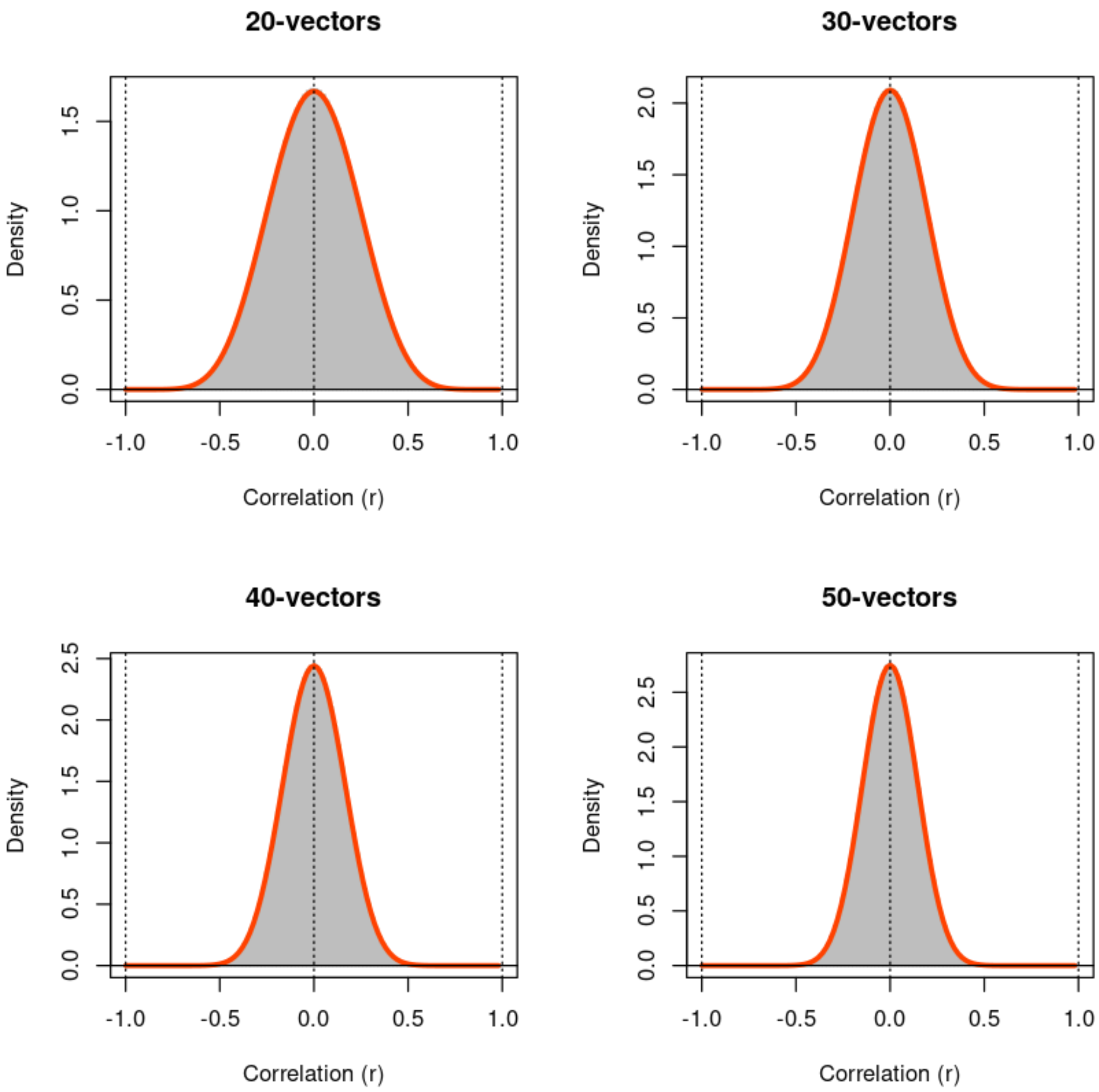
20-vectors
30-vectors
40-vectors
50-vectors
Density
Correlation (r)
-1.0
-0.5
0.0
0.5
1.0
1.5
2.0
2.5

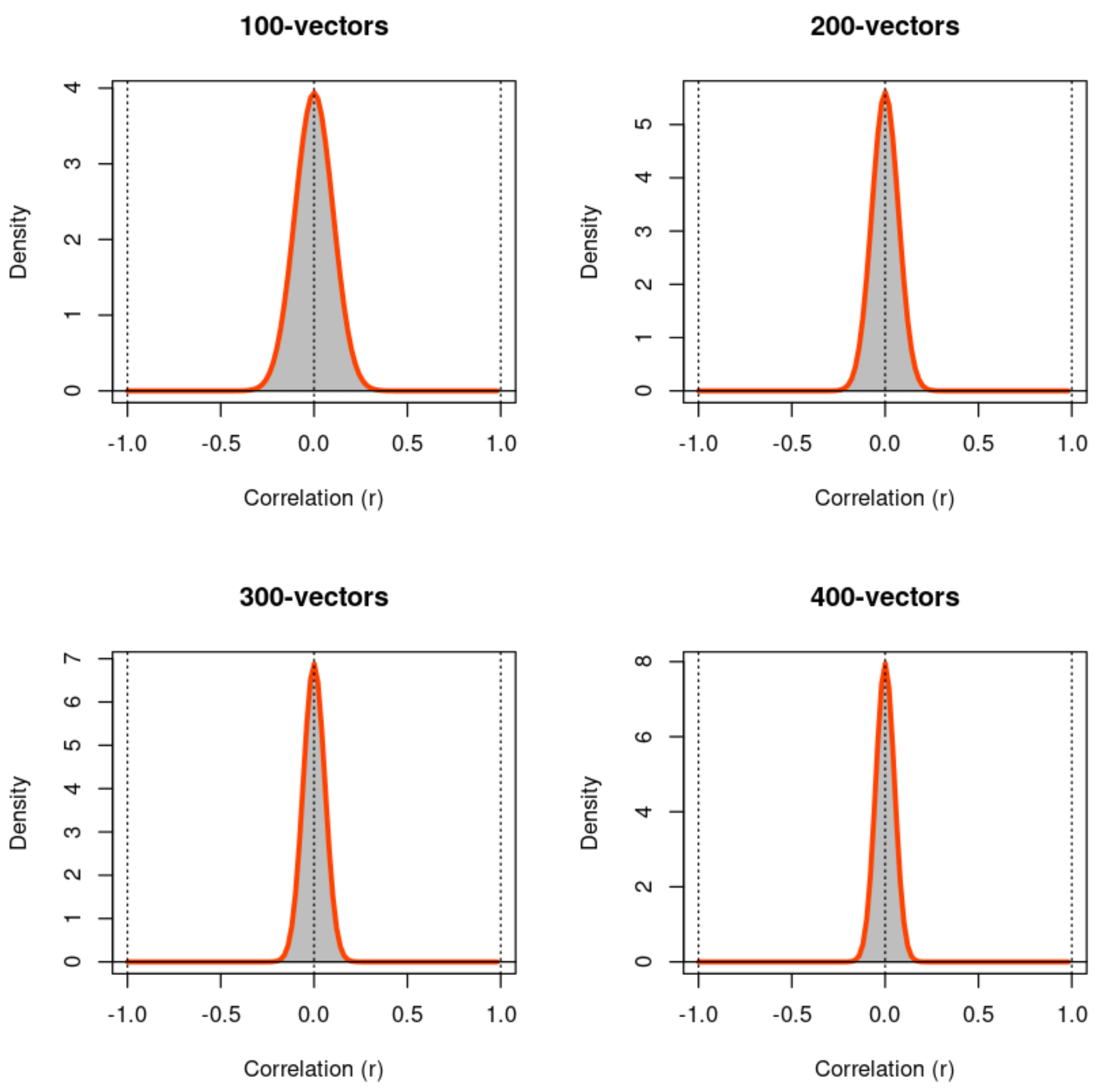


In [8]:

```
marginal <- function(...) { ifelse(runif(...) < 1/2, -1, 1) * rexp(...) }

hist(marginal(100000),main="Symmetrized exponential marginal", xlab='x',
     col='gray', border='gray', prob=TRUE, xlim=c(-6,6), nclass=201)
xx <- c(0:600)/100
abline(h=0); abline(v=c(-6:6),lty=3)
lines(xx,dexp(xx)/2,lwd=3,col='orangered')
lines(-xx,dexp(xx)/2,lwd=3,col='orangered')
legend(4,0.35,legend = c('sample','model'),col=c('gray','orangered'),bg='
box()
```

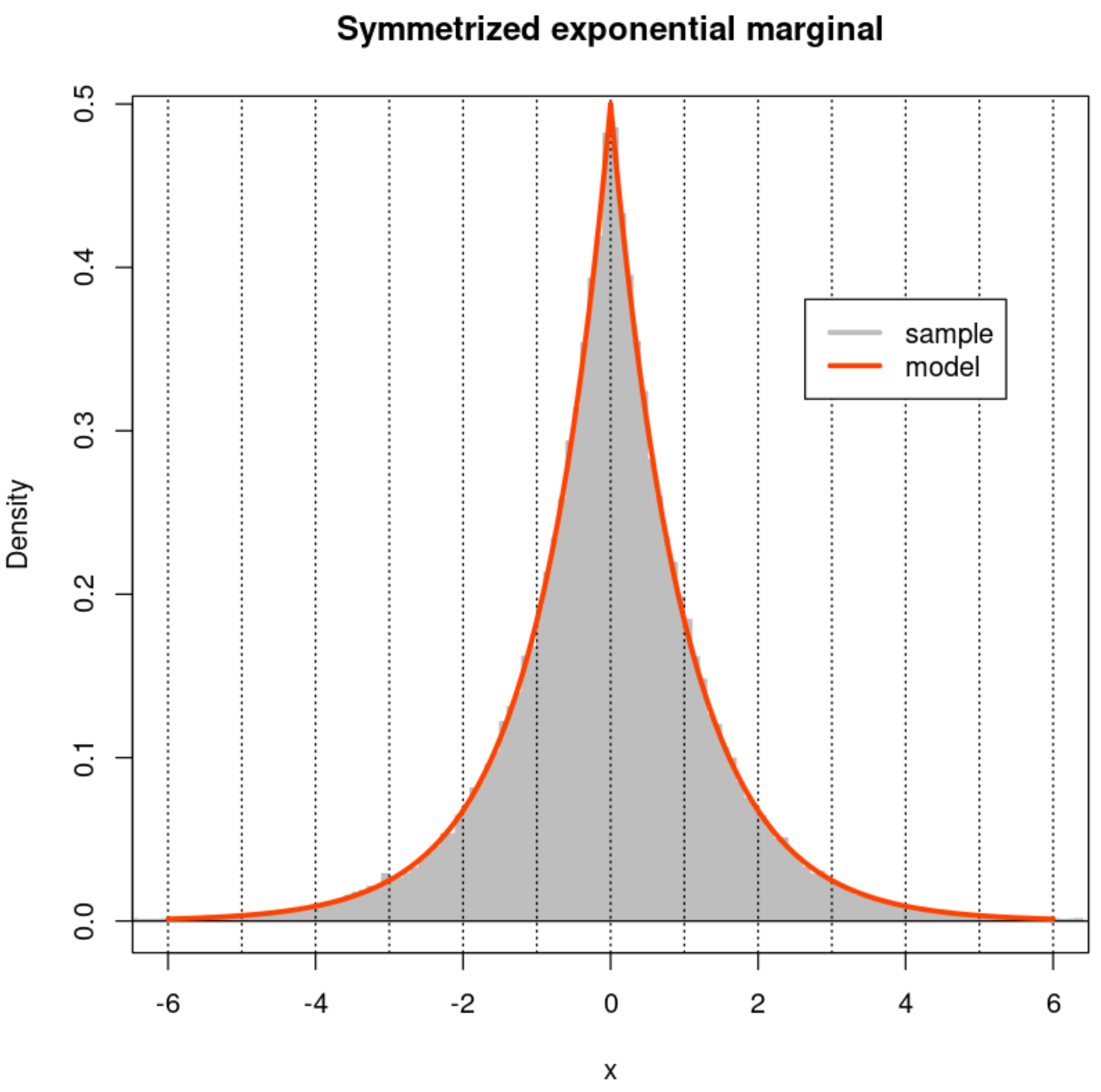


In [9]:

```
fine <- 101
par(mfrow=c(2,2))
xx <- (c(-50:49) + 0.05)/50
for (n in Ns) {
  X <- matrix(marginal(n*M),ncol=M,nrow=n)
  Z <- cor(X,method='p')
  hist(Z[row(Z) < col(Z)],nclass=fine,xlab="Correlation (r)",
       main=paste(n,"-vectors",sep=''),
       col='gray', border='gray', prob=TRUE, xlim=c(-1,1))
  lines(xx, dbeta((1+xx)/2,n/2-1,n/2-1)/2, col='orangered',lwd=3)
  abline(h=0) ; abline(v=c(-1:1),lty=3)
  box()
}
```

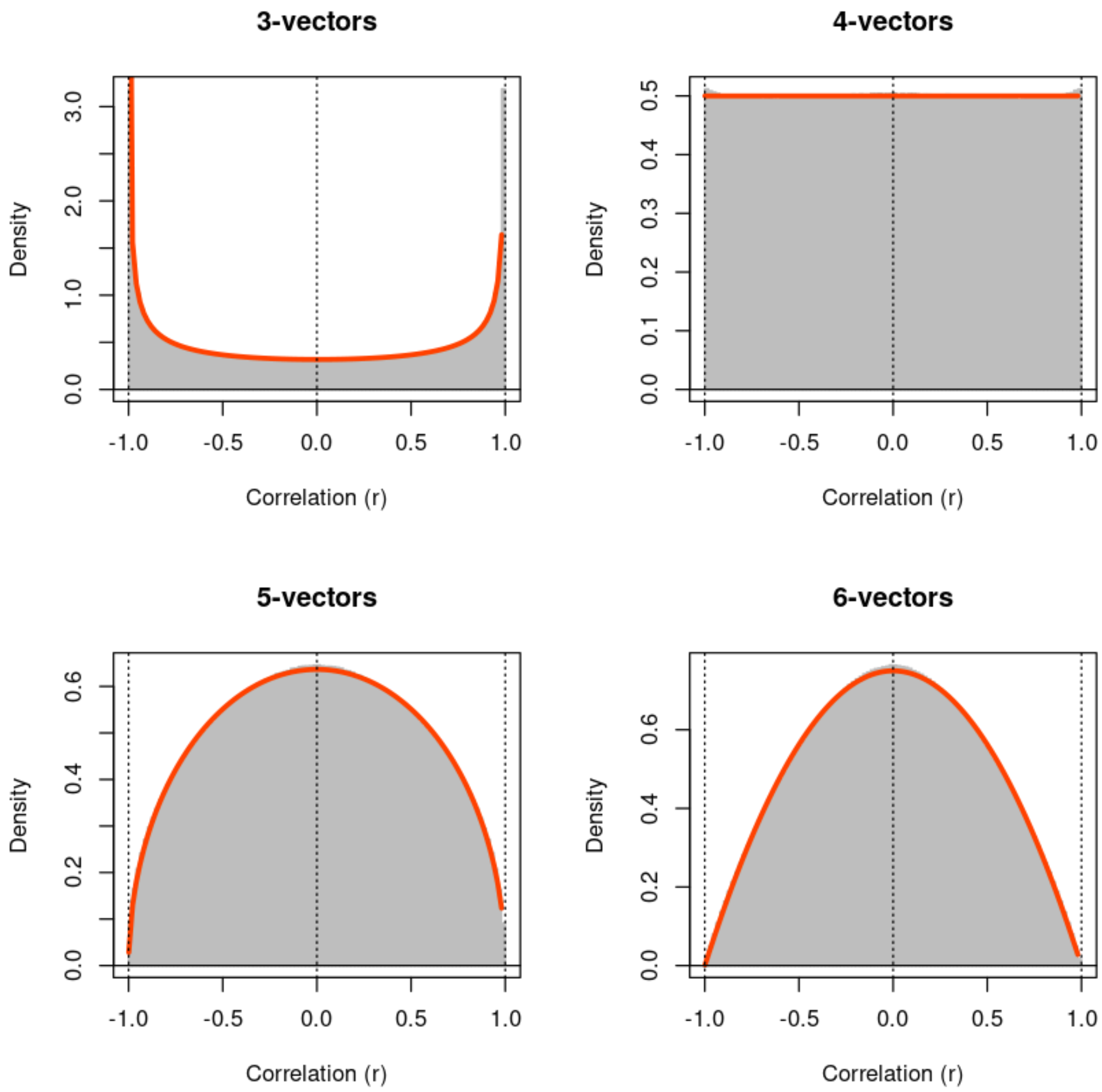

3-vectors
4-vectors
5-vectors
6-vectors
Density
Correlation (r)
-1.0
-0.5
0.0
0.5
1.0

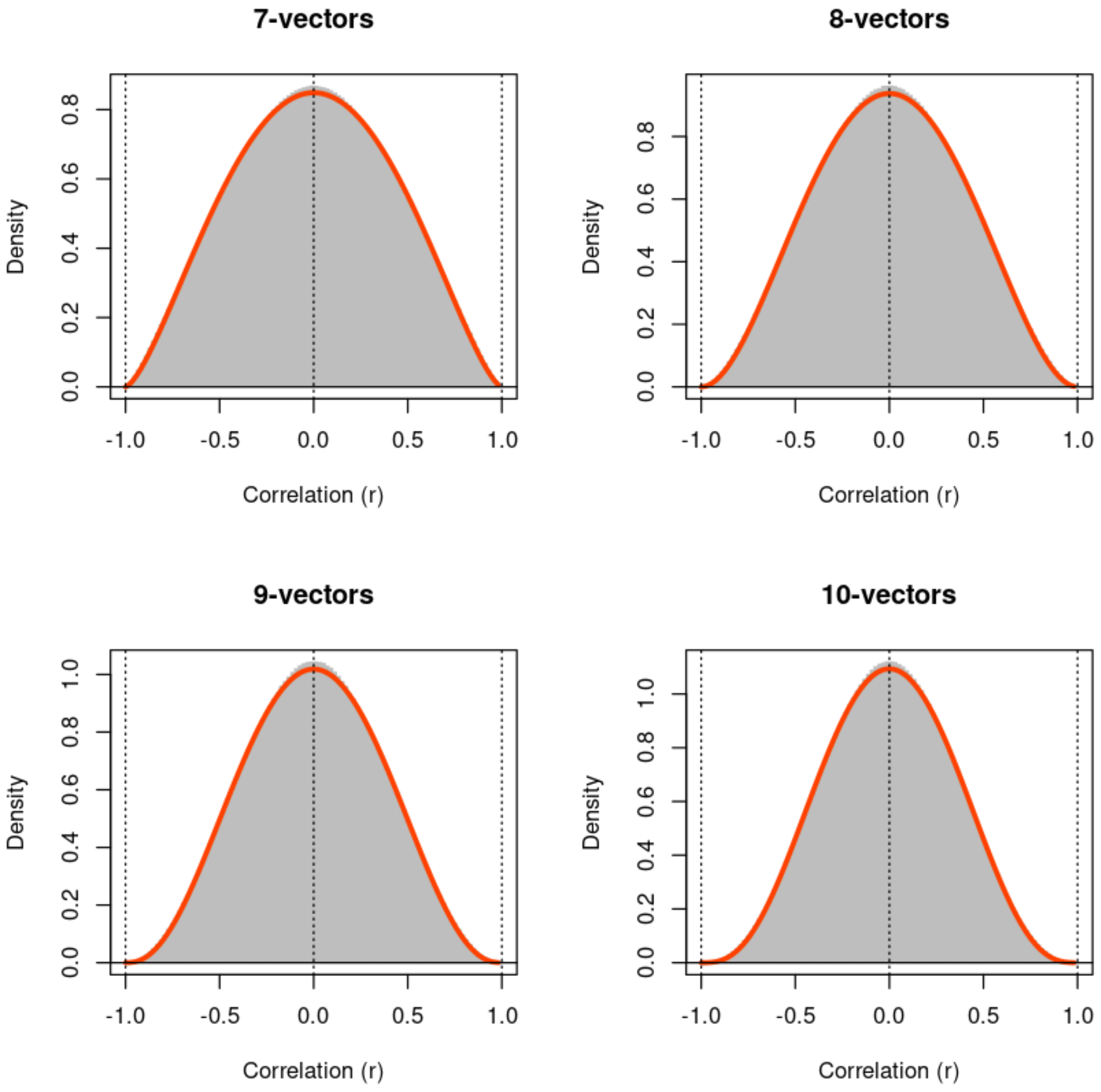

7-vectors
8-vectors
9-vectors
10-vectors
Density
Correlation (r)
-1.0
-0.5
0.0
0.5
1.0
0.2
0.4
0.6
0.8

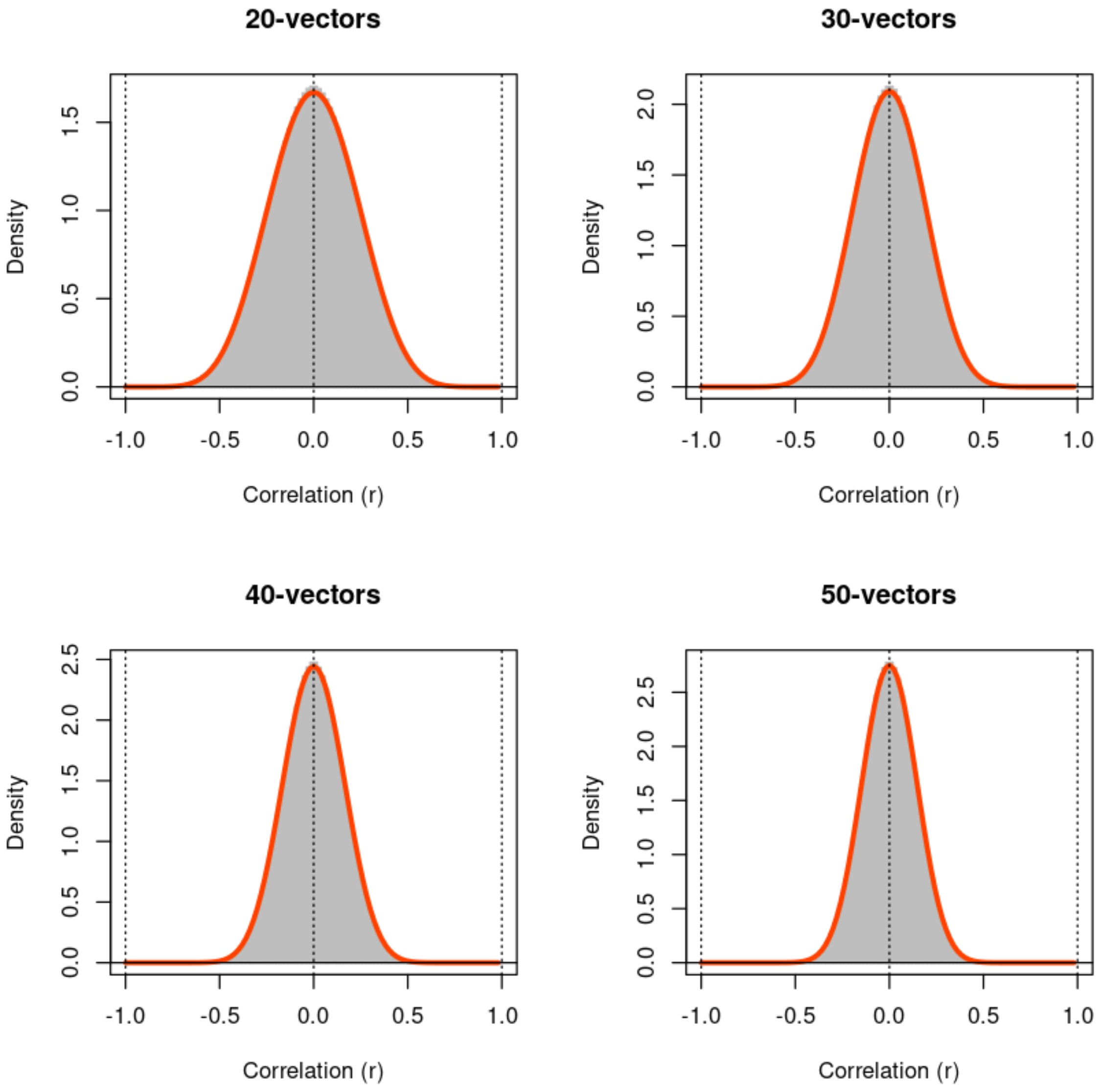
20-vectors
30-vectors
40-vectors
50-vectors
Density
Correlation (r)
-1.0
-0.5
0.0
0.5
1.0
1.5
2.0
2.5

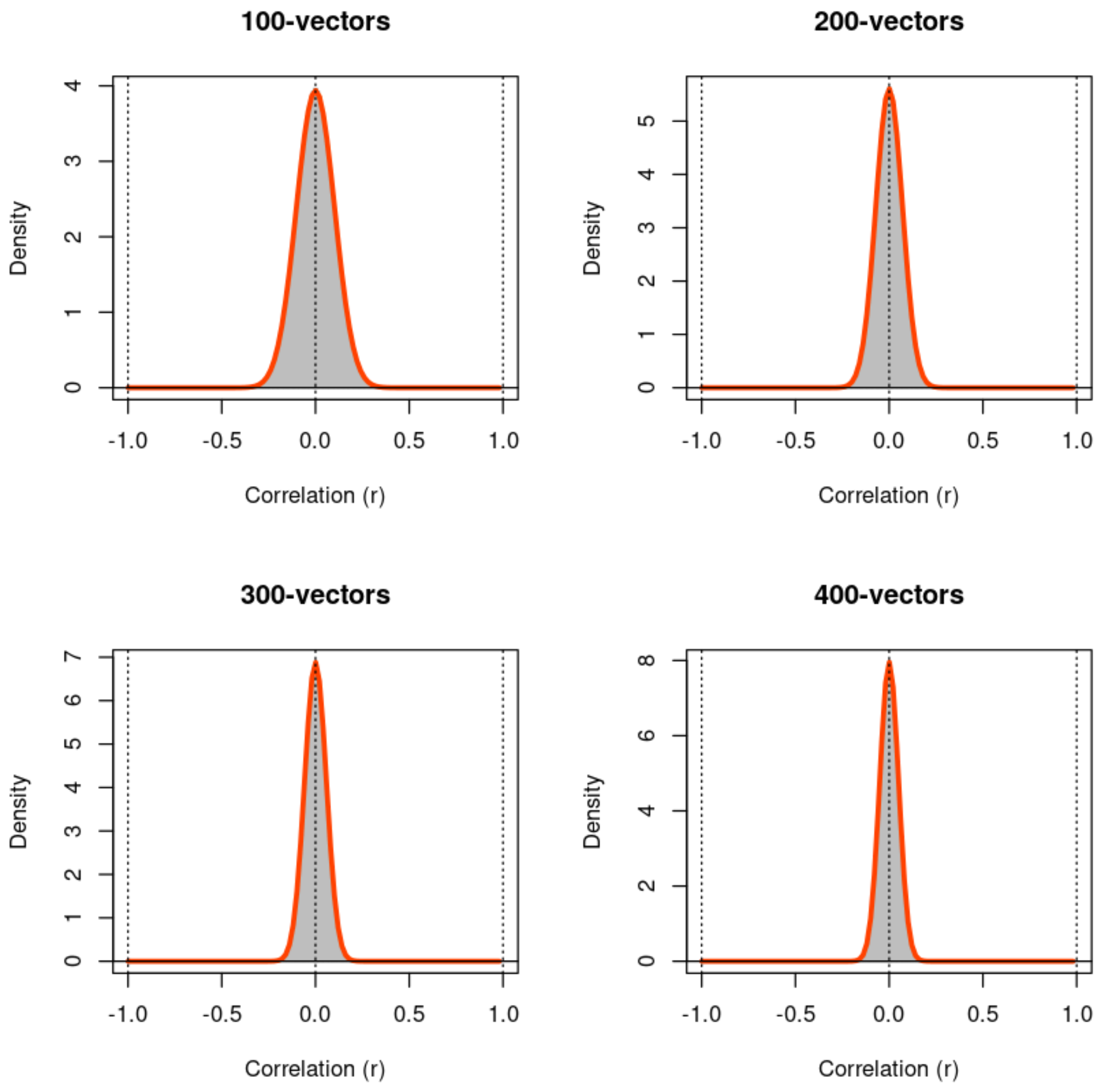


In [32]:
```
mu <- 2
marginal <- function(...) { mu - rexp(rate=1,...) }

xx <- c(-600:200)/100
xr <- range(xx) + c(-1,1)/2
hist(marginal(100000),main="Mean-minus-exponential marginal (mean=0)", xl
     col='gray', border='gray', prob=TRUE, xlim=xr, nclass=201)
abline(h=0); abline(v=c(round(min(xx)):round(max(xx))),lty=3)
abline(v=0)
lines(xx,dexp(2-xx),lwd=3,col='orangered')
legend(4,0.8,legend = c('sample','model'),col=c('gray','orangered'),bg='w
box()
```

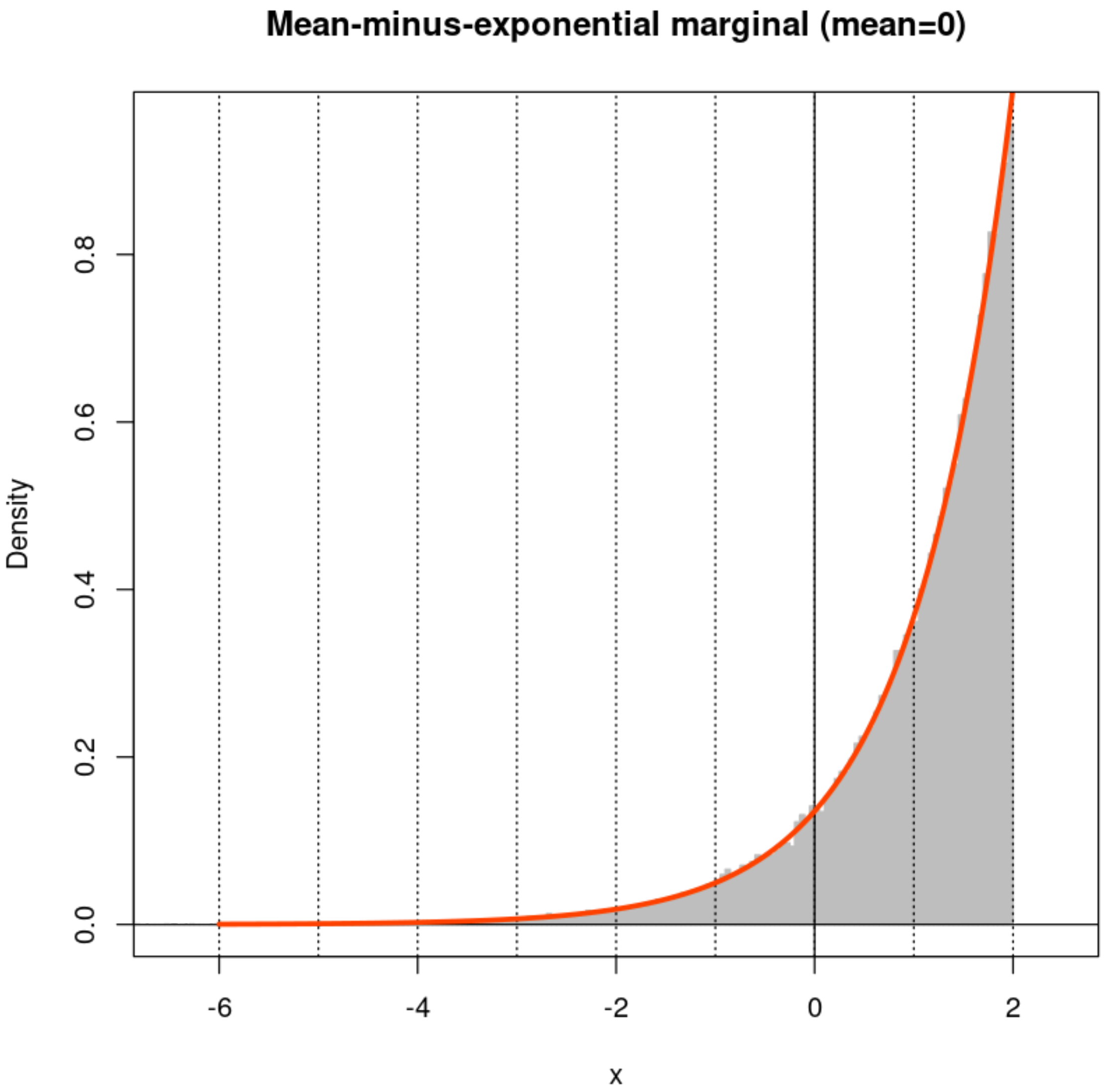


In [33]:

```
fine <- 101
par(mfrow=c(2,2))
xx <- (c(-50:49) + 0.05)/50
for (n in Ns) {
  X <- matrix(marginal(n*M),ncol=M,nrow=n)
  Z <- cor(X,method='p')
  hist(Z[row(Z) < col(Z)],nclass=fine,xlab="Correlation (r)",
       main=paste(n,"-vectors",sep=''),
       col='gray', border='gray', prob=TRUE, xlim=c(-1,1))
  lines(xx, dbeta((1+xx)/2,n/2-1,n/2-1)/2, col='orangered',lwd=3)
  abline(h=0) ; abline(v=c(-1:1),lty=3)
  box()
}
```

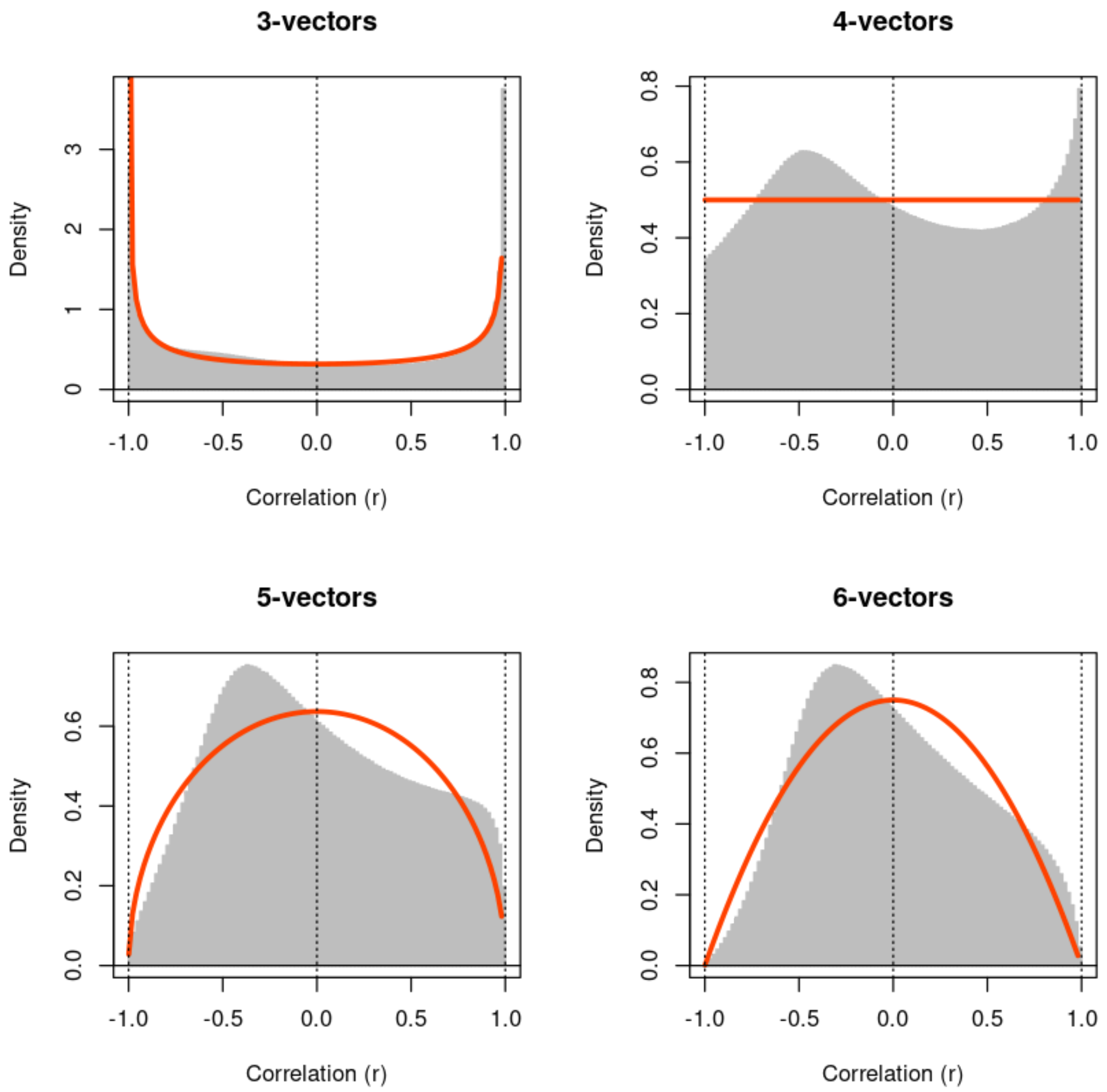
3-vectors
4-vectors
5-vectors
6-vectors
Density
Correlation (r)
-1.0
-0.5
0.0
0.5
1.0
0
1
2
3
0.0
0.2
0.4
0.6
0.8

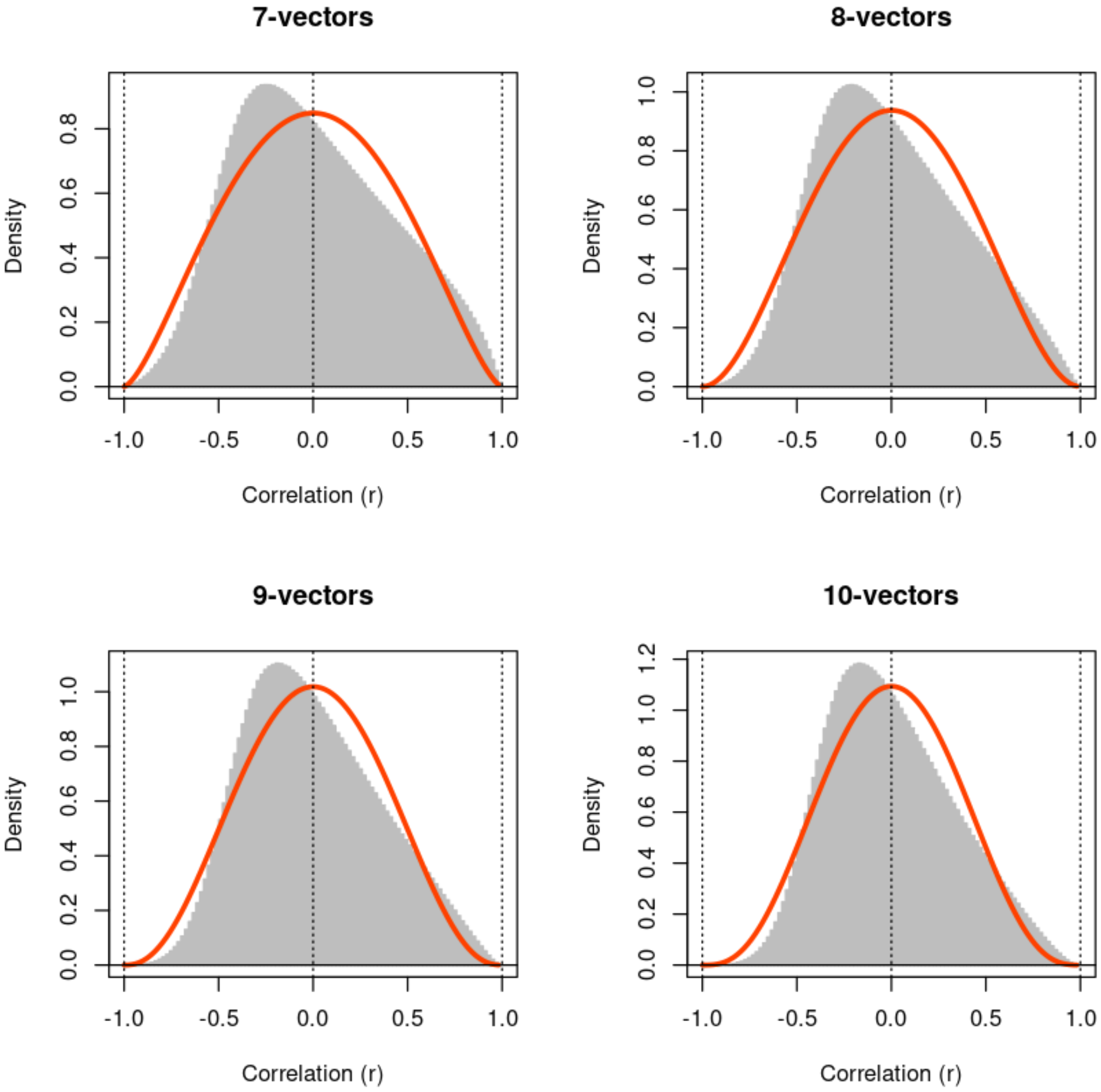
7-vectors
8-vectors
9-vectors
10-vectors
Density
Correlation (r)
-1.0
-0.5
0.0
0.5
1.0
0.0
0.2
0.4
0.6
0.8
1.0
1.2

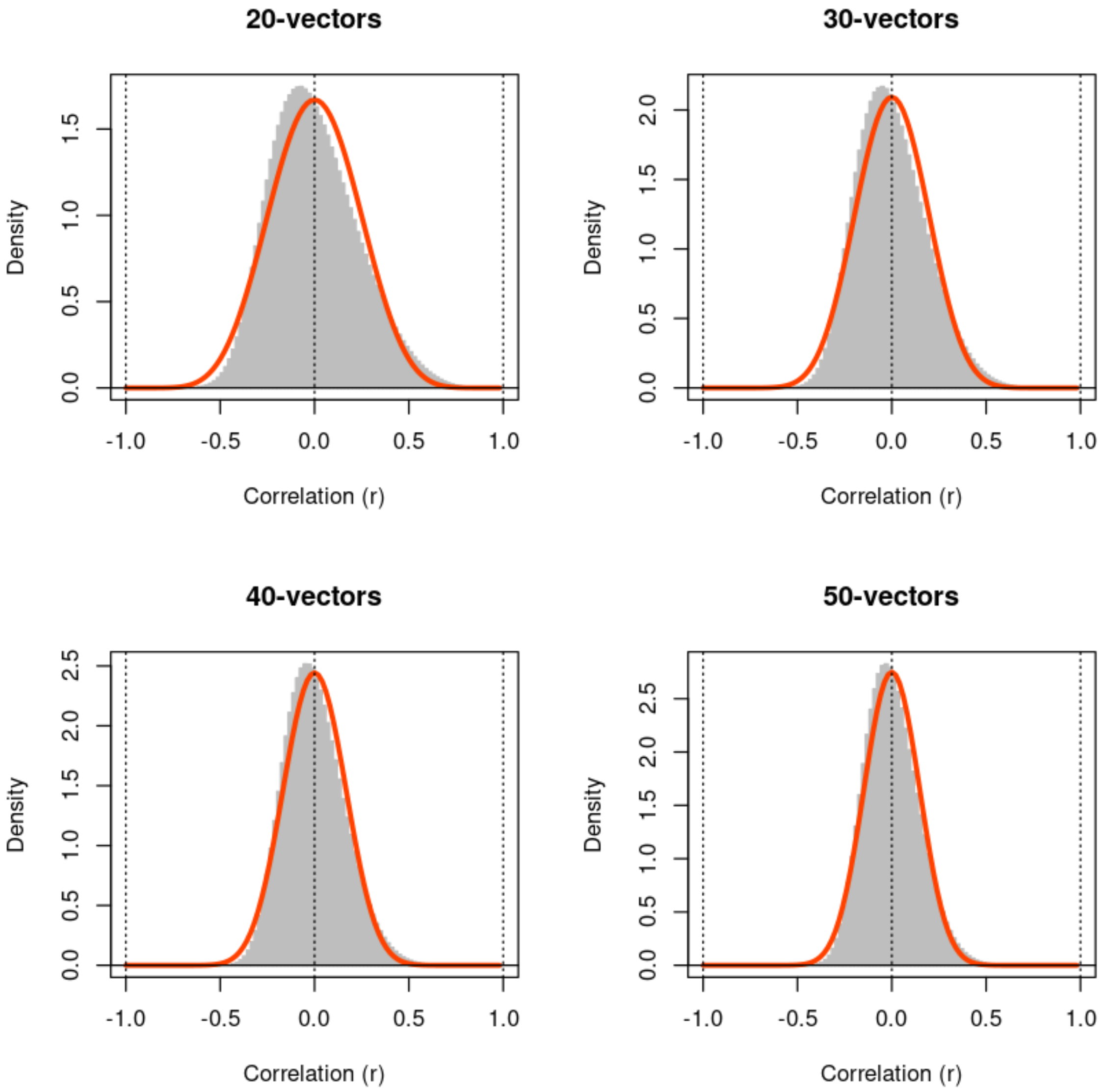
20-vectors
30-vectors
40-vectors
50-vectors
Density
Correlation (r)
-1.0
-0.5
0.0
0.5
1.0
1.5
2.0
2.5

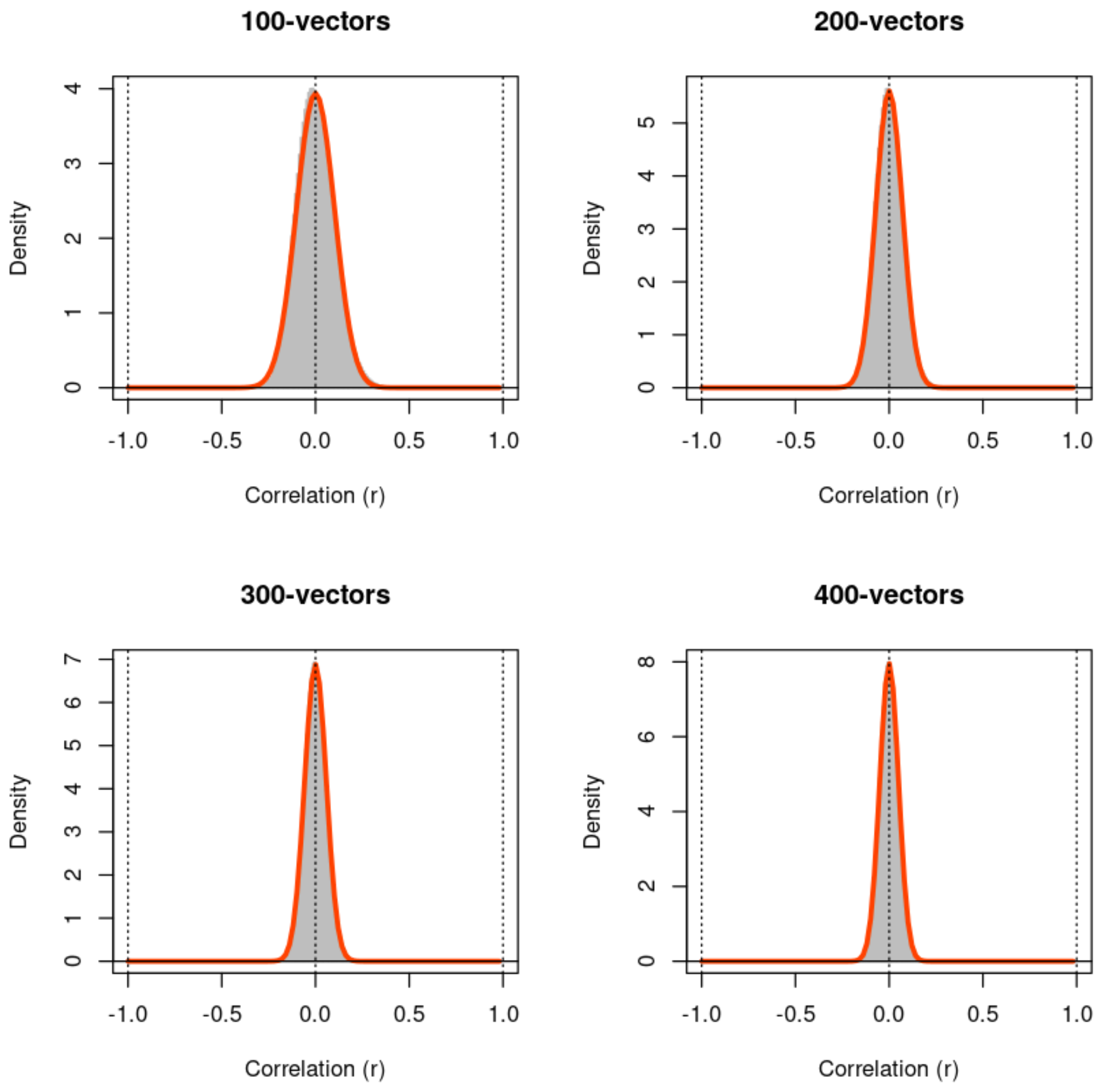


In [34]:
```
marginal <- function(...) { rt(..., df=3) }

x <- marginal(100000)
hist(x[abs(x) < 6],main="t-distributed marginal, df=3 (finite variance)",
     col='gray', border='gray', prob=TRUE, xlim=c(-6,6), nclass=300)
xx <- c(-600:600)/100
abline(h=0); abline(v=c(-6:6),lty=3)
lines(xx,dt(xx,df=3)/(1-2*pt(-6,df=3,lower.tail=TRUE)),lwd=3,col='oranger
box()
legend(4,0.3,legend = c('sample','model'),col=c('gray','orangered'),bg='w
```

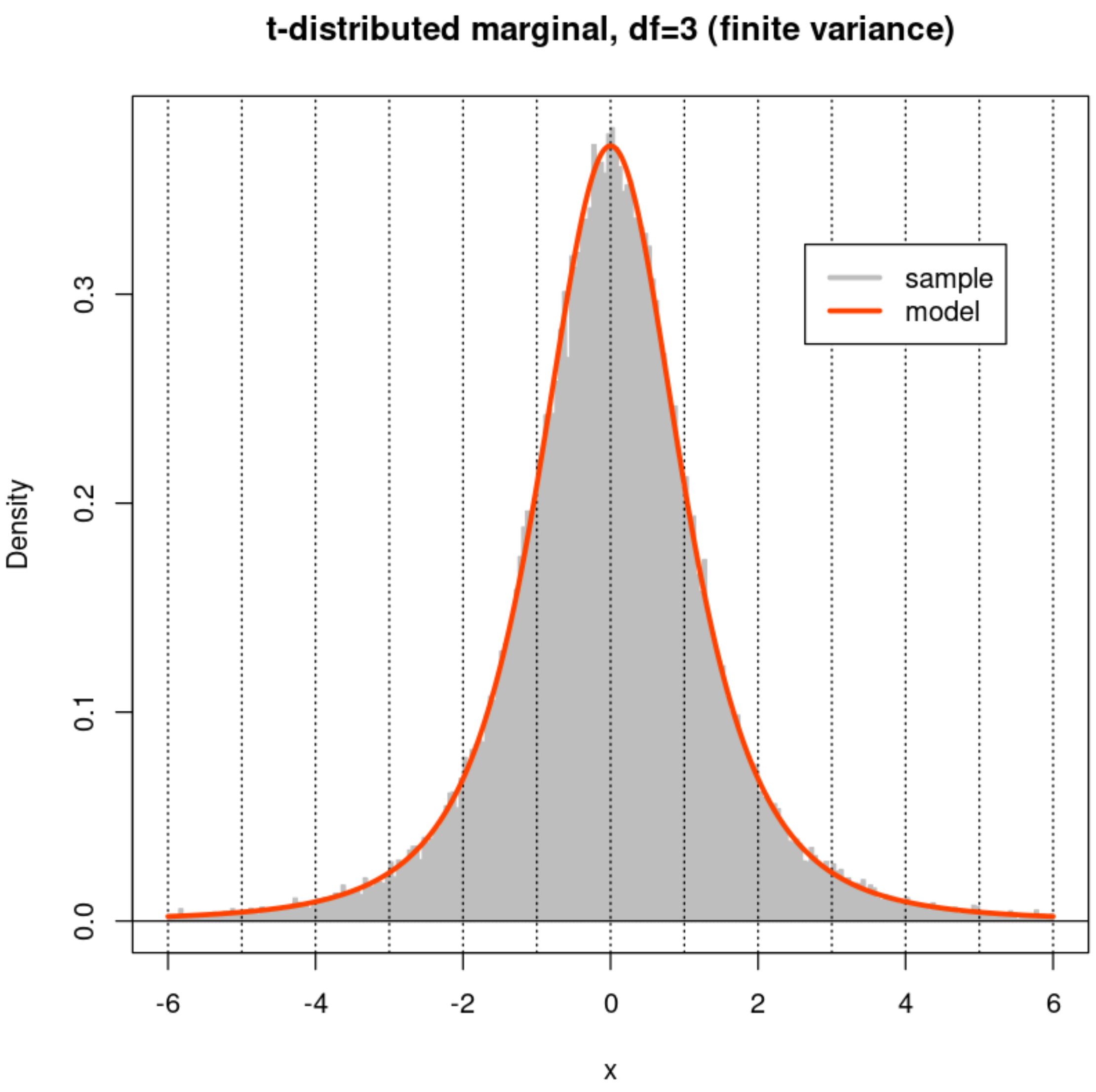


In [35]:

```
fine <- 101
par(mfrow=c(2,2))
xx <- (c(-50:49) + 0.05)/50
for (n in Ns) {
  X <- matrix(marginal(n*M),ncol=M,nrow=n)
  Z <- cor(X,method='p')
  hist(Z[row(Z) < col(Z)],nclass=fine,xlab="Correlation (r)",
       main=paste(n,"-vectors",sep=''),
       col='gray', border='gray', prob=TRUE, xlim=c(-1,1))
  lines(xx, dbeta((1+xx)/2,n/2-1,n/2-1)/2, col='orangered',lwd=3)
  abline(h=0) ; abline(v=c(-1:1),lty=3)
  box()
}
```

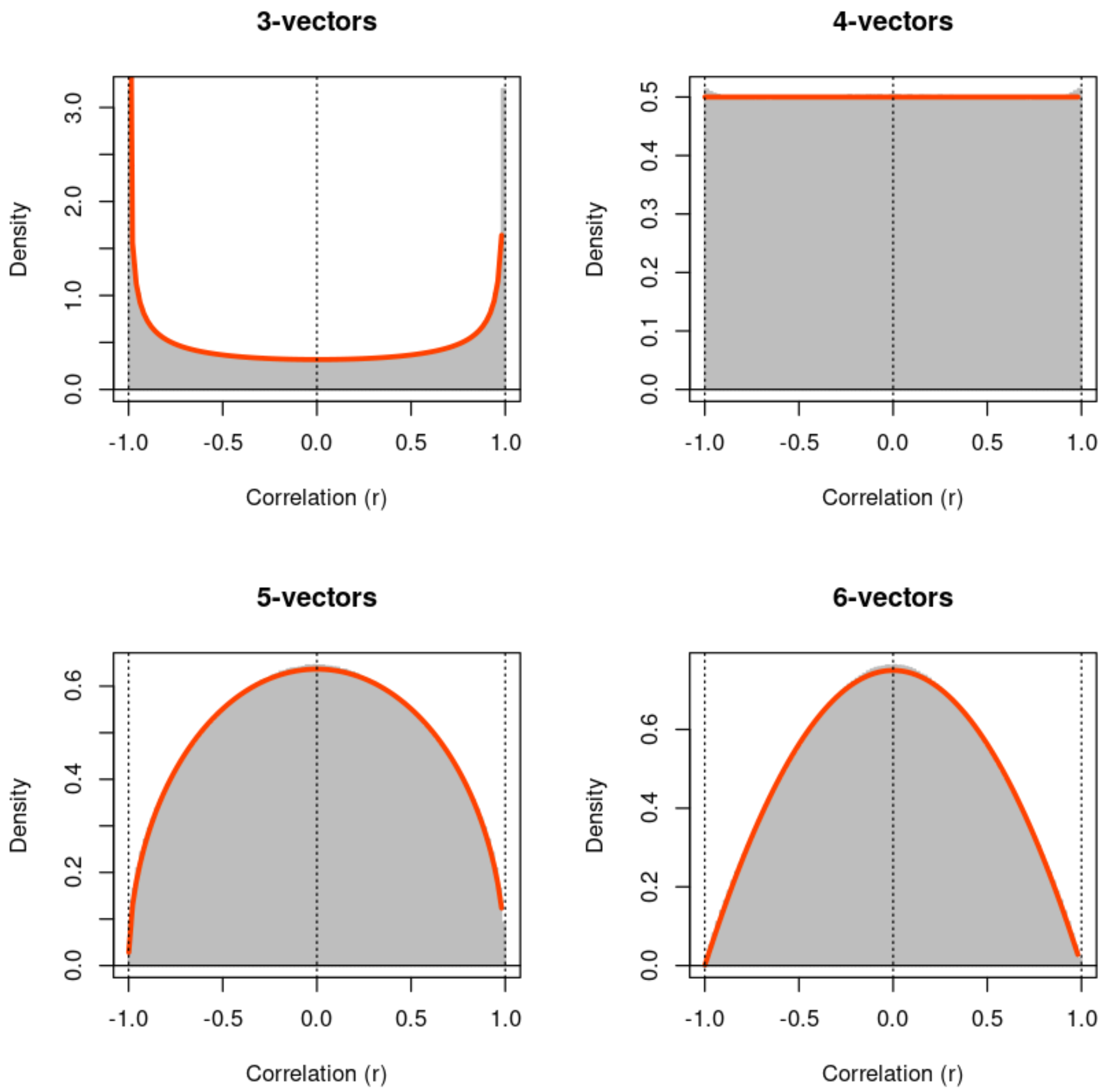
3-vectors
4-vectors
5-vectors
6-vectors
Density
Correlation (r)
-1.0
-0.5
0.0
0.5
1.0
0.0
1.0
2.0
3.0
0.1
0.2
0.3
0.4
0.5
0.6

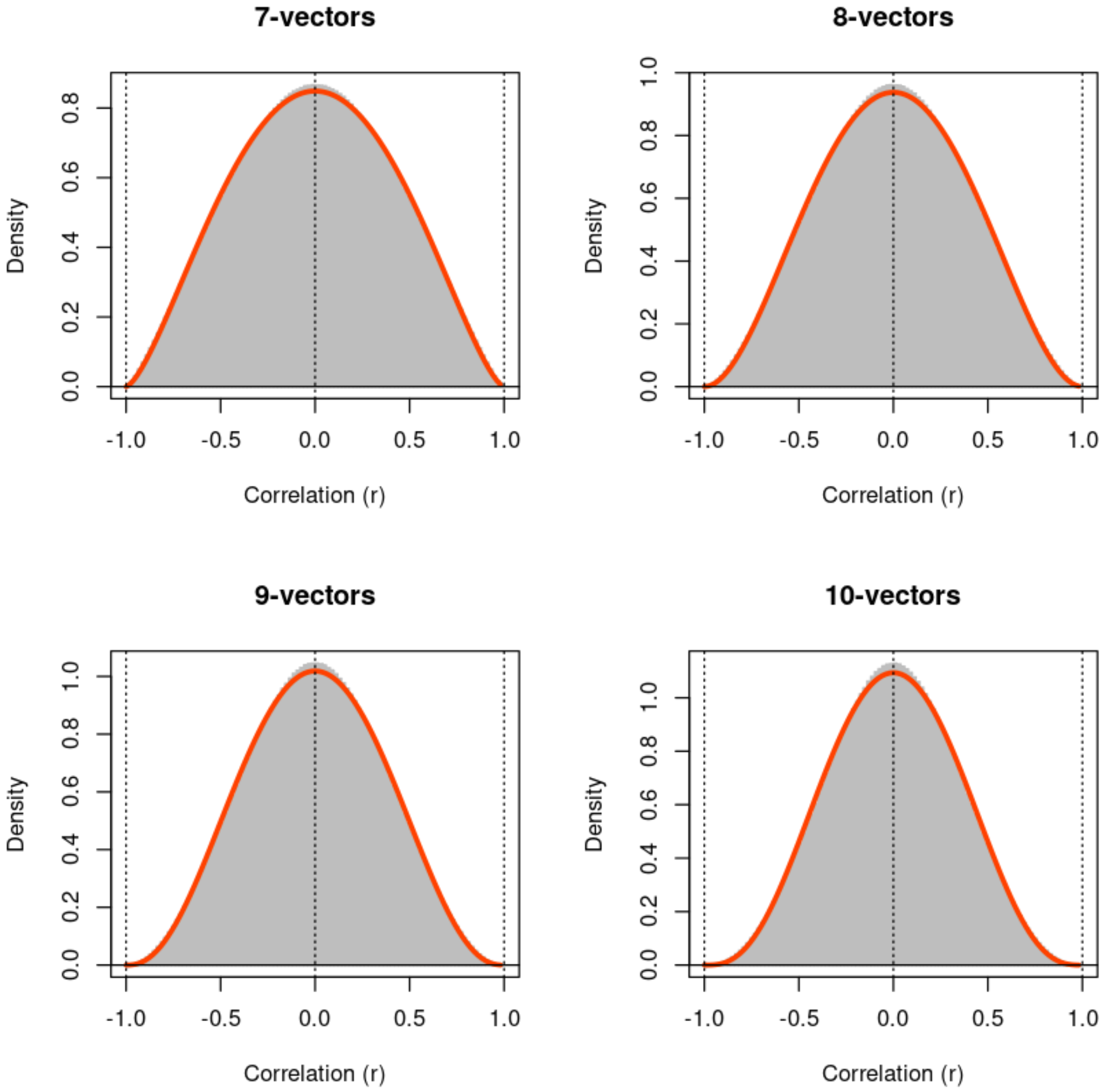
7-vectors
8-vectors
9-vectors
10-vectors
Density
Correlation (r)
-1.0
-0.5
0.0
0.5
1.0
0.2
0.4
0.6
0.8

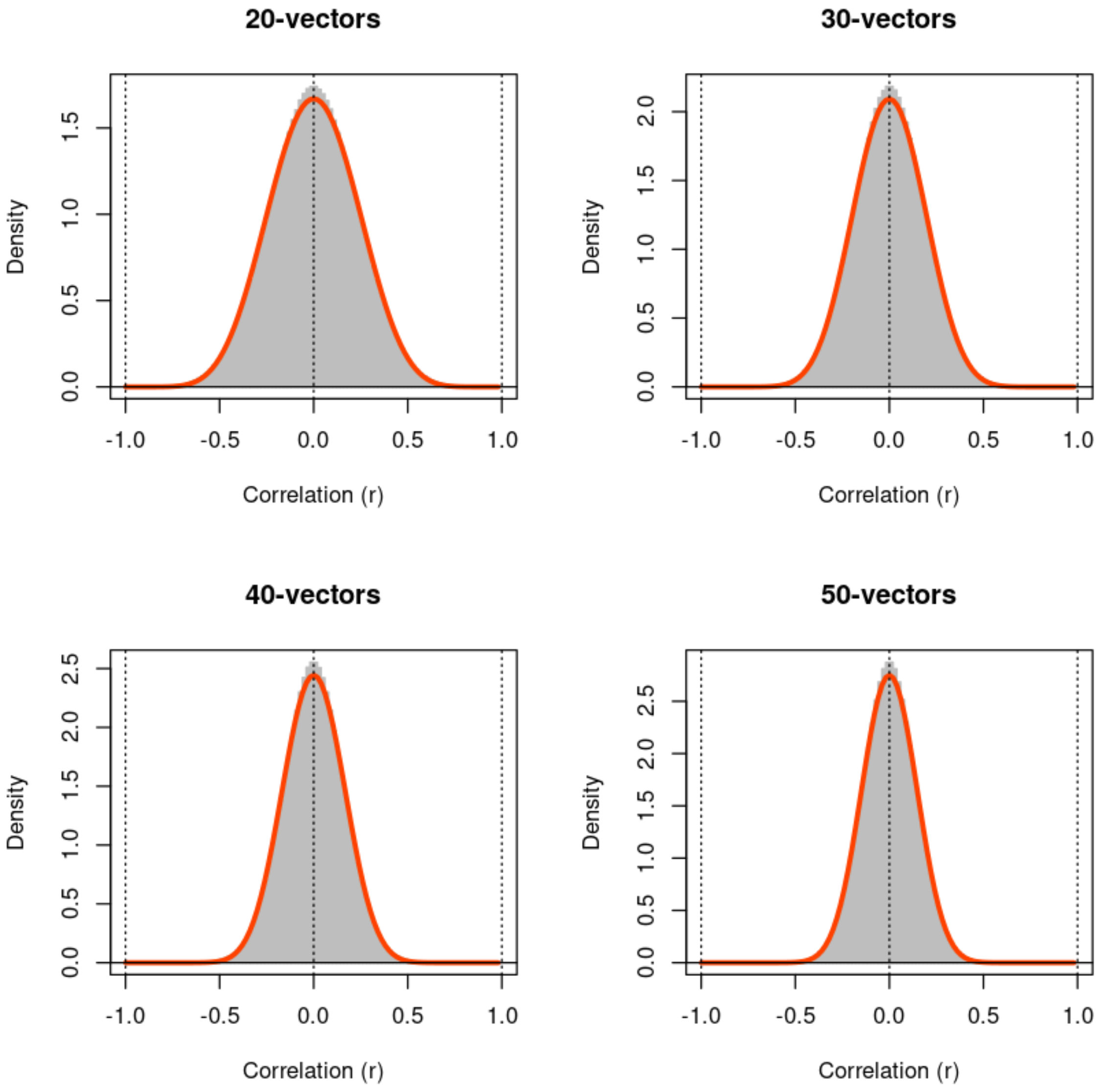
20-vectors
30-vectors
40-vectors
50-vectors
Density
Correlation (r)
-1.0
-0.5
0.0
0.5
1.0
1.5
2.0
2.5

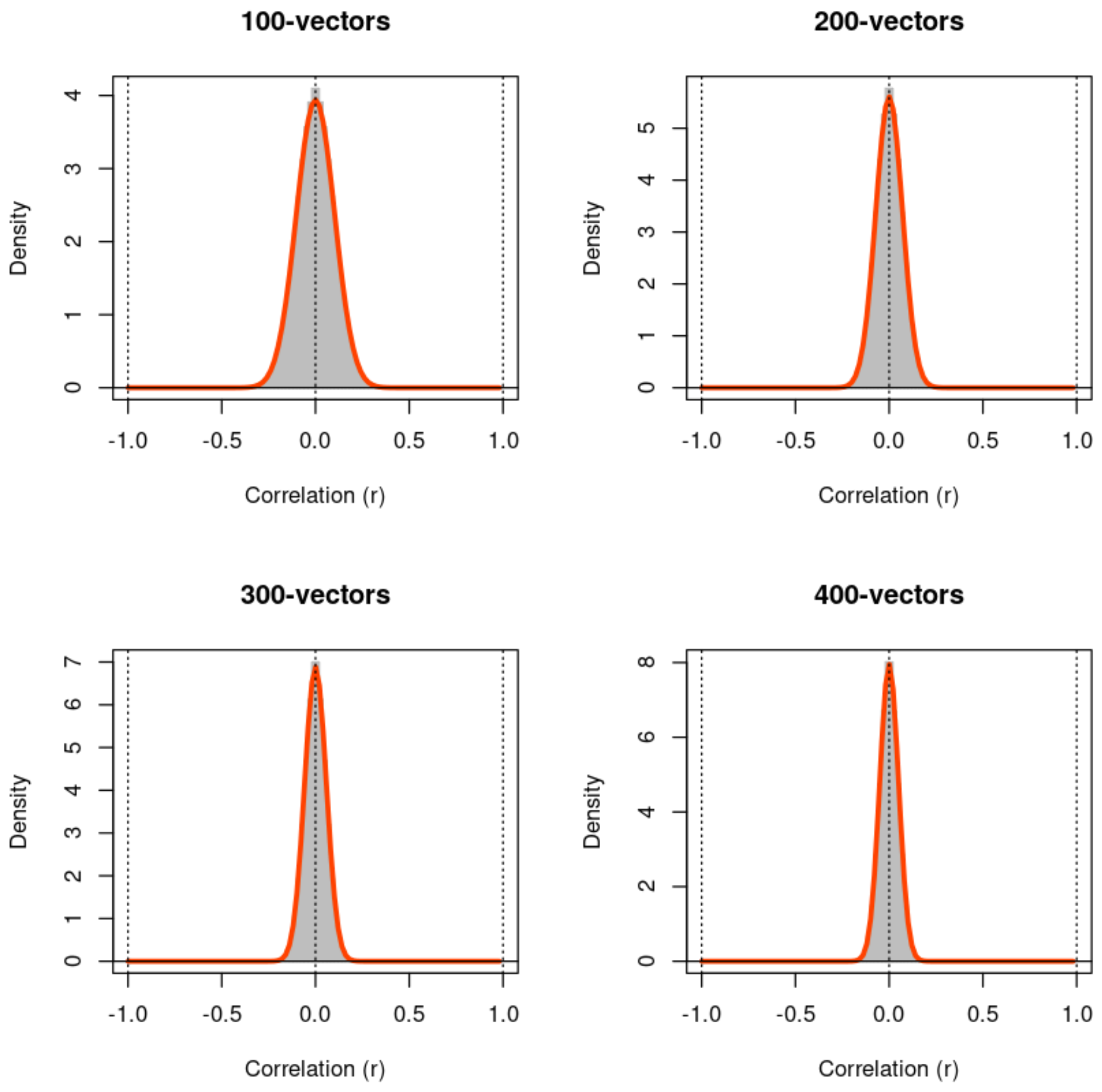


In [36]:

```
marginal <- function(...) { rt(..., df=2) }

x <- marginal(100000)
hist(x[abs(x) < 6],main="t-distributed marginal, df=2 (nonfinite variance
     col='gray', border='gray', prob=TRUE, xlim=c(-6,6), nclass=300)
xx <- c(-600:600)/100
abline(h=0); abline(v=c(-6:6),lty=3)
lines(xx,dt(xx,df=2)/(1-2*pt(-6,df=2,lower.tail=TRUE)),lwd=3,col='oranger
legend(4,0.25,legend = c('sample','model'),col=c('gray','orangered'),bg='
box()
```

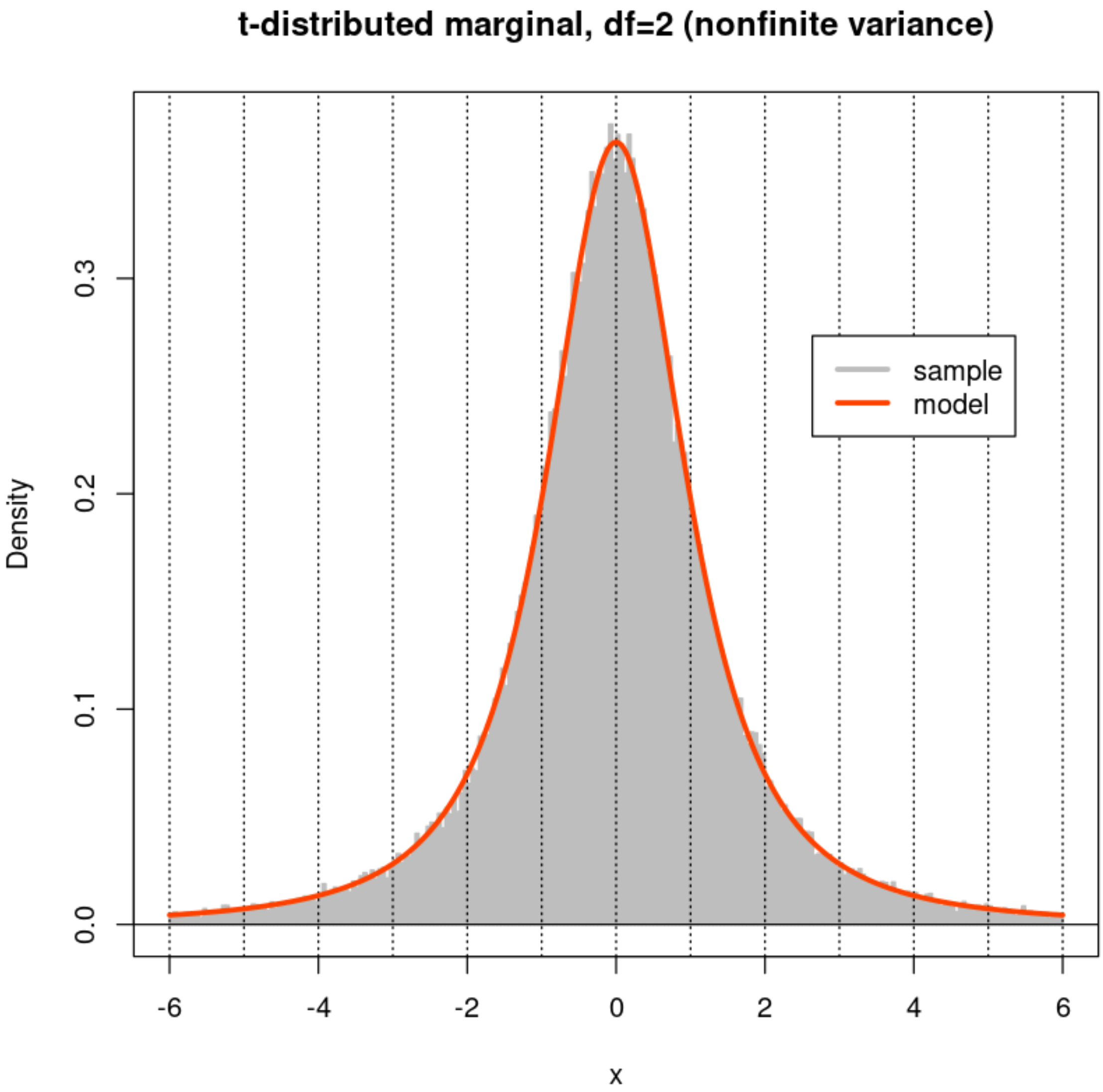


In [37]:
```
fine <- 101
par(mfrow=c(2,2))
xx <- (c(-50:49) + 0.05)/50
for (n in Ns) {
  X <- matrix(marginal(n*M),ncol=M,nrow=n)
  Z <- cor(X,method='p')
  hist(Z[row(Z) < col(Z)],nclass=fine,xlab="Correlation (r)",
       main=paste(n,"-vectors",sep=''),
       col='gray', border='gray', prob=TRUE, xlim=c(-1,1))
  lines(xx, dbeta((1+xx)/2,n/2-1,n/2-1)/2, col='orangered',lwd=3)
  abline(h=0) ; abline(v=c(-1:1),lty=3)
  box()
}
```

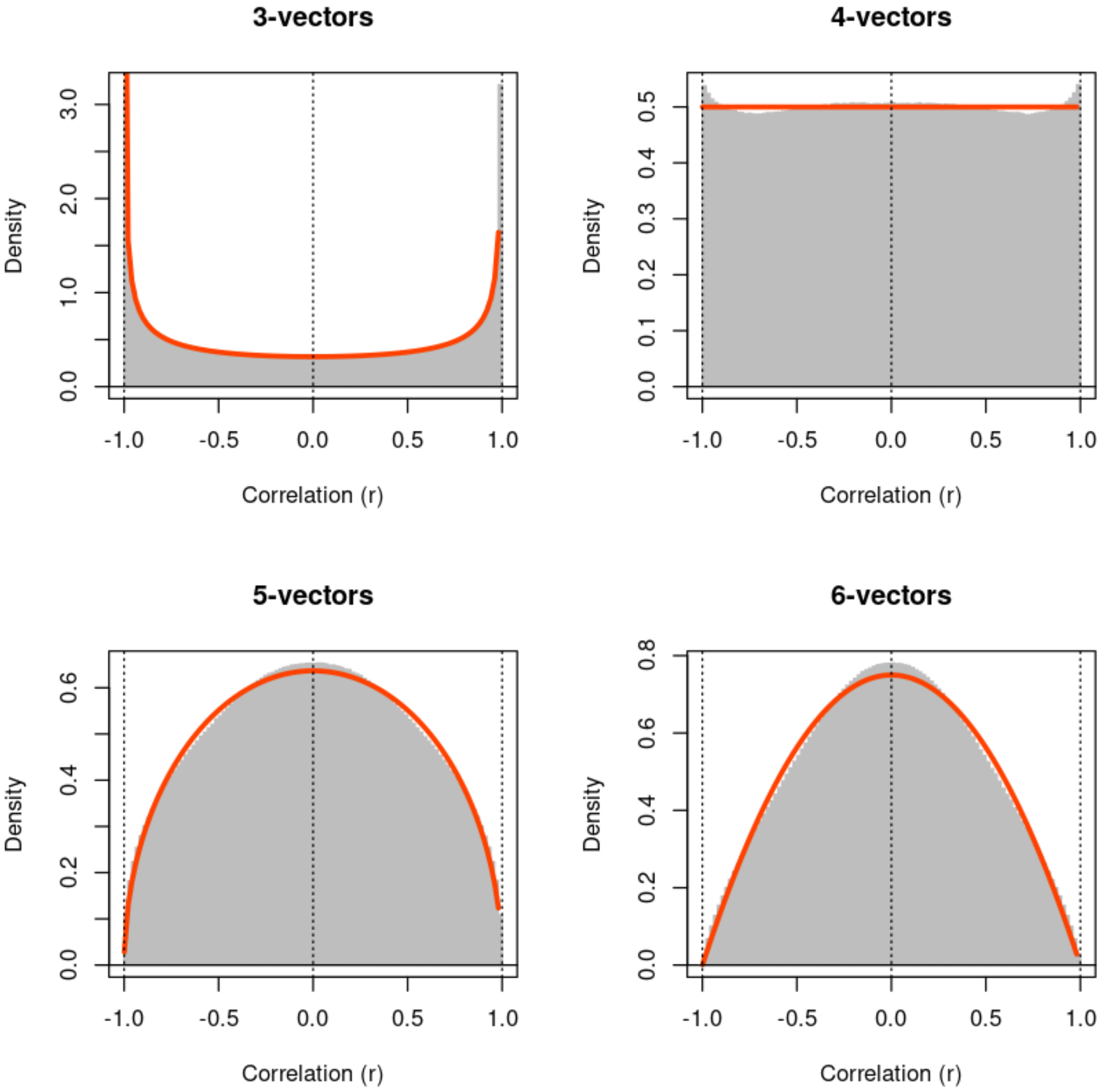
3-vectors
4-vectors
5-vectors
6-vectors
Density
Correlation (r)
-1.0
-0.5
0.0
0.5
1.0
0.0
1.0
2.0
3.0
0.1
0.2
0.3
0.4
0.5
0.6
0.8

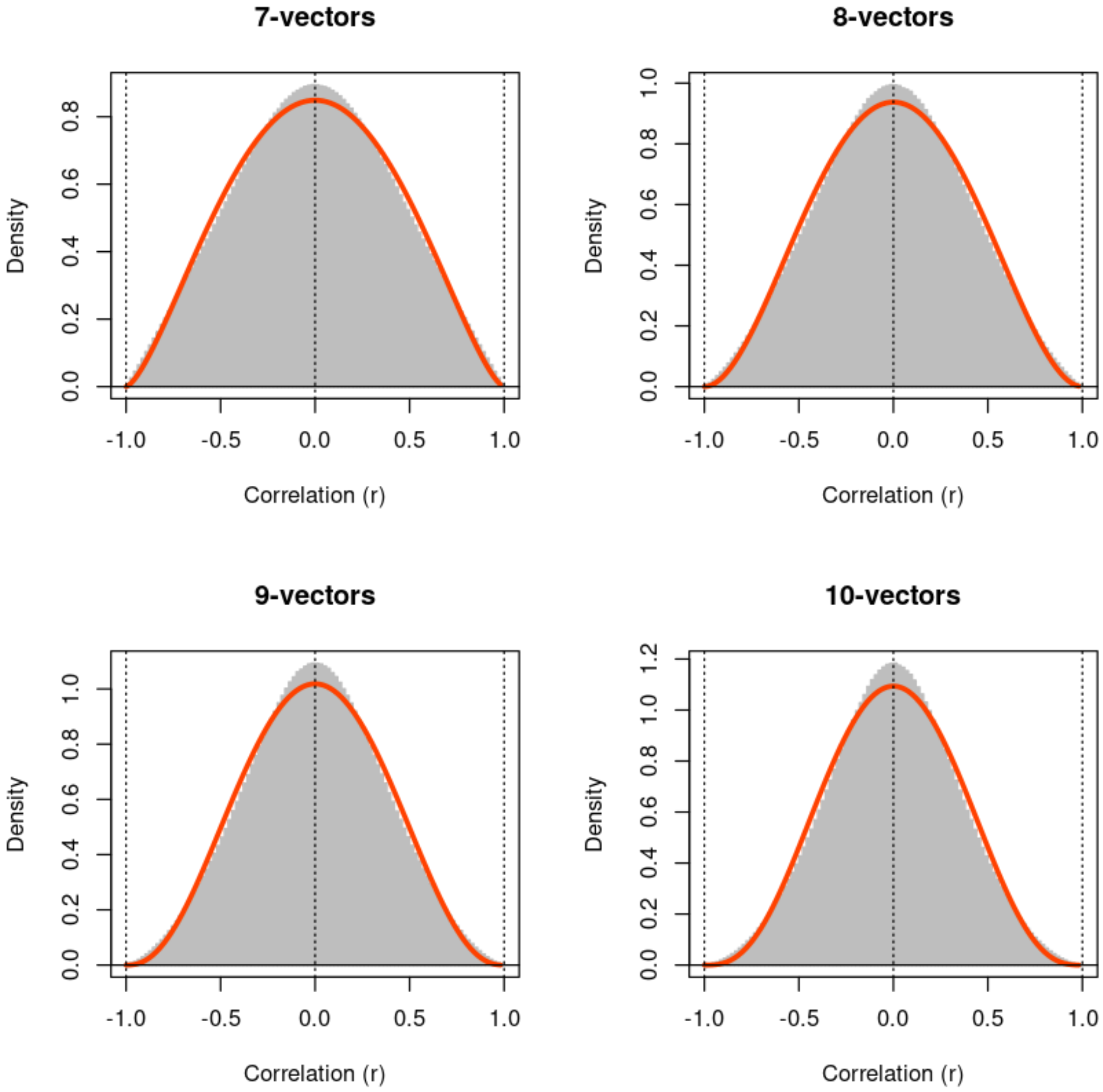

7-vectors
8-vectors
9-vectors
10-vectors
Density
Correlation (r)
-1.0
-0.5
0.0
0.5
1.0

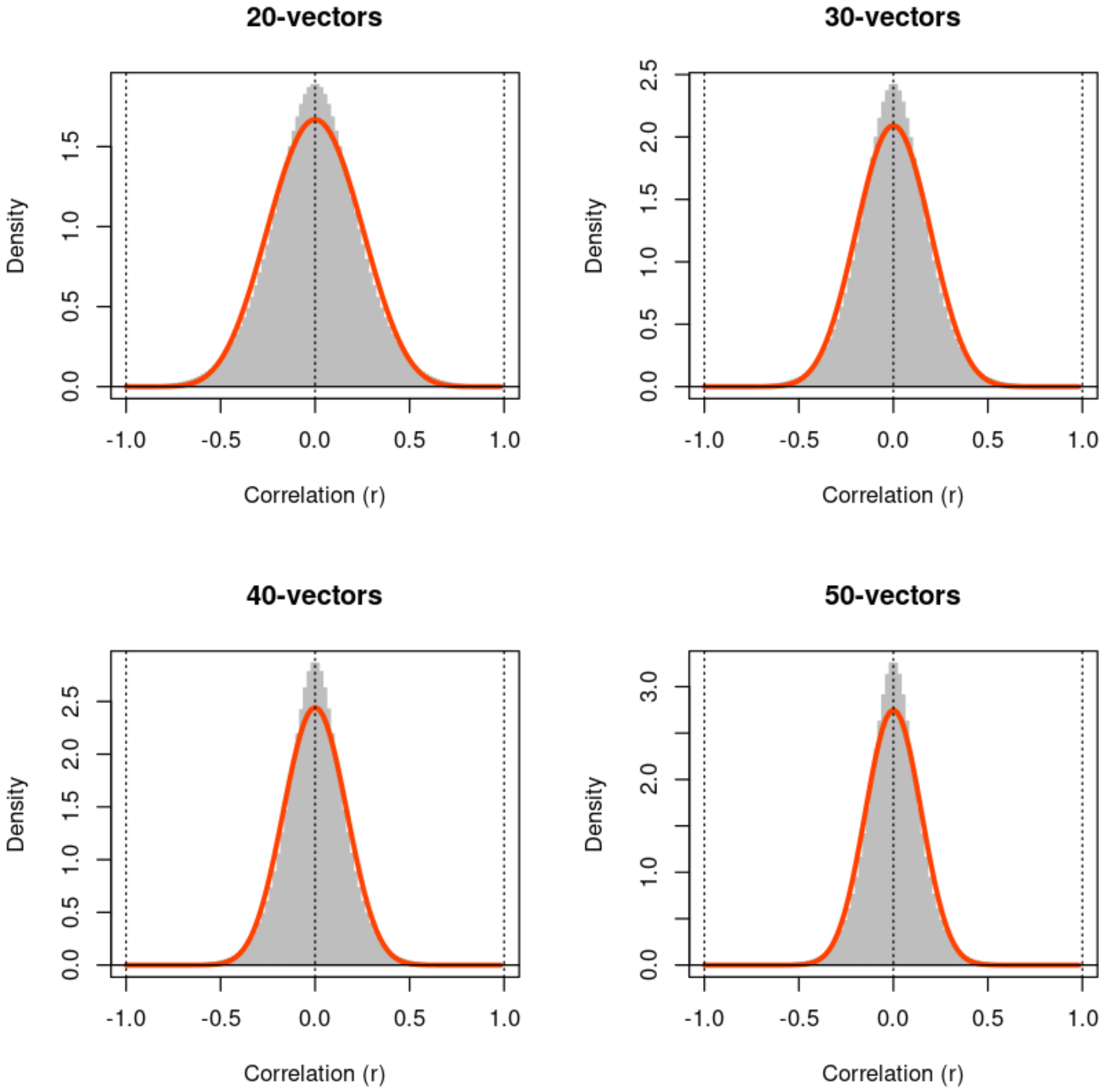
20-vectors
Density
0.0
0.5
1.0
1.5
-1.0
-0.5
0.0
0.5
1.0
Correlation (r)
30-vectors
Density
0.0
0.5
1.0
1.5
2.0
2.5
-1.0
-0.5
0.0
0.5
1.0
Correlation (r)
40-vectors
Density
0.0
0.5
1.0
1.5
2.0
2.5
-1.0
-0.5
0.0
0.5
1.0
Correlation (r)
50-vectors
Density
0.0
1.0
2.0
3.0
-1.0
-0.5
0.0
0.5
1.0
Correlation (r)

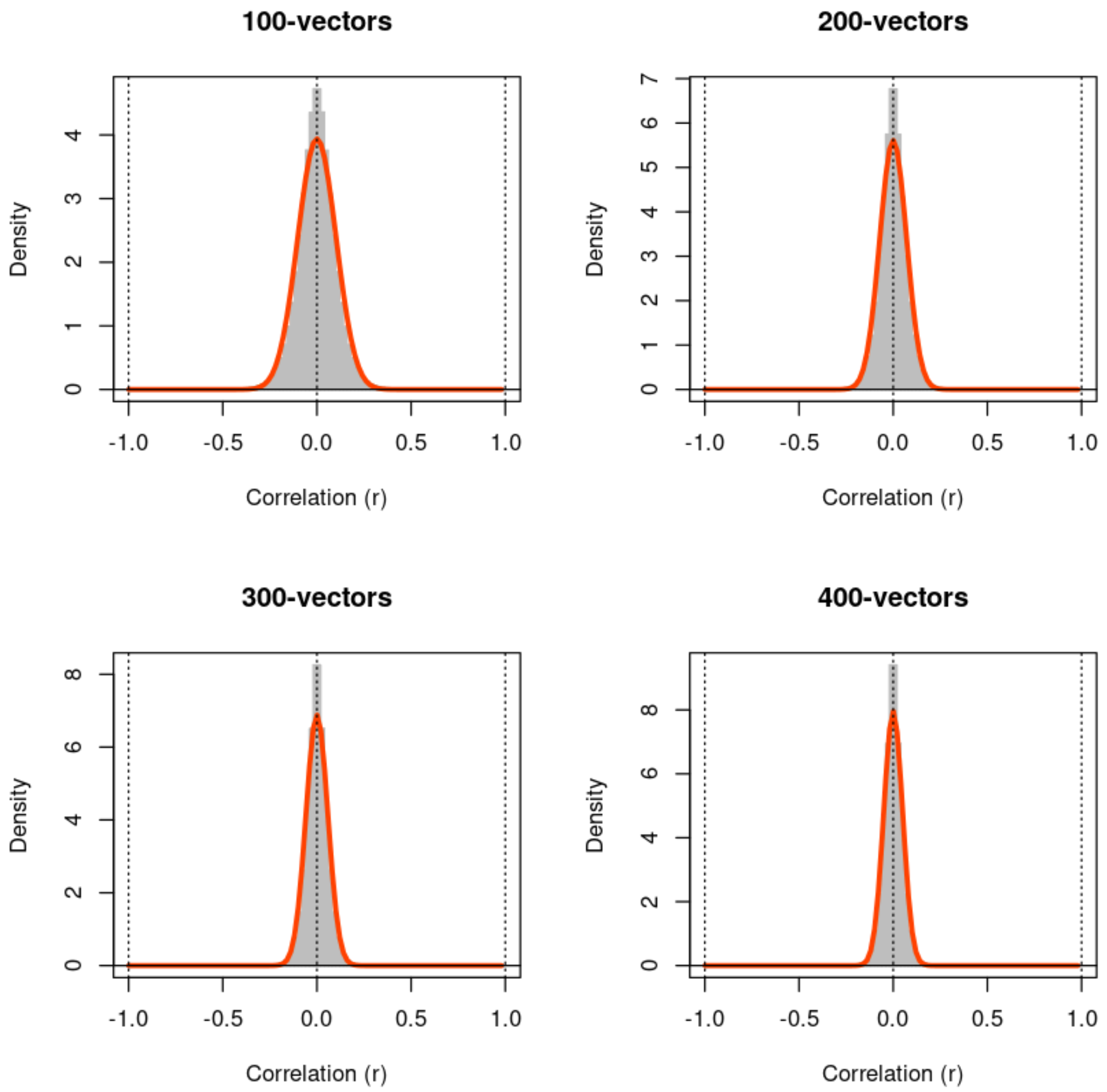


In [38]:
```
marginal <- function(...) { rt(..., df=1) }
x <- marginal(100000)
hist(x[abs(x) < 6],main="t-distributed marginal, df=1 (nonfinite variance
     col='gray', border='gray', prob=TRUE, xlim=c(-6,6), nclass=101)
xx <- c(-6000:6000)/1000
abline(h=0); abline(v=c(-6:6),lty=3)
lines(xx,dt(xx,df=1)/(1-2*pt(-6,df=1,lower.tail=TRUE)),lwd=3,col='oranger
legend(4,0.25,legend = c('sample','model'),col=c('gray','orangered'),bg='
box()
```

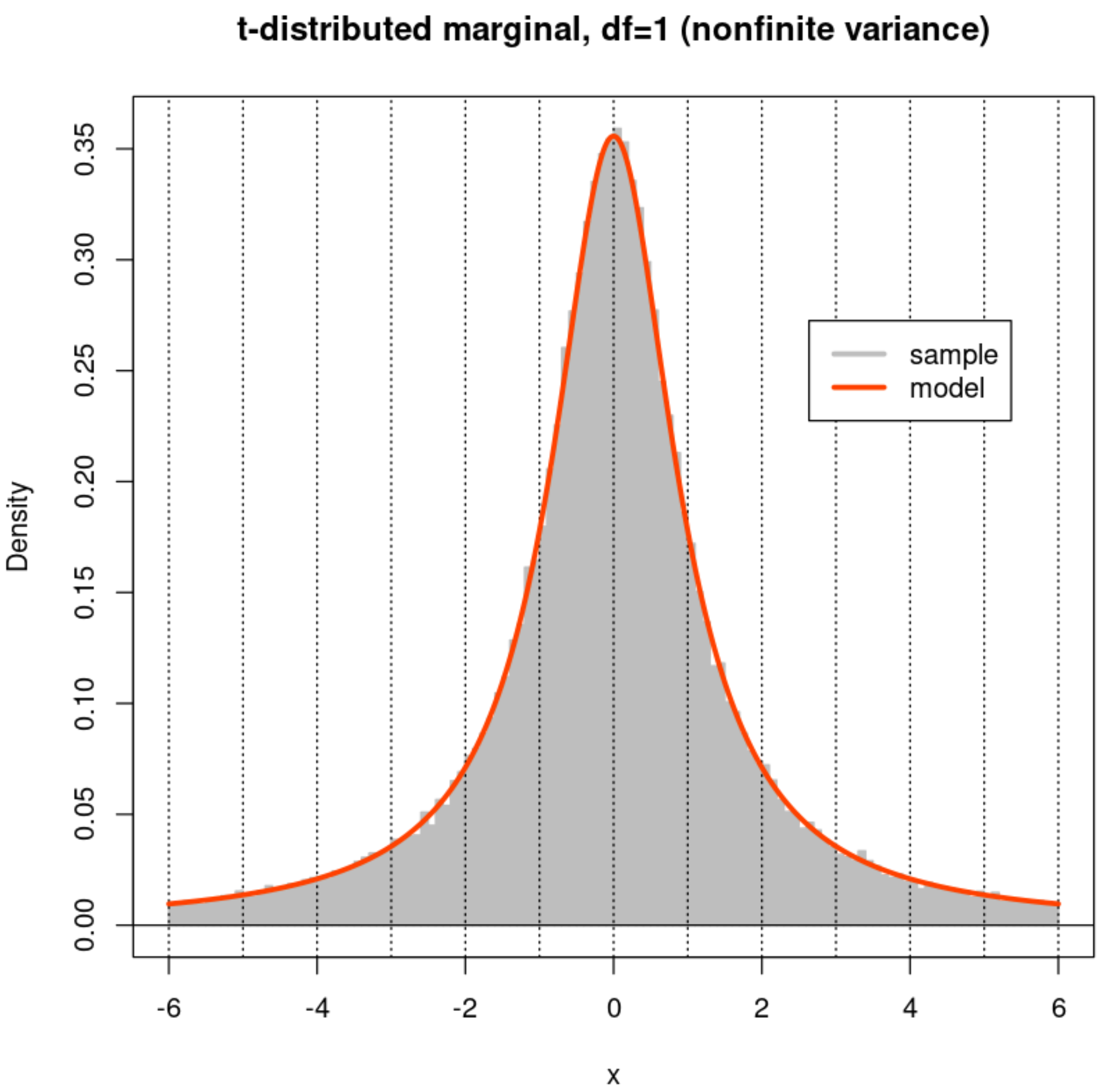


```
In [39]: fine <- 101
par(mfrow=c(2,2))
xx <- (c(-50:49) + 0.05)/50
for (n in Ns) {
  X <- matrix(marginal(n*M),ncol=M,nrow=n)
  Z <- cor(X,method='p')
  hist(Z[row(Z) < col(Z)],nclass=fine,xlab="Correlation (r)",
       main=paste(n,"-vectors",sep=''),
       col='gray', border='gray', prob=TRUE, xlim=c(-1,1))
  lines(xx, dbeta((1+xx)/2,n/2-1,n/2-1)/2, col='orangered',lwd=3)
  abline(h=0) ; abline(v=c(-1:1),lty=3)
  box()
}
```

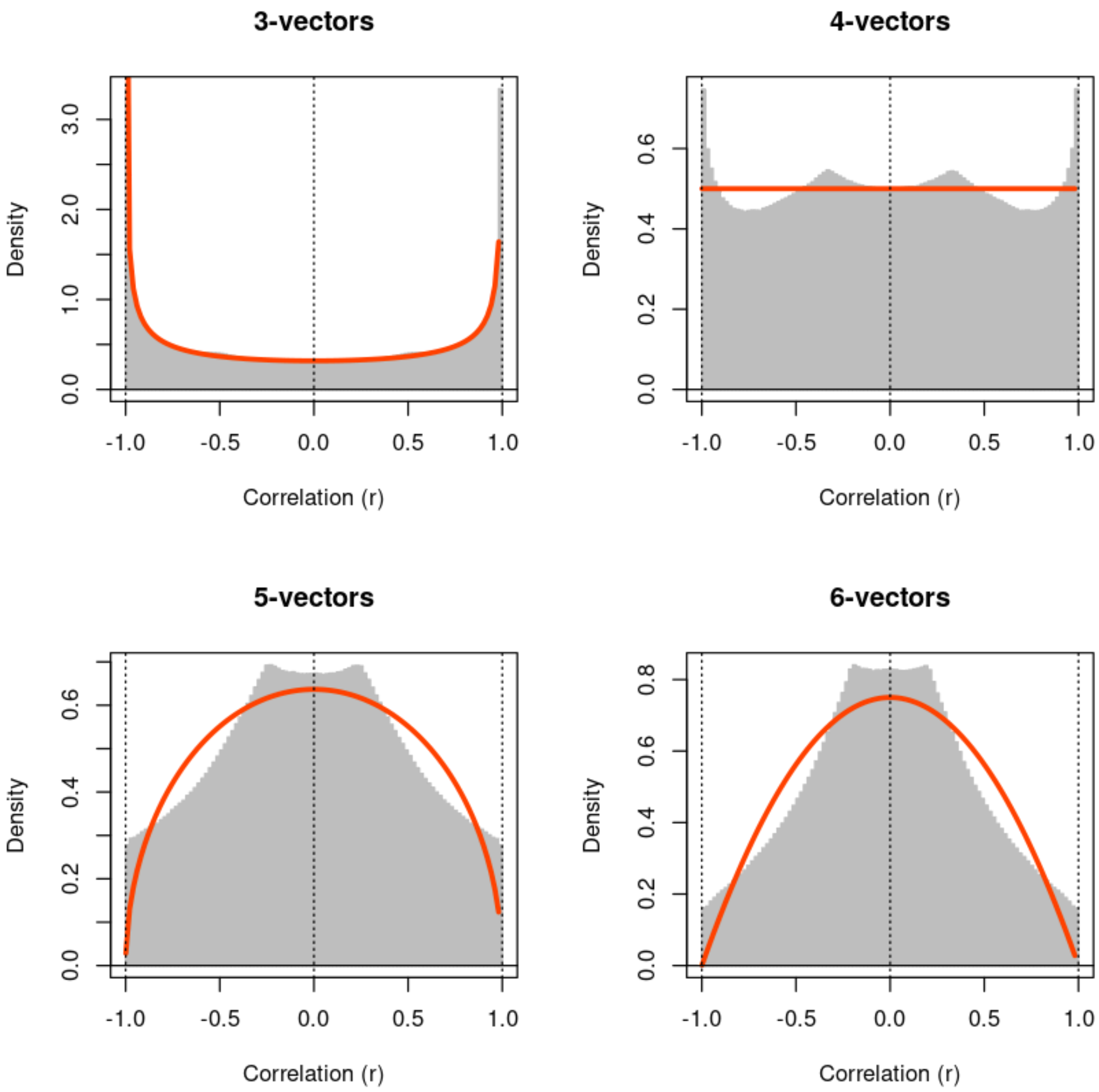
3-vectors
4-vectors
5-vectors
6-vectors
Density
Correlation (r)

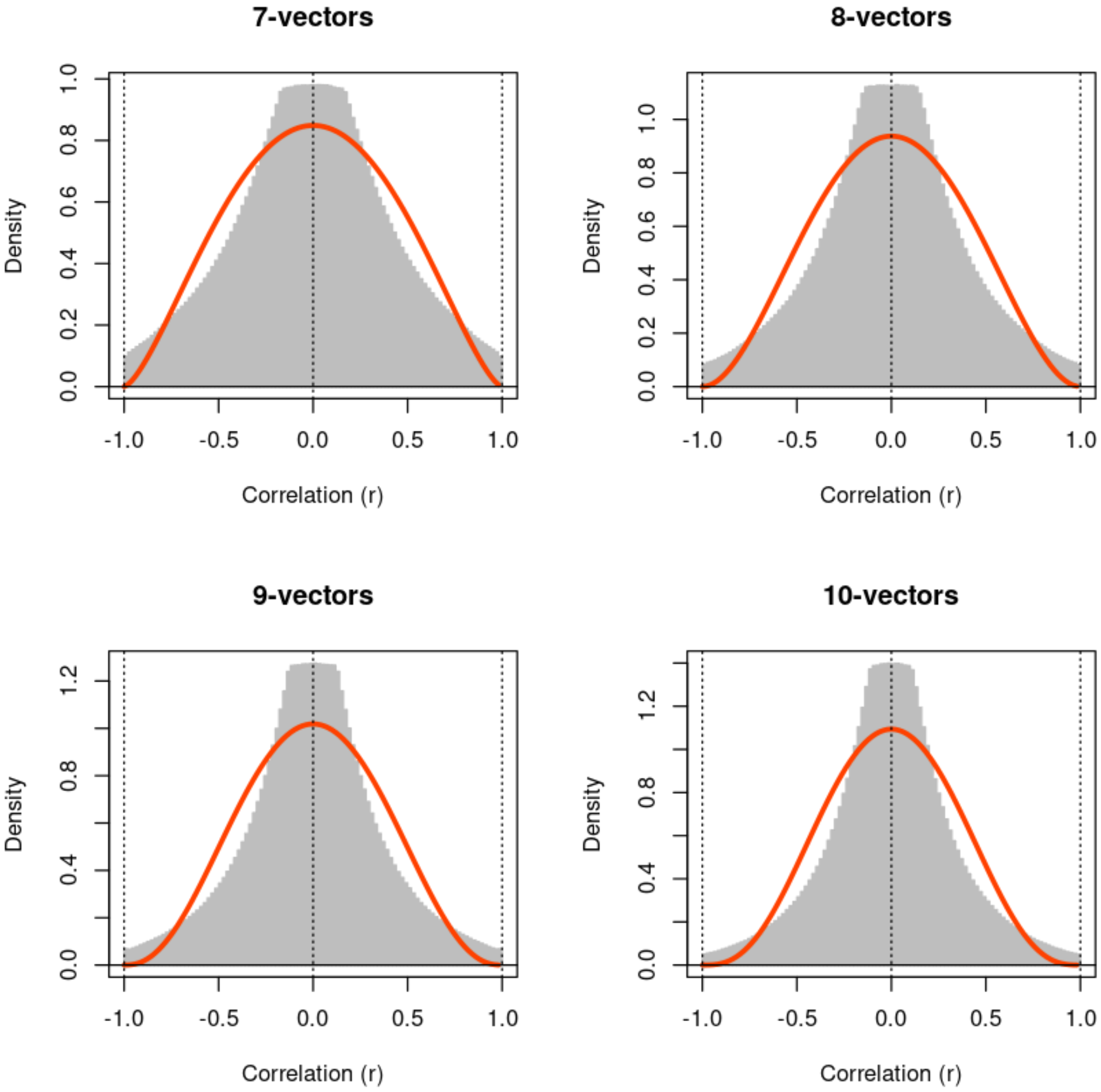
7-vectors
8-vectors
9-vectors
10-vectors
Density
Correlation (r)
-1.0
-0.5
0.0
0.5
1.0
0.2
0.4
0.6
0.8
1.2

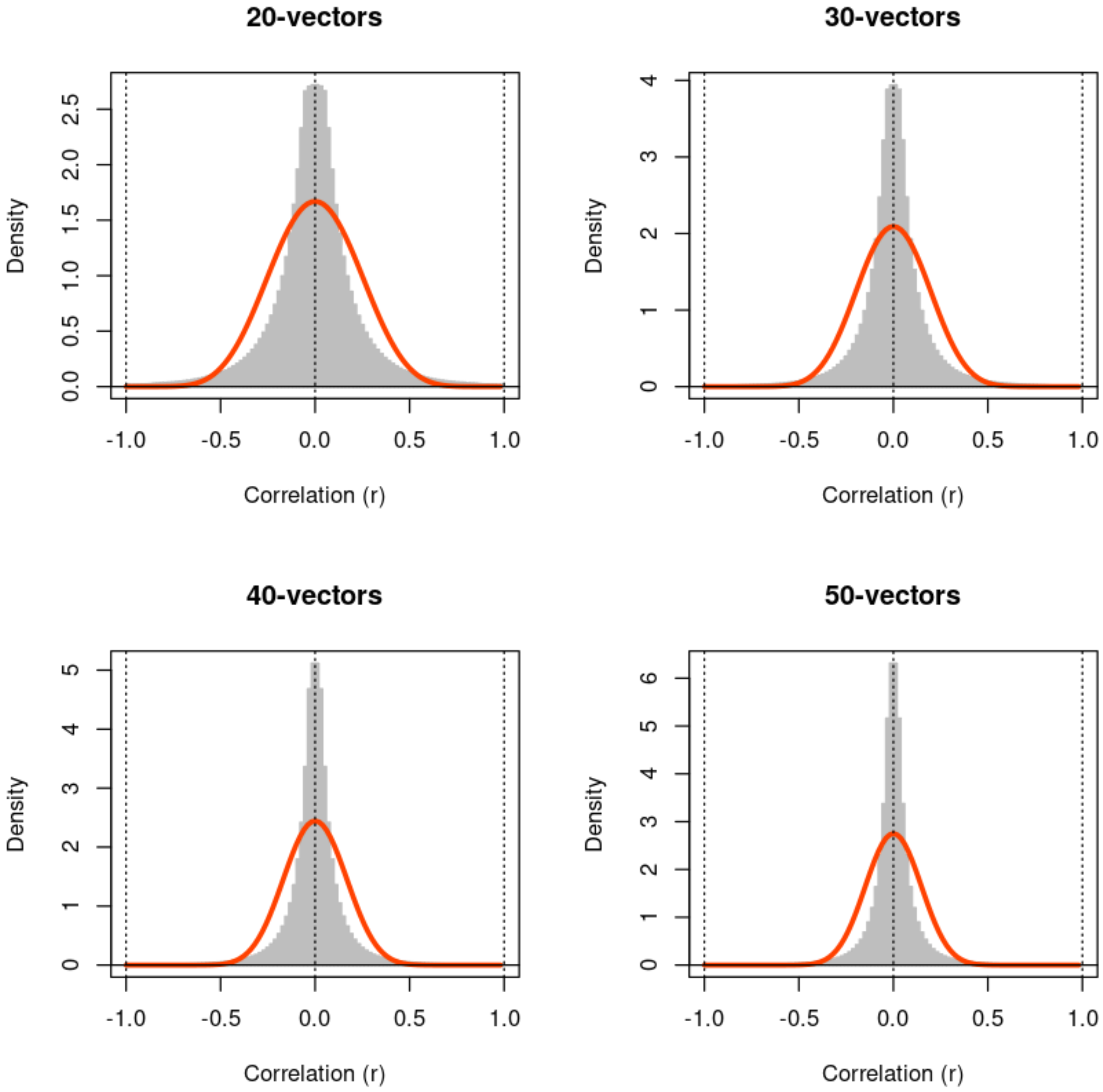
20-vectors
30-vectors
40-vectors
50-vectors
Density
Correlation (r)
-1.0
-0.5
0.0
0.5
1.0

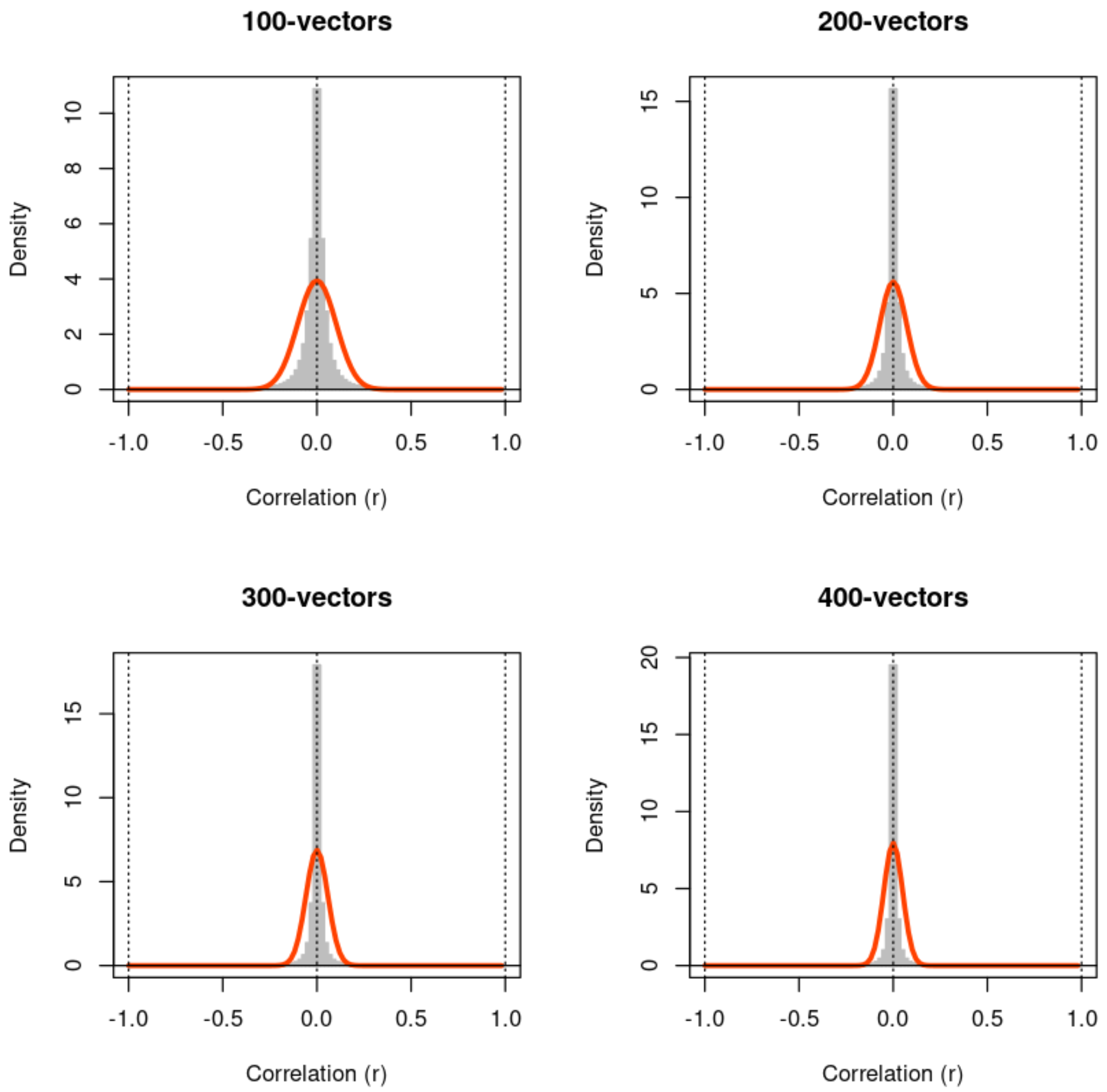
100-vectors
200-vectors
300-vectors
400-vectors
Density
Correlation (r)
-1.0
-0.5
0.0
0.5
1.0

```
In [18]: #end
```
