## Supplement S4 for "Model-based Standardization of Correlation Coefficients Improves Multi-Omic Clustering and Biological Signal Discovery"

**Cholesky Decomposition and Covariance Structure**

A symmetric, *m* x *m* positive definite matrix $\mathbf{\Sigma_m}$ has an unique Cholesky decomposition

$$\mathbf{\Sigma_m = L \cdot L^T},$$

where **L** is a lower triangular matrix and $\mathbf{L^T}$ is its (conjugate) transpose[1]. To produce a matrix **R** with n rows (samples) and m columns (analytes) with designed analyte-analyte correlations, let

| | |
|---|---|
| $\mathbf{\Sigma_m}$ | be a symmetric positive definite *m* x *m* correlation matrix ($\mathbf{\Sigma_m} = \{\rho_{ij}\}$), with diag($\mathbf{\Sigma_m}$) = **1** and $\rho_{ij}$ the designed correlation between columns *i* and *j*, |
| **L =** Chol( $\mathbf{\Sigma_m}$) | be its Cholesky decomposition, and |
| **Z** = $\{z_{ij}\}$ | be an *n* x *m* matrix of standard normal random variables ($z_{ij} \sim \mathbf{N}(0,1)$). |

Then the matrix **R** = **E**[**R**] + **L Z**. will have $\mathbf{\Sigma_m}$ as its covariance matrix[1]:

$$\begin{aligned}\text{cov}(\mathbf{R}, \mathbf{R}) &= \mathrm{E}[(\mathbf{R} - \mathrm{E}(\mathbf{R}))(\mathbf{R} - \mathrm{E}(\mathbf{R}))^T] \\ &= \mathrm{E}[(\mathbf{L \bullet Z})\ (\mathbf{L \bullet Z})^T] \\ &= \mathrm{E}[\mathbf{L(Z \bullet Z}^T)\mathbf{L}^T] \\ &= \mathbf{L} \bullet \mathrm{E}[\mathbf{Z \bullet Z}^T] \bullet \mathbf{L}^T \\ &= \mathbf{L \bullet I \bullet L}^T \\ &= \mathbf{\Sigma_m}.\end{aligned}$$

Our correlation structure is complex, and includes stochastically varying levels of correlation between analytes in the same cluster and additional stochastically varying levels of correlation among analytes from the same ‘omics batch, and designing a positive definite correlation matrix following these constraints would be challenging. Therefore, we used the method above repeatedly in a step-by-step approach (see Methods).

1. Kaiser, Henry F., and Kern Dickman. 1962. “Sample and Population Score Matrices and Sample Correlation Matrices from an Arbitrary Population Correlation Matrix.” *Psychometrika* 27 (2): 179–82.
