## Supplement S5 for "Model-based Standardization of Correlation Coefficients Improves Multi-Omic Clustering and Biological Signal Discovery"

### Beta null models for the correlation types in the Arivale cohort

**Legend (all plots):** Histogram of non-self, non-redundant correlations of the specified type. Green line: Method of Moments estimate.

Blue line: fit by `standardcor::estimateShape()` using default parameters (initial) or using nonzero `left` and `right` parameters (adjusted) chosen by eye to fit the mode and full width at half height.

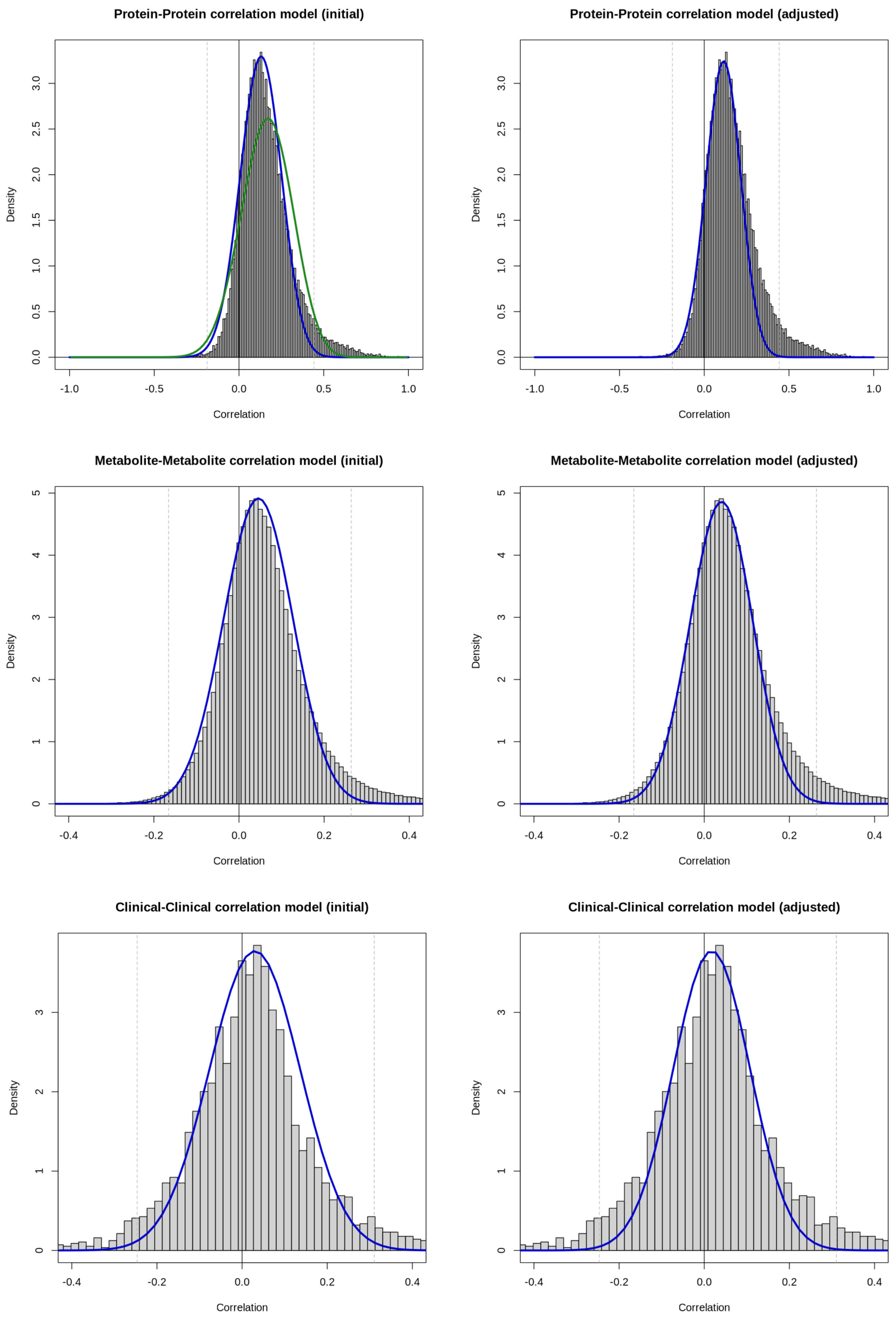

Protein-Protein correlation model (initial)
Protein-Protein correlation model (adjusted)
Metabolite-Metabolite correlation model (initial)
Metabolite-Metabolite correlation model (adjusted)
Clinical-Clinical correlation model (initial)
Clinical-Clinical correlation model (adjusted)
Density
Correlation

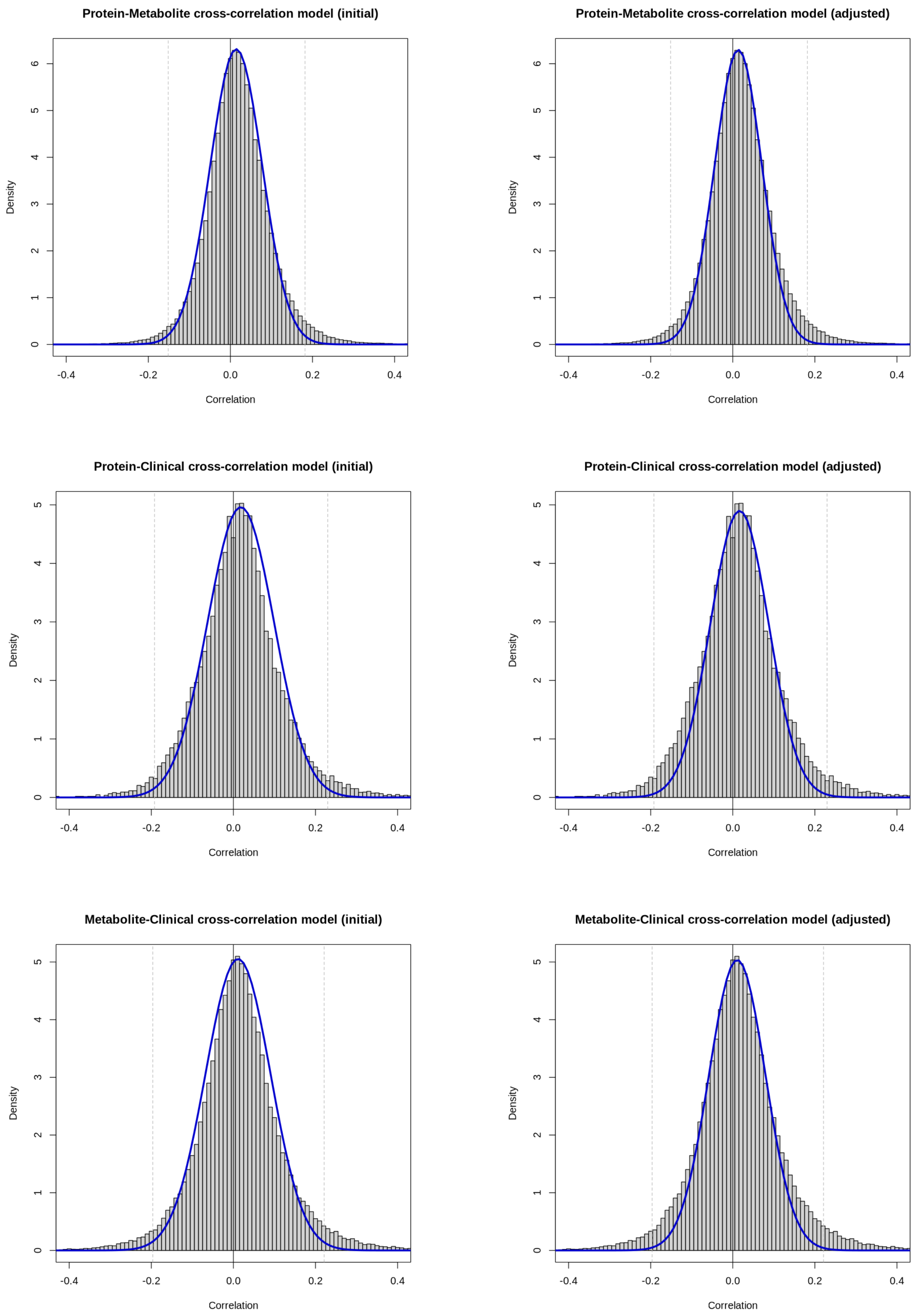
Protein-Metabolite cross-correlation model (initial)
Protein-Metabolite cross-correlation model (adjusted)
Protein-Clinical cross-correlation model (initial)
Protein-Clinical cross-correlation model (adjusted)
Metabolite-Clinical cross-correlation model (initial)
Metabolite-Clinical cross-correlation model (adjusted)
Density
Correlation
-0.4
-0.2
0.0
0.2
0.4
0
1
2
3
4
5
6
